# Ectopic adipogenesis in response to injury and material implantation in an autoimmune mouse model

**DOI:** 10.1101/2023.10.05.561105

**Authors:** Tran B. Ngo, Aditya Josyula, Sabrina DeStefano, Daphna Fertil, Mondreakest Faust, Ravi Lokwani, Kaitlyn Sadtler

**Author notes:** These authors contributed equally (Tran B. Ngo, Aditya Josyula).

## Abstract

Due to the limited capacity of mammals to regenerate complex tissues, researchers have worked to understand the mechanisms of tissue regeneration in organisms that maintain that capacity. One example is the MRL/MpJ mouse strain with unique regenerative capacity in ear pinnae that is absent from other strains, such as the common C57BL/6 strain. The MRL/MpJ mouse has also been associated with an autoimmune phenotype even in the absence of the mutant *Fas* gene described in its parent strain MRL/lpr. Due to these findings, we evaluated the differences between the responses of MRL/MpJ versus C57BL/6 strain in traumatic muscle injury and subsequent material implantation. One salient feature of the MRL/MpJ response to injury was a robust adipogenesis within the muscle. This was associated with a decrease in M2-like polarization in response to biologically derived extracellular matrix scaffolds. In pro-fibrotic materials, such as polyethylene, there were fewer foreign body giant cells in the MRL/MpJ mice. As there are reports of both positive and negative influences of adipose tissue and adipogenesis on wound healing, this model could provide an important lens to investigate the interplay between stem cells, adipose tissue, and immune responses in trauma and materials implantation.

## INTRODUCTION

Tissue regeneration is a complex biological process to repair and renew injured tissue by replacing it with functional tissue. This restoration can be either partial, resulting in the recovery of a portion of the original structure and function, or complete, reflecting the process seen during development. However, complete tissue regeneration, such as limb regeneration in salamanders, is rare among mammals, with notable instances found in the regrowth of fingertips in children and digital tips in fetal mice [1]. These regenerative processes heavily depend on forming a specialized structure called a blastema, particularly associated with the keratinized ectodermal appendage found exclusively at the tips of digits [2]. In addition to digit regeneration, the Hedgehog signaling pathway, a well-studied regulator of embryonic development, is also critical in controlling the fate of fibro/adipogenic progenitors during skeletal muscle regeneration in adult mice [3].

Among mouse species, complete regeneration is observed in response to whole-thickness skin injuries in the African spiny mice [4]. Another well-studied model in the context of healing and regeneration is the Murphy’s Roth’s Large mouse strain (MRL/MpJ or MRL), also being investigated for its autoimmune characteristics [5]. The MRL/MpJ mouse strain, which has an altered response to injuries, can offer insights into the barriers hindering human scar-free healing. The MRL mouse was bred from C57BL/6, LG, AKR, and C3H backgrounds, with most of its genome coming from the LG (Large) mouse (Supplementary Figure 1). It is worth noting that diverse laboratory mouse strains exhibit distinct wound-healing capacities and varying tendencies toward immune system activation. Therefore, direct comparisons with routinely used laboratory mouse strains such as C57BL/6 are warranted.

The MRL mouse has an interesting capacity to fully regenerate ear punches compared to a C57BL/6 (B6) mouse, which results in scarring [6]. This healing capacity does appear limited as other studies evaluating myocardial infarction and full-thickness skin wounds still result in scarring in the MRL mice [7-9]. During ear punch wound healing, the MRL mouse develops a blastema-like structure and a layer of white adipose on the wound edge, which ultimately closes without extensive collagen deposition [9, 10]. White fat is an immunologically active organ, and fat pads, such as the omentum, have been shown to migrate to the site of injury in humans [11]. Additionally, type-2 immune signals have been associated with wound healing, white fat homeostasis, and metabolism [12]. Regarding metabolism, the MRL/MpJ mouse strain has been reported as the first naturally occurring mouse strain resistant to high-fat diet-induced metabolic changes [13]. For example, the MRL adults and neonatal mice share characteristic transcription patterns, such as the up-regulation of genes engaged in PPAR signaling and retinol metabolism and the repression of immune response genes [14]. These metabolic differences may contribute to the MRL/MpJ mouse strain’s unique immune profile and “super-healing” characteristics.

There are many immunological factors that have been implicated in metabolic-related pathways, such as white fat homeostasis, including interleukin-33 (IL-33) and type-2 innate lymphoid cells (ILC2s) that express ST2 (the receptor for IL33), IL-5 and eosinophils, as well as IL-4 and IL-10 along with M2 macrophages and Th2/regulatory T cells [15-17], which are important regulators for adipocyte proliferation, differentiation, and adipokine release. MRL mice with a Fas mutation (MRL/lpr) have shown resistance to obesity triggered by a high-fat diet. This resistance is attributed to elevated levels of IL-4 and IL-10; this effect could be linked to STAT6 activation, potentially promoting M2 polarization by enhancing IL-4 and IL-10 levels [18]. In wound healing, type-2 immune signals regulate collagen deposition to close wounds and assist in the proliferation and differentiation of progenitor cells. These signals are important in fibro-adipogenic lineage commitment and can directly affect the amount of fat, muscle, or scar tissue deposited during recovery from physical trauma [3].

Inflammatory processes associated with wound healing that result in collagen deposition occur after biomedical material implantation. Decellularized extracellular matrix (ECM) is used clinically in various applications, including dural repair, hernia repair, skin wounds, and diabetic ulcers. In this study, ECM was used as the pro-regenerative material applied to the wound site of the murine volumetric muscle loss model (VML) [19]. ECM is being tested with success in larger volume defects such as muscle trauma, though research is still ongoing [20-22]. On the other hand, polyethylene (PE) is a hydrophobic polymer used widely in medical device implants due to its durable and moldable nature; however, when wear particles are generated and embedded into surrounding tissue, this leads to excessive inflammation and scar tissue deposition [23-25]. Utilizing a murine volumetric muscle loss model, we also applied PE as the pro-inflammatory/pro-fibrotic material to the wound site [19]. In this work, we sought to evaluate the differential immunologic and resulting tissue structure outcomes in trauma and different material implantation in B6 versus MRL mice and determine if the white fat accumulation during ear punch healing was also prevalent during response to biomaterial implants.

## MATERIALS & METHODS

### Extracellular matrix decellularization

Small intestine (SI) was isolated from Yorkshire Pigs (Wagner Meats). Fecal matter was removed with tap water, and the mesentery and muscular layers were removed before portioning into 12- inch sections. SI was then opened using a razor blade, and the inner mucosal layer was removed via manual debridement. The resulting submucosa (SIS) was rinsed thoroughly in distilled water and then sterile distilled water with 1% antibiotic antimycotic solution (Gibco). Excess was stored at -80°C until decellularization. Antibiotic-treated SIS was rinsed thoroughly in distilled water and then incubated in 4% ethanol and 0.1% peracetic acid (Sigma) for 30 minutes with vigorous shaking. The resulting decellularized extracellular matrix (ECM) was rinsed in successive washes of sterile distilled water followed by sterile 1xPBS (Fisher) until pH returned to neutral. Samples were washed one final time with sterile distilled water before being drained and frozen overnight at -80°C. The ECM was then lyophilized and then milled into a fine powder using a SPEX SamplePrep Cryogenic milling device.

### Volumetric Muscle Loss Surgery

Surgery and material implantation occurred as previously described [21, 26-28]. The day before surgery, 6- to 8-week-old female C57BL/6J mice (Jackson Labs) were anesthetized under isoflurane, and hair was removed from their hindlimbs up to the xiphoid process using an electric razor followed by depilatory cream. On the day of surgery, mice were anesthetized and received 1mg/kg buprenorphine SR (ZooPharm) and eye lubrication before sterilizing the surgical site with three rounds of betadine followed by 70% isopropyl alcohol. An incision of 1 cm was made in the skin overlying the quadriceps muscle group, and a 3 mm x 3 mm section of muscle was surgically removed from the mid-belly of the quadriceps muscle group using scissors. The resulting tissue gap was treated with either vehicle control (saline), 50 µL extracellular matrix powder (ECM, hydrated to 3 mg/ml with saline), or 50 µL high-density polyethylene (PE) powder (Goodfellow Scientific). ECM was prepared as noted above; PE was washed with sterile distilled water followed by 70% ethanol and placed under UV light for 30 minutes. The resulting suspension was dried overnight in a biosafety cabinet. The NIH Clinical Center Animal Care and Use Committee approved and supervised all procedures under protocols NIBIB20-01 and NIBIB23-01.

### Histology

Quadriceps muscles were removed from the bone and fixed in 10% neutral buffered formalin (Sigma) for two days, rinsed in distilled water and dehydrated in a graded ethanol series before clearing in xylenes and infiltrating with paraffin wax (Leica). The resulting formalin fixed paraffin embedded (FFPE) samples were mounted on paraffin blocks and sectioned into 5 µm sections then stained with hematoxylin and eosin (H&E, Sigma) or picrosirius red (PSR, Abcam) using standard protocols as per manufacturer’s instructions. Slides were imaged on an EVOS microscope.

### Immunohistochemistry

FFPE-sectioned slides underwent rehydration to prepare samples for immunofluorescent staining and were subsequently incubated for 20 minutes in a citrate antigen retrieval buffer in a vegetable steamer. Following this, a gradual cooling process for 20 minutes occurred on the benchtop. The activity of endogenous peroxidases were quenched by a 5-minute treatment in a solution of 0.3% hydrogen peroxide within 1xPBS. For staining, the VECTASTAIN® Elite ABC-HRP Kit was employed following the guidelines provided by the manufacturer (Vector Laboratories). Initially, after a wash in 1xPBS, the samples were blocked in 2.5% normal goat serum for one hour. Rabbit monoclonal anti-CD11b (Abcam) and anti-CD206 (Cell Signaling), were diluted at a 1:200 ratio in blocking buffer and incubated for one hour at room temperature. After three washes in 1xPBS, biotinylated secondary antibodies were introduced for a 30-minute incubation. Then, a 30-minute incubation with the VECTASTAIN Elite ABC Reagent was followed by three additional 1xPBS washes. ImmPACT® DAB EqV Peroxidase (HRP) Substrate (Vector Laboratories) was applied for either 1 minute and 30 seconds. Slides were rinsed in tap water, subjected to a 5-minute counterstaining with Harris Hematoxylin (Sigma), rinsed again, de-stained with acid ethanol, dehydrated, and mounted with Permount (Fisher Scientific).

### Flow cytometry

Quadriceps muscles were removed and diced finely using surgical scissors before digestion in 10 ml of 0.25 mg/ml Liberase TM (Millipore Sigma) with 0.2 mg/ml DNAse I (Millipore Sigma) for 45 minutes in serum-free DMEM (ThermoFisher Scientific), shaking at 37°C. The resultant material was mechanically pressed through a 70 µm cell strainer that was then washed with 1xPBS + 5 mM EDTA to a volume of 50 ml. The suspension was centrifuged at 350 xg for 5 minutes and the cell pellet was washed once more in 1xPBS + EDTA before soaking in PBS + EDTA on ice for 10 minutes to dissociate further. The samples were transferred to a 96-well plate, and cells were counted via a hemocytometer using Trypan Blue (ThermoFisher) before staining in LIVE/DEAD Blue Fixable viability dye (ThermoFisher Scientific). Cells were stained with a surface antibody cocktail for both myeloid and lymphoid linage panels as previously reported protocols [28, 29]. As a note, for the lymphoid lineage panel, CD49b-BV421 (Biolegend) was substituted for CD49b- V450 antibody compared to the previous protocol, as the latter was discontinued by the manufacturer. Samples were run and spectrally unmixed on a 5-laser Cytek Aurora Flow cytometer (Cytek Biosciences) through the SpectroFlo software, and data was analyzed on FlowJo (BD Biosciences) and GraphPad Prism (for Gating Strategy and example plots see Supplementary Figure 2, 3).

### Bulk RNA Sequencing Data Analysis

Whole quadriceps muscles were flash-frozen on dry ice for bulk RNA isolation and sequencing through Azenta GeneWiz (Supplementary Table 1). Bulk RNAseq analysis was performed on the paired-end reads from 24 independent samples. FastQC was used to determine basecall accuracy, and the quality score was verified to be > 30 for each sample. Pseudoaligment was performed using Kallisto to map reads to a publicly available reference transcriptome (Ensemble GRCm39). Transcript abundance files were created across all 24 samples, and data was imported into R (Version 4.2.1, Funny-Looking Kid) for downstream analyses. The Ensemble Mus musculus v79 annotation database mapped transcripts to gene IDs. EdgeR package was used to determine counts per million and subsequently filtered by counts per million > 1 for at least 3 samples (smallest group of comparison method). Counts were then log2 transformed. Data was normalized using the Trimmed Mean of M-values method. Differential expression was visualized using standard functions to generate heatmaps and volcano plots for each pairwise comparison. Differentially expressed genes were confirmed with an independent analysis through Azenta Genewiz. Gene Ontology enrichment was performed using the gprofiler2 package on select differentially expressed genes. The C2 subcollection of genes was read into R from the molecular signatures database. Gene Set Enrichment Analysis was carried out using the clusterprofiler package and visualized through bubble plots and running enrichment score profiles using the GSVA package (Supplementary Figure 4). The code used for analyzing bulk RNA sequencing data is available in **Supplementary Methods.**

### Ternary plots and protein-protein interaction

To display ternary plots, we compiled a list of genes that comprised the “Coates – Macrophage M1 vs M2 up” gene set. The code used is available in **Supplementary Methods**. We then rescaled the average log2(CPM) data (0-100) using the ggtern package, such that for each gene, the highest count across the three treatment groups was set to a score of 100. Combining interaction scores for leading-edge genes obtained from enriched gene sets in GSEA were text mined from the String database to map protein-protein interactions. The circlize package was then used to create chord diagrams with links and sector sizes representing individual interactions and the extent of interaction, respectively.

### Rheological measurements

Strain sweep experiments were conducted using a Kinexus Pro+ Rheometer (NETZSCH) with a temperature control system. Tissue samples were carefully examined at a temperature of 25°C, positioned on a lower plate with a diameter of 60 mm with an outer ring for aqueous solution trap and an upper plate with a diameter of 20 mm. The quadriceps muscle samples were collected from healthy animals and cut into a longitudinal section of approximately 5 mm in length, 4 mm in height and 4 mm in width. The tissue was gently positioned between the fixed bottom part of the rheometer and an oscillating top plate, ensuring that the muscle fibers were aligned in parallel with the plates. An initial normal load force of 0.25 N was applied to the sample. The strain sweep tests were then performed within a shear strain range of 0.01 – 2%, maintaining an oscillation frequency of 1 rad/s. For data acquisition, 8 data points per decade were collected per sample as previously described [30]. Under the described strain sweep test setting, the measurement of B6 muscle samples were taken with a gap of 2.139 – 2.641 mm and the MRL muscle samples were compressed with a gap of 2.611 – 3.536 mm (n = 5).

## RESULTS

### Immunologic differences in response to muscle injury in C57BL/6 and MRL/MpJ strain

To model traumatic injury, we employed the VML model, creating an injury site in the mouse’s quadriceps muscle group, followed by treatments of the saline vehicle as a control, implantation of a pro-regenerative (decellularized extracellular matrix, ECMtx), or pro-fibrotic (polyethylene, PEtx) material. Muscle tissue morphology was evaluated at 7-, 21-, and 42-days post-injury via histopathology, and the tissue response and immune microenvironment to implanted materials were evaluated at 21-days post-injury with flow cytometry and bulk RNA sequencing (Figure 1a, Supplementary Figure 2-5). Even without injury, we found differences in gene expression in quadriceps muscles of B6 and MRL mice (**Figure 1b**). Several of these differences were pseudogenes and unknown transcripts. The patterning of these differences shifted but was still significantly present, with new differences in transcriptomes appearing after a VML control injury treated with a saline vehicle control (**Figure 1c, Supplementary Figure 6**). When focusing on the standard C57BL/6 model, we could isolate strongly changed genes with known protein-coding transcripts modulated with injury (**Figure 1d**). These included a robust increase in *Ly86* (associated with TLR4 signaling and response to bacteria), *Ccl8* (encoding MCP-2, a chemokine receptor for mast cells, eosinophils, and basophils), *Cd74* (involved in MHCII antigen presentation), several complement genes (*C1qb*, *C1qa*, *C3ar1*), muscle-associated *Myh8*, *Col19a1* (an uncharacterized, but likely fibril-forming collagen), and *Itgax* (encoding CD11c, which is expressed on dendritic cells and macrophages), among others. Several genes were significantly lower (p-adj < 0.05) in injured tissues, including Acsm3 (involved in acyl-CoA metabolic processes, suggesting differences in metabolism), *Cnn1* (inhibitor of vascular smooth muscle cell proliferation), and *Plin5* (white fat), among others.

**Figure 1.**
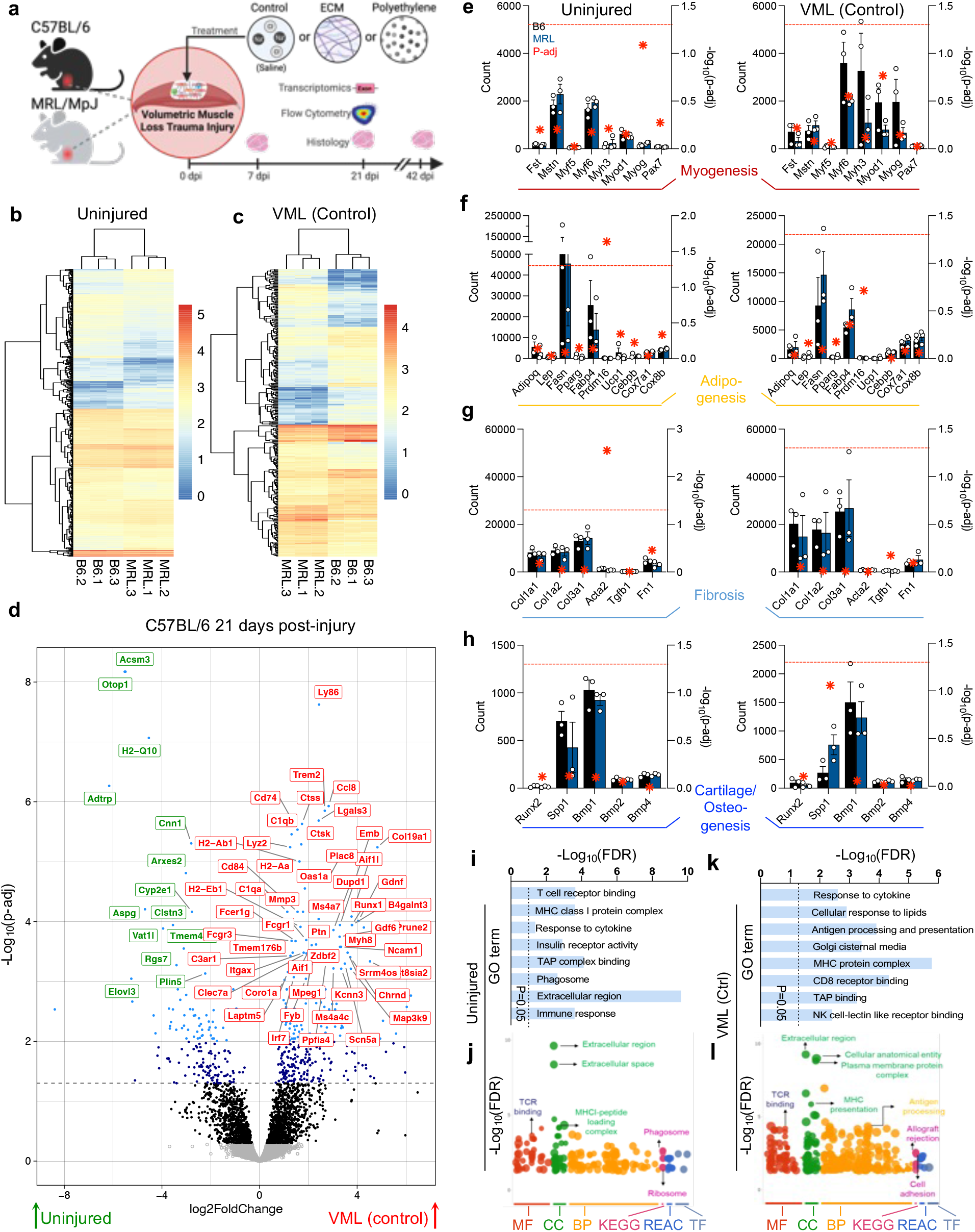
Baseline and injury-associated differences in the skeletal muscle transcriptome of C57BL/6 and autoimmune MRL/MpJ mice. (**a**) Schematic of study design. (**b - c**) Transcriptome differences in (**b**) uninjured and (**c**) volumetric muscle loss (VML) control injury mouse quadriceps muscles, data are log-normalized counts. (**d**) Significantly altered genes in C57BL/6 mice after VML injury. Red = upregulated in injury, Green = downregulated in injury, dashed lined = adjusted p-value (p-adj) 0.05. (**e-h**) Gene counts of selected (**e**) myogenesis, (**f**) adipogenesis, (**g**) fibrosis, (**h**) cartilage/osteogenesis associated genes. Gene counts on the left y-axis, Black = C57BL/6, Blue = MRL/MpJ, red stars = - log10p-adj (right y-axis), dashed line = adjusted p-value 0.05. Data are means ± SEM, n = 3. (**i**) Gene ontology (GO) enrichment in MRL uninjured mice versus C57BL/6. (**j**) GO and pathway enrichment in uninjured MRL mice versus C57BL/6. (**k**) Gene ontology enrichment in MRL control injured mice versus C57BL/6. (**l**) GO and pathway enrichment in control injured MRL mice versus C57BL/6. MF = Molecular function, CC = Cellular compartment, BP = biological process, KEGG = Kyoto Encyclopedia of Genes and Genomes, REAC = Reactome, TF = Transfac database. The Red dashed line indicates p-adj=0.05 in (**e – h**) plotted on right y-axis.

Focusing on genes linked with different tissue development pathways, we saw general increases in myogenesis-associated genes after injury in both B6 and MRL mice (**Figure 1e**), with decreases in *Mstn* encoding the inhibitory protein myostatin. In uninjured mice, the MRL/MpJ strain had significantly higher (P-adj < 0.05) transcript levels of *Prdm16* associated with brown fat, with *Fasn* and *Fabp4* (white-fat associated) trending lower in MRL mice pre-injury (in comparison to uninjured B6 group) and lower in B6 mice (in comparison to MRL group) post-injury, albeit not significantly due to the variability in transcript counts (**Figure 1f**). There was a general increase in fibrillar collagens associated with scar tissue deposition after injury (**Figure 1g**), with MRL mice having lower *Acta2* (encoding alpha-smooth muscle actin, a myofibroblast marker) than B6 mice before injury. There were fewer differences in cartilage development and osteogenesis markers after injury except in B6 mice, where an increase in *Spp1* was observed (**Figure 1h**). When comparing gene ontology enrichment in uninjured (**Figure 1i, j**) and VML control injury (**Figure 1k, l**) in B6 and MRL mice, several patterns emerged. MRL mice enriched GO-associated pathways strongly in the extracellular region, T cell receptor binding, and the MHCI protein complex (FDR < 0.05, **Figure 1i**). Regarding KEGG pathway enrichment, there were increases in the phagosome and ribosome pathways (**Figure 1j**). After injury, the MHC protein complex was still enriched with antigen processing, presentation, and CD8 receptor binding (**Figure 1k**). There was also a significant increase in allograft rejection pathways in MRL mice (FDR < 0.05, **Figure 1l**), correlating with the known propensity of MRL mice to autoimmune phenotypes.

### Injury and biomaterial implantation result in white adipogenesis in MRL/MpJ strain

During injury recovery, several differences between MRL and B6 mice were observed via histopathology (**Figure 2a-b**). Early on, at 7 days post-injury, the pathologic evaluation looked similar between B6 and MRL with both ECM (**Figure 2a**) and PE (**Figure 2b**) treatments. While some adipogenesis is observed in B6 mice in agreement with prior studies [26], there was a robust increase in white fat in MRL mice by 21 days post-injury. This adipogenesis was present around materials but also within the injured skeletal muscle. While fat pad migration into an injury site can occur, the morphology of this adipogenesis suggested de-novo white fat formation, given its integration with muscle fibers. There was some evidence for a slight increase in hemosiderin deposition (brown color in hematoxylin and eosin-stained section) associated with differences in debris clearance and an inability to phagocytose hemoglobin and red blood cell debris properly.

**Figure 2.**
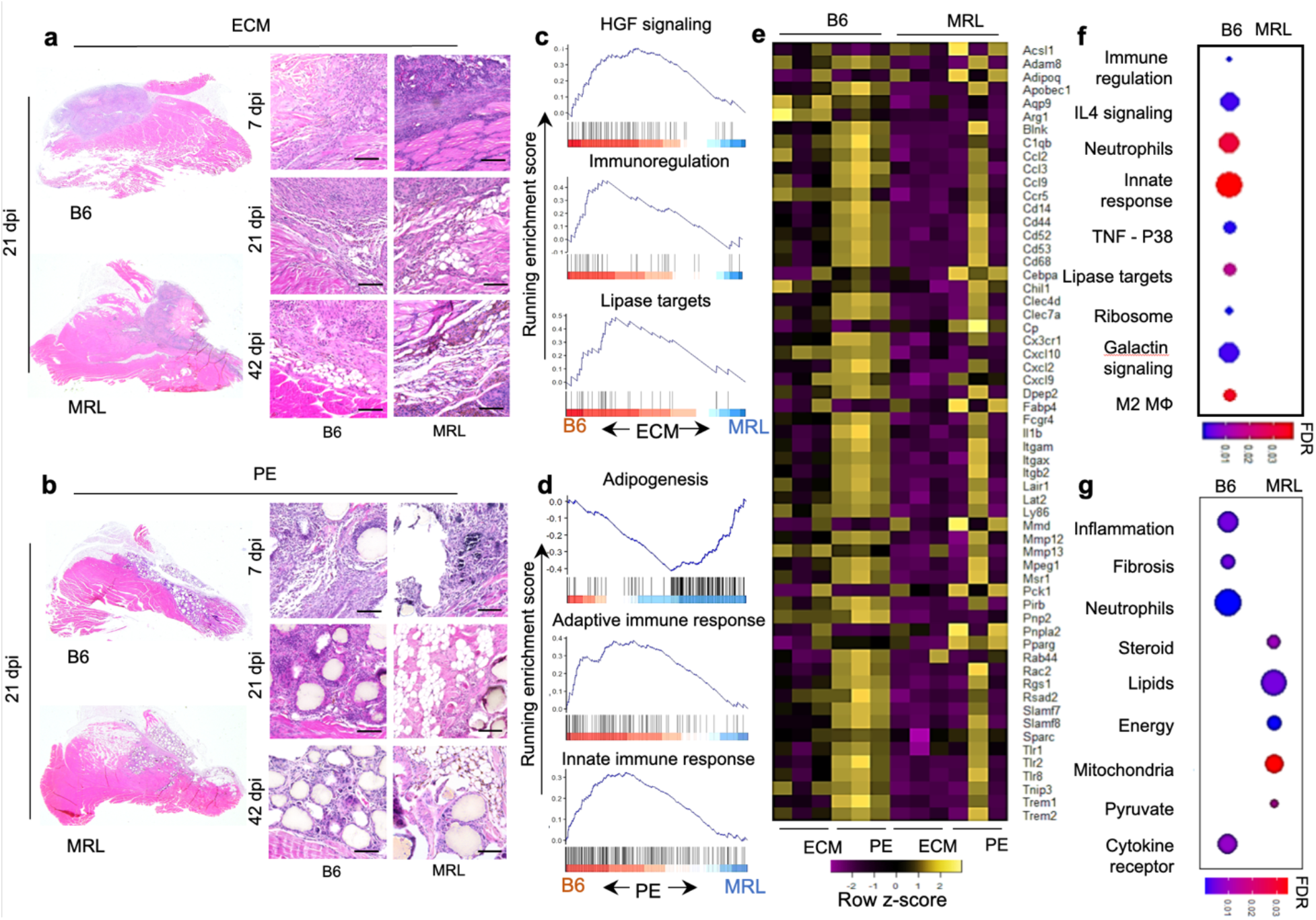
Hypertrophic intramuscular adipogenesis with divergent immune responses in MRL mice after volumetric muscle loss injury and biomaterial implantation compared to B6 mice. Hematoxylin and eosin (H&E) images of muscle at 7 days, 21 days, and 42 days post-injury (dpi) of B6 and MRL mice with (**a**) decellularized extracellular matrix (ECM) treatment (**b**) with polyethylene (PE) particle treatment, scale bars = 100 µm, representative of n = 5 mice. GSEA representation of select enriched adipogenic, lipid metabolic, and immune response pathways (FDR<0.05) in differentially expressed genes in B6 and MRL mice 21 dpi in response to (**b**) ECM and (**e**) PE material implantation. (**c, f**) Running enrichment scores for gene sets associated with metabolic and immune response pathways in mice that received ECM and PE implants, respectively. (**g**) Heatmap of leading-edge genes obtained from enriched gene sets. Color gradient scale bars in **f** and **g** represent false discovery rate (q-value), n = 3.

In ECMtx wounds, B6 mice had a stronger HGF signaling response (inducing angiogenesis and organogenesis) and immunoregulatory response (FDR < 0.001) (**Figure 2c**). Correlated with the outcomes in histology, where we saw increased white fat in MRL mice, we observed increased lipase target genes (Lian_Lipa_3M gene set; FDR < 0.001) in B6 mice treated with ECM. Next, we performed gene set enrichment analysis on injured muscle tissue that received ECM and PE materials. Comparing PEtx mice, we found an increase in adipogenesis genes and an overall enrichment in multiple adipogenesis gene sets (Burton-Adipogenesis; FDR < 0.001,) in MRL mice supporting histopathology of the wound area (**Figure 2d**). Interestingly, in the B6 background, PEtx enriched, the gene sets “Reactome-Adaptive immune system” and “Reactome-Innate immune system” at 3 weeks post-injury (FDR < 0.001). In ECMtx mice, we saw enrichment of multiple immune-related gene sets in B6 mice, including IL-4 signaling (FDR<0.001) (associated with M2 macrophages and a Th2 polarized immune response), neutrophil degranulation (FDR < 0.001) (associated with early granulocytic infiltration to degrade debris and fight infection), innate response (which includes granulocytes as well as macrophages), and M2 macrophages (which correlates with the IL-4 signaling enrichment, **Figure 2f**). Furthermore, PEtx enriched several metabolic function gene sets, including lipid, steroid, and pyruvate metabolism and mitochondrial function; a heatmap of select markers that were differentially expressed in PEtx compared to ECMtx in both mouse backgrounds is presented in Supplementary Figure 7.

### Enrichment of innate immune responses after injury depends on the mouse strain

Overall, myeloid infiltration into the wound microenvironment was characterized through immunostaining for CD11b (**Figure 3a**). CD11b+ myeloid cells were detected within the fat pad against the injury site in B6 and MRL control injuries. In ECMtx mice, these cells were largely infiltrating the material with a preferential presence towards the material’s more distal surface as opposed to the relatively fewer CD11b+ myeloid cells at the injury interface. MRL mice appeared to have a denser cellular infiltration than the B6 mice. With PEtx, a dense CD11b staining compounded with granular nuclei resembling neutrophils was found surrounding each PE particle, along with staining between particles within more fibrous areas with lower nucleic density.

**Figure 3.**
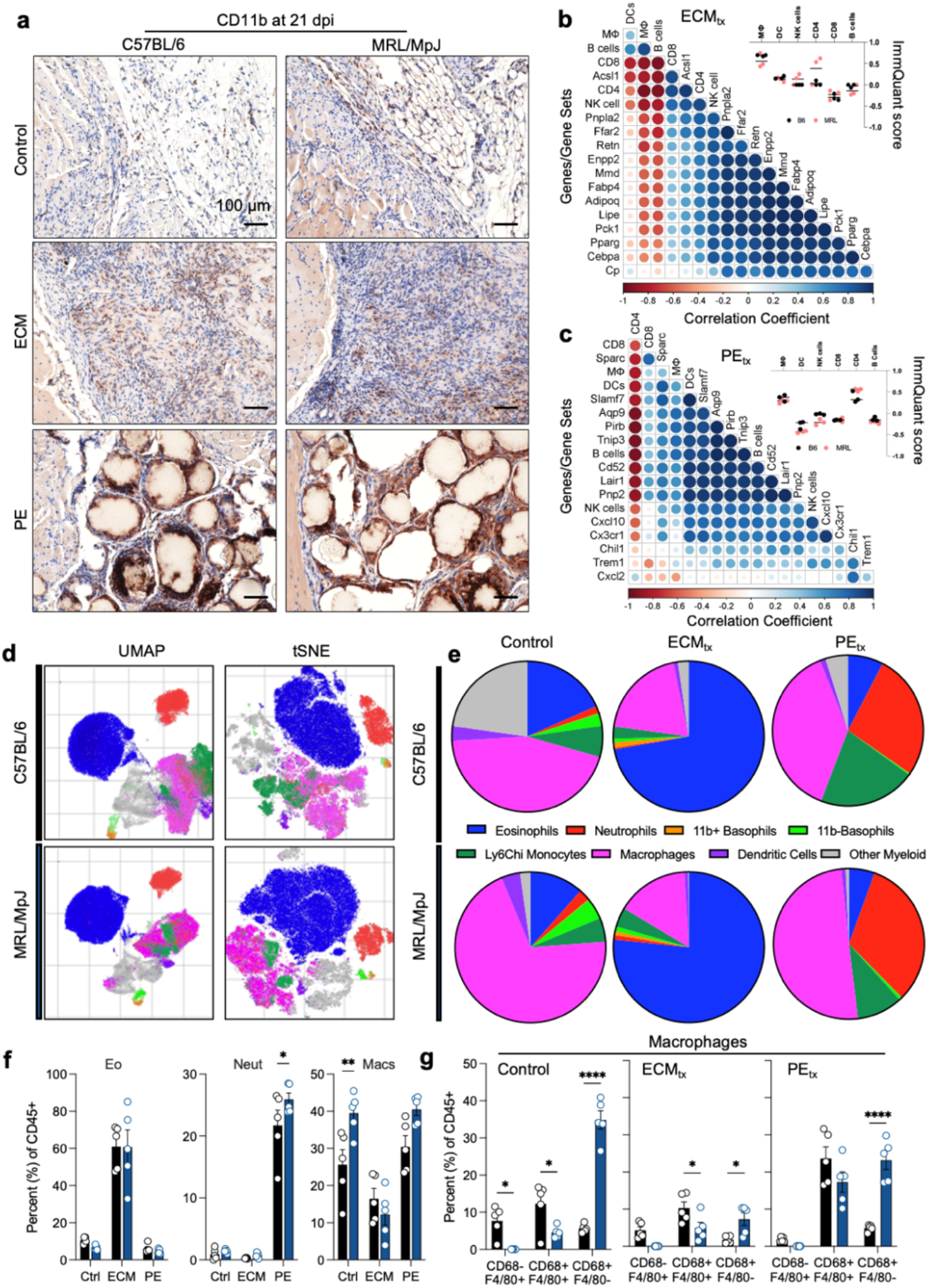
Immune response to biomaterial implants in a traumatic muscle injury. (**a**) CD11b immunohistochemistry at 21 days post-injury (DAB = brown, Hematoxylin = purple). (**b-c**) Correlation plots of (**b**) ECMtx and (**c**) PEtx muscle injuries. (**d**) UMAP and tSNE projections of myeloid infiltrate determined via flow cytometry at 21 days post-injury. (**e**) Myeloid cell populations in C57BL/6 (top row) and MRL/MpJ (bottom row) mice. (**f**) Top three myeloid cell populations in B6 and MRL mice. Eo = eosinophils, Neut = neutrophils, Macs = macrophages. (**g**) Sub-populations of macrophages as determined by CD68 and F4/80 expression. Data are means ± SEM, n = 5 (FACS, IHC) or 3 (RNAseq). ANOVA with Tukey post-hoc correction for multiple comparisons; * = P < 0.05, ** = P < 0.01, **** = P < 0.0001.

### Adipogenic gene expression negatively correlates with F4/80+ MHCII- expressing macrophage gene signature

Next, we wanted to understand which immune cell subsets were correlated with adipogenic gene expression. We compiled a list of over 14,000 genes and created a gene expression matrix using log-transformed counts per millions of transcripts across all samples. We input this matrix into ImmQuant [31] to generate ImmQuant scores (signifying the extent of enrichment of immune cell types) of various immune cell subsets based on gene expression data in the ImmGen database. Having previously seen adipogenic gene sets enriched in PEtx by GSEA, we created a matrix of ImmQuant scores and adipogenic gene expression data within PEtx mice. We then calculated correlation coefficients for select adipogenic marker genes and ImmQuant scores of immune cell subsets of interest. First, we found that a macrophage subset expressing F4/80 but not CD68 or MHC II had an ImmQuant score greater than 0.2 in all samples (**Figure 3b** inset). Further, this subset negatively correlated with adipogenic gene expression (including *Adipoq*, *Cebpa*, *Lipe*, and *Pparg*) measured in bulk RNA sequencing analysis (**Figure 3b**). Second, a CD4+ T cell signature with ImmQuant scores greater than 0.2 in most MRL samples (**Figure 3c** inset) positively correlated with an adipogenic gene expression (**Figure 3b**). Additionally, we performed a similar analysis examining the same immune cell signatures and leading-edge genes in the HGF signaling gene set in mice that received ECMtx. Interestingly, in contrast to adipogenic gene expression, we found that the macrophage subset signature correlated positively, and the CD4+ T cell signature correlated negatively with the expression of *Pirb*, *Lair1*, *Pnp2*, *Cxcl10,* and *Cx3cr1* (**Figure 3c**). Further, we text-mined the string database (string-db.org; [32]) for interaction scores of various proteins encoded by leading-edge genes in the enriched HGF signaling and adipogenesis gene sets, along with key innate and adaptive immune markers. Visualizing this data through chord diagrams (Supplementary Figure 8), we found that the macrophage marker CD68 linked to the adipogenic proteins *Adipoq* and *Pparg*. Among the proteins involved in HGF signaling, the most notable interactions were *Itgb2* with the lipase target *C1qb* and *Msr1*-*Csf2ra*

Through a 22-color flow cytometry panel, we could further analyze the myeloid compartment and identify several myeloid cell types and subtypes (**Figure 3d**). These included eosinophils (SiglecF+), neutrophils (Ly6G+), CD11b+ basophils (CD200R3+), CD11b- basophils, Ly6C^hi^ monocytes/macrophages (F4/80+Ly6c^hi^), macrophages (F4/80+ and/or CD68+), dendritic cells (CD11c+), and other myeloid cells (CD45+CD11b+Lin-) that varied with treatment type and mouse strain (**Figure 3e**). ECMtx robustly increased eosinophils compared to control and PEtx injury in B6 and MRL mice (**Figure 3f**). Conversely, PEtx recruited a strong neutrophil presence, as noted in histopathology. MRL mice had a slightly increased presence of neutrophils compared to B6 mice, albeit overall patterns compared to control injury and ECMtx remained the same. There was a significantly higher proportion of macrophages in control injuries in MRL mice compared to B6 mice. When evaluating these macrophages in more detail, we found that their phenotype was very different in MRL mice, as they had higher levels of CD68+ macrophages in all treatment groups with the most robust differences in control injuries and PEtx. Ternary plots of markers from the gene set “GOBP: Eosinophil mediated immunity” (Supplementary Figure 9a, b) and “Biocarta: Neutrophil degranulation” (Supplementary Figure 9c-d) showed PEtx and ECMtx skewing neutrophil and eosinophil marker expression respectively, further corroborating our flow cytometry data.

### MRL/MpJ mice display a stunted M2-macrophage polarization in response to ECM treatment

To visualize how biomaterial treatment changed gene expression in macrophages, we rescaled average log2(CPM) values of genes in the “Coates-Macrophage M1 vs M2 up” gene set across the three treatment groups and created ternary plots. We found that in C57BL/6 mice, genes that were previously ascribed to an M2 macrophage phenotype, including *Arg1*, *Pink1, Maf*, *and Chil3*, were potentiated by ECMtx (**Figure 4a**). In contrast, PEtx resulted in an M1 macrophage phenotype typified by skewed expression of *Marco, H2-D1*, *Myo1f*, *C1qb,* and *Thbs1* (**Figure 4a**); however, in MRL/MpJ mice that received ECMtx, the M2 macrophage signature was attenuated, most notably with reduced *Arg1* and *Chil3* expression (**Figure 4b**). Furthermore, PEtx in MRL/MpJ mice amplified the M1 macrophage signature seen in C57BL/6 mice, which received PEtx (**Figure 4b**). Tissue-level gene expression of the M2 macrophage markers *Arg1* and *Gata3* strongly correlated with the expression of *Retnla* and *Cdh1,* respectively, suggesting potential co-expression or co-regulation (Supplementary Figure 10). These data suggest either a preferential M1 macrophage recruitment or proliferation of tissue-resident macrophage population in MRL/MpJ mice after VML and material implantation. Additionally, the lipid-associated macrophage gene signature, including *Ctsb*, *Fabp4*, *Lpl,* and *Plin2,* was enriched in PEtx in both MRL and B6 mice (Supplementary Figure 11).

**Figure 4.**
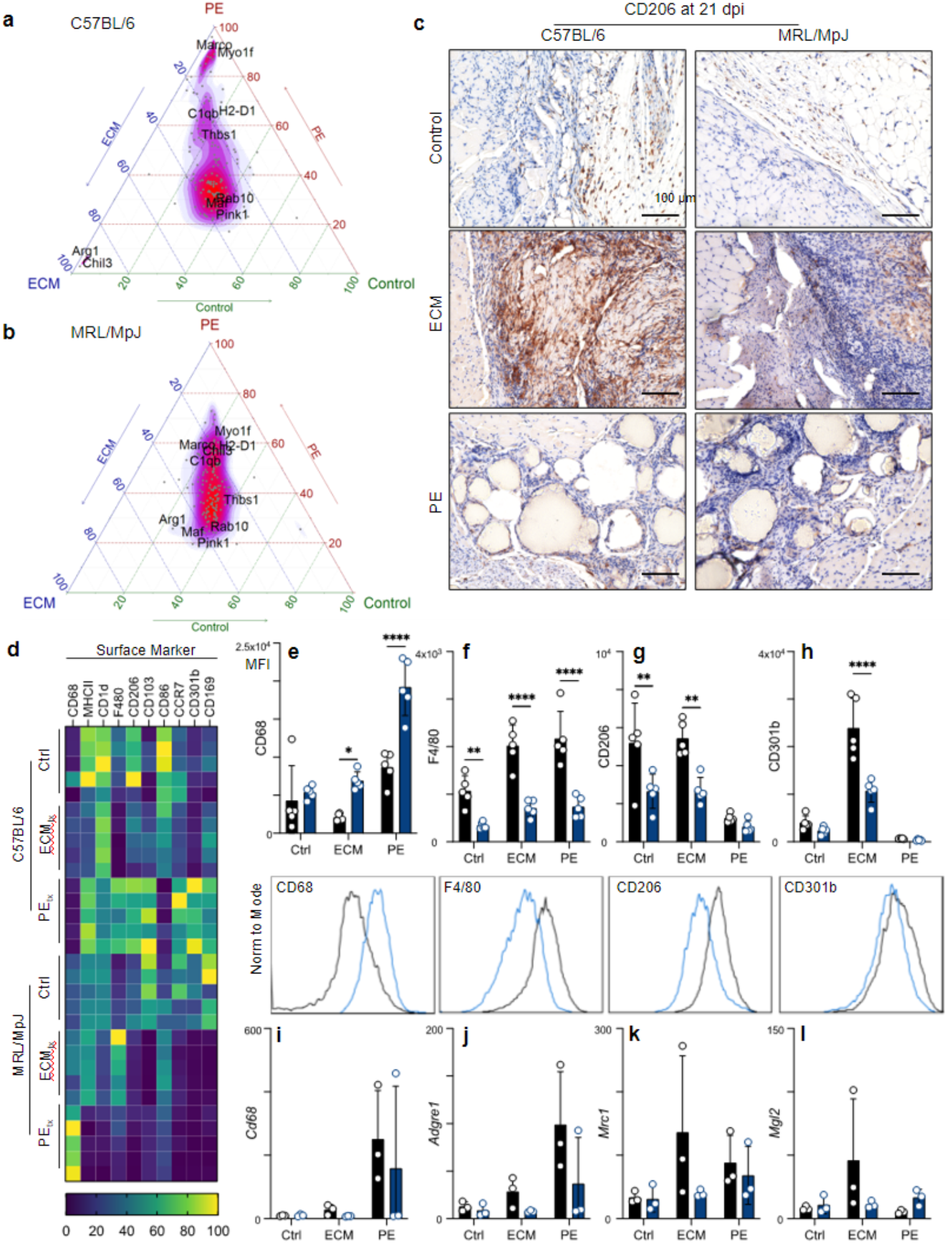
MRL/MpJ mice display decreased M2-macrophage polarization in response to ECM implants. (**a**) Ternary plot displaying normalized expression of genes in the gene set Coates-Macrophage M1 vs M2 up in C57BL/6 mice and (**b**) in MRL/MpJ mice. (**c**) Immunohistochemical staining of CD206 (DAB) counterstained with hematoxylin (purple) at 21 days post-injury (dpi). (**d**) Surface marker expression on macrophages by flow cytometry (mean fluorescence intensity, MFI) at 21 dpi. (**e** – **h**) MFI of (**e**) CD68, (f) F4/80, (**g**) CD206, and (**h**) CD301b with representative histograms beneath each plot. (**i** – **j**) Transcript counts for the genes encoding (**i**) CD68, (**j**) F4/80, (**k**) CD206, (**l**) CD301b. Data are means ± SEM, n = 5 (FACS, IHC) or 3 (RNAseq).

Correlated with CD11b, CD206 expressing cells were found in the fat pad against the injury site, with increased staining apparent in the B6 mice compared to the MRL mice (**Figure 4c**). This was even more apparent in the ECMtx mice, where robust CD206 staining was detected around the biomaterial in B6 mice, and while still present in MRL mice, it was much less apparent. There was minimal CD206 staining in PEtx mice, with a few CD206+ cells lying in the fibrous spaces between particles. There were large differences in surface marker expression by flow cytometry within the macrophages recruited to materials (**Figure 4d**). As previously mentioned, higher levels of CD68 were detected on macrophages in MRL mice with decreases in F4/80 expression (**Figure 4e, f**). When evaluating M2 macrophage markers, we repeated our finding that CD206 was significantly decreased in MRL mice compared to B6 mice (**Figure 4g**). Investigating further, we looked at CD301b expression (another M2 marker), wherein we found a significantly lower expression on macrophages in ECMtx MRL mice (**Figure 4h**). Comparing to RNA sequencing data (**Figure 4i- l**), we found that all of these patterns were recapitulated in the gene expression level, except for CD68 (**Figure 4i**). Interestingly, in PEtx mice, some markers of M2-like macrophage polarization were increased in MRL mice, such as *Retlna* (3.32-fold higher in MRL, p-adj = 0.006), *Pparg* (5.95, p-adj = 0.0006), *Cd163* (3.3, p-adj = 0.00003), and *Mgl2* (2.8-fold higher in MRL, p-adj = 0.005) (**Figure 4i**). Though overall M2-associated genes were significantly higher in ECMtx in comparison to PEtx (B6: *Retnla* 25-fold higher in ECMtx, p-adj = 2x10^-17^; *Cd163* 4.65-fold, p-adj = 0.001; *Mgl2* 8.77-fold, p-adj = 2x10^-6^).

### Regional alterations in T cell phenotype and activation in MRL/MpJ mice

In addition to the local tissue microenvironment, we detected differences in the draining lymph node through phenotyping adaptive immune cell presence and activation. In all three treatment groups, there was a higher proportion of ⍺β T cells than B cells in the inguinal lymph nodes in MRL mice (**Figure 5a**). There were minimal alterations in the more minor cell types, with a trending but insignificant decrease in γδ T cells in MRL mice (**Figure 5b**). Overall, there was a bias towards more CD4+ T cells in all treatment groups in MRL mice (**Figure 5c**). Due to the differences in the T and B cell populations in the draining lymph node, we evaluated the spleen to determine if there were differences in follicles and follicle structure. We found a decrease in white pulp and an increase in red pulp in MRL/MpJ mice with more defined follicles (T Cell/B Cell zones) in B6 mice (**Figure 5d**).

**Figure 5.**
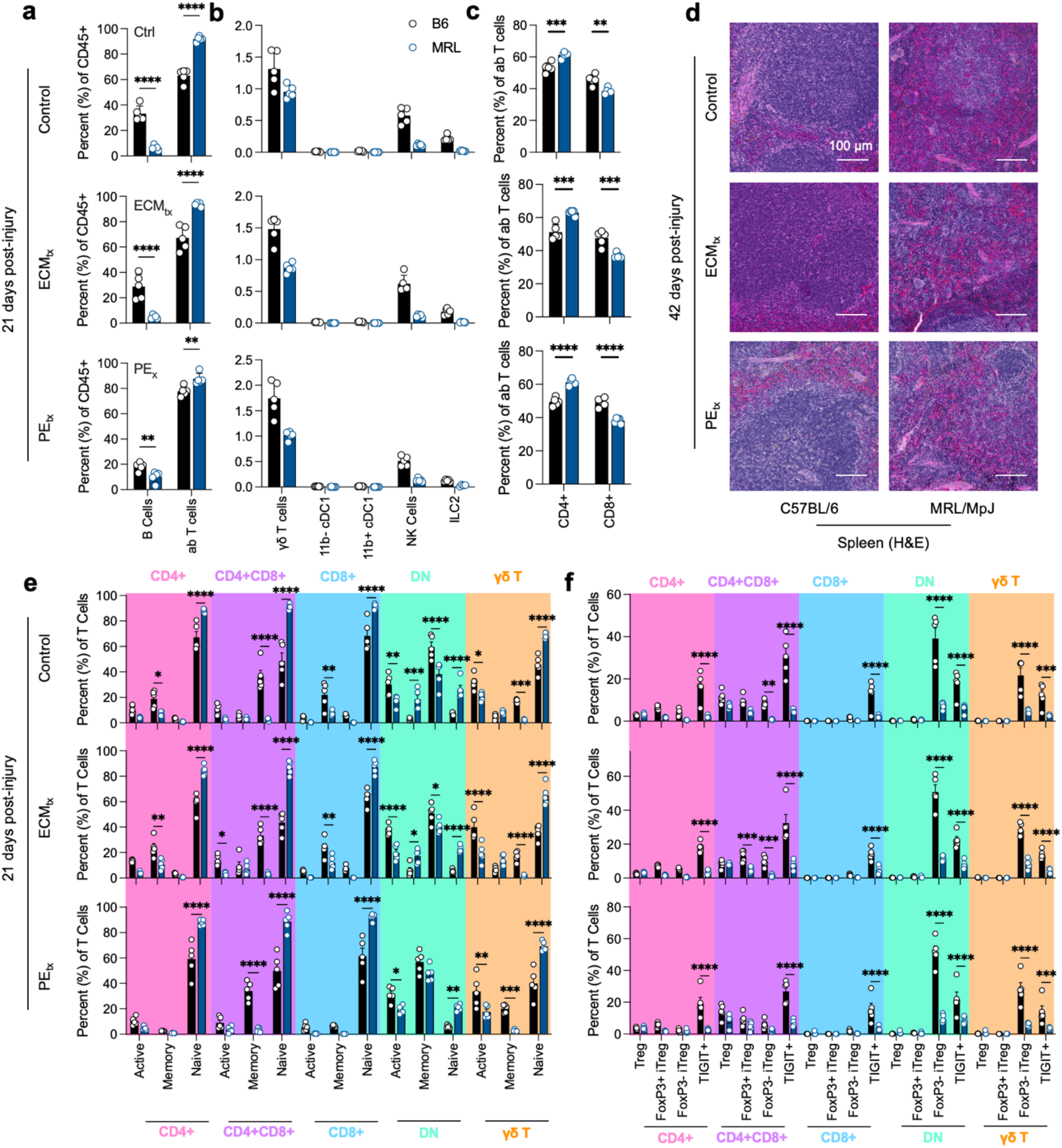
Decreased regional B and T cell responses to trauma in MRL/MpJ mice. (**a**) B cell and ⍺β T cell proportion in the inguinal lymph node at 21 days post-injury. (**b**) Proportions of lower prevalence adaptive immune cells at 21 days post-injury. (**c**) CD4 and CD8 T cell fractions as percent of ⍺β T cells. (**d**) Hematoxylin and eosin staining of spleen at 42 days post-injury. (**e**) T cell activation as determined by CD62L-CD44+ (active), CD62L+CD44+ (memory), and CD62L+CD44- (naïve). (**f**) Tregs in response to injury, Treg = Foxp3+HELIOS-, FoxP3+ iTreg = FoxP3+HELIOS+, FoxP3- iTreg = FoxP3-HELIOS+, TIGIT+. Data are means ± SEM, n = 5. ANOVA with Tukey post-hoc correction for multiple comparisons; * = P < 0.05, ** = P < 0.01, *** = P < 0.001, **** = P < 0.0001.

Due to the strong presence of T cells in MRL mice, we further analyzed the activation profile of these cells through the expression of CD62L and CD44 (**Figure 5e**). Overall, for all types of T cells tested, we found a lower proportion of Active and Memory T cells with a bias towards more naïve T cells (CD62L+CD44-) in MRL mice. The populations with the greatest proportion of active T cells (CD62L-CD44+) were double negative (CD4-CD8-) ⍺β T cells and γδ T cells. We also evaluated the presence of regulatory T cells based on the expression of FoxP3 and HELIOS, designating canonical and induced Tregs (**Figure 5f**). As with activation, there was a decrease in Tregs across all treatment groups in MRL mice. This was most prevalent with TIGIT+ Tregs (a marker of Tregs and immune exhaustion) and HELIOS+FoxP3- iTregs.

### Functional differences in foreign body giant cell formation

A major sign of the canonical foreign body response is the presence of multinucleate foreign body giant cells (FBGCs). We quantified FBGC response by calculating the number of giant cells (including Langhan cells with horseshoe nuclei) in hematoxylin and eosin-stained sections of PEtx VML (**Figure 6a**). In both B6 and MRL mice, minimal FBGCs were present 7 days post-injury. This increased in B6 mice by 21 days post-injury and was highest at 42 days post-injury. In contrast, MRL mice had significantly fewer FBGCs in response to PEtx, which peaked at 21 days and remained at this level –less than half of B6– at 42 days post-injury (**Figure 6b**). As mechanical cues can alter the immune response, especially in biomaterials, we evaluated the pre-injury mechanical properties of skeletal muscle in B6 and MRL mice through rheologic testing (**Figure 6c**). We found that MRL mice had a lower storage and loss modulus when compared to B6 mice, suggesting softer starting tissue mechanics. Interestingly, when evaluating mechano-sensing genes, we found a significantly higher gene count of *Piezo1*, a stiffness-sensitive ion channel, in MRL mice (**Figure 6d**). There were significant immune differences in the response to PE treatment in MRL mice, including decreased *Klra8* (and activating receptor for NK-mediated cell killing), higher *Cd1d* (associated with lipid antigen presentation), along changes in antigen presentation machinery (H2-associated genes) (**Figure 6e**).

**Figure 6.**
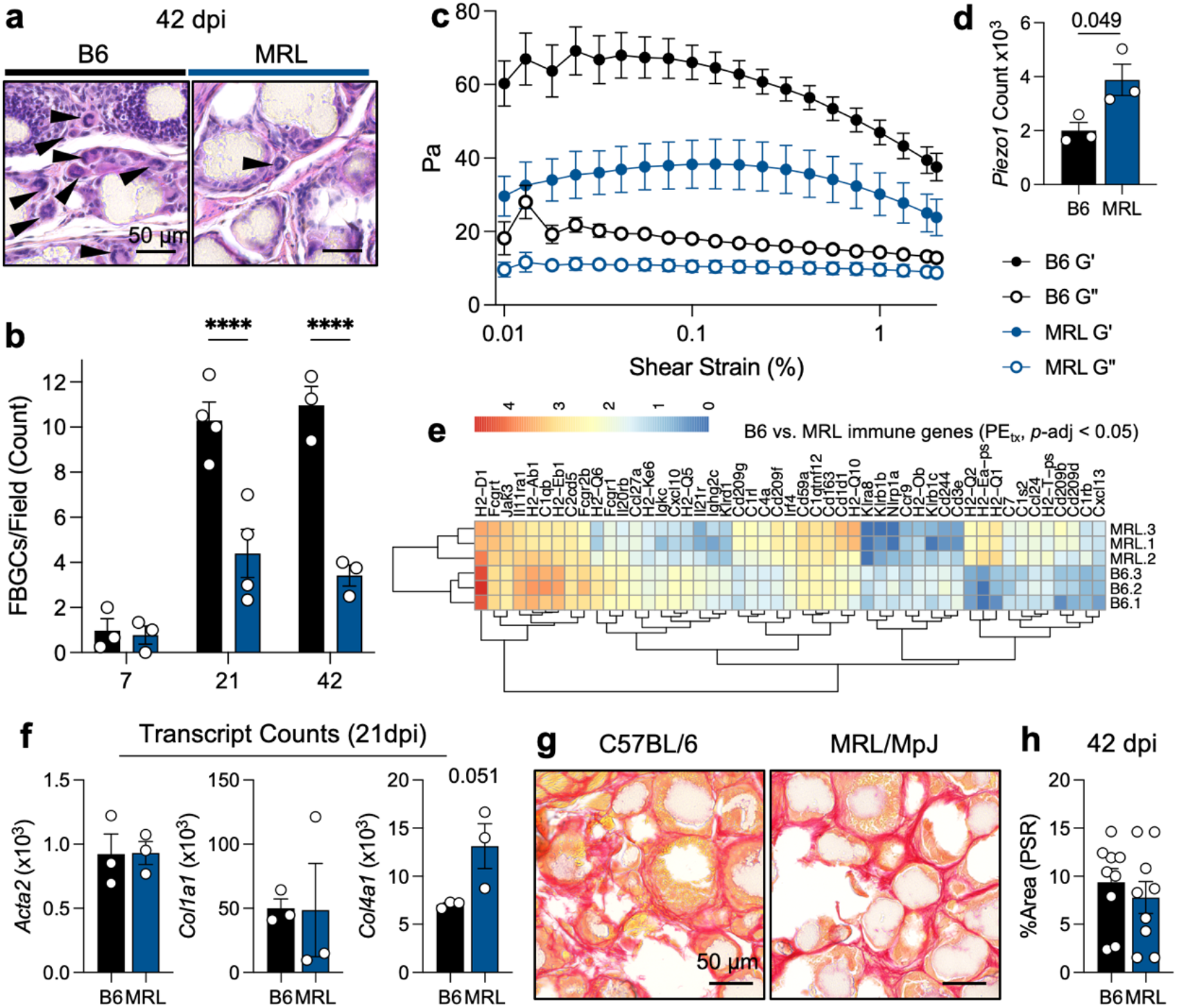
Differential fibrosis and foreign body giant cells correlate with the differences in muscle mechanics and immune response. (**a**) Representative images with foreign body giant cells in C57BL/6 (B6) and MRL/MpJ (MRL) mice at 42 days post-injury (dpi) by hematoxylin and eosin staining. (**b**) Quantification of foreign body giant cells (including Langhans cells) at 42 dpi. ANOVA with Tukey post-hoc correction for multiple comparisons, P < 0.0001. (**c**) Rheological properties of uninjured C57BL/6 (black) and MRL/MpJ (blue) quadriceps muscles. Closed circles = G’ (storage modulus), Open circles = G” (loss modulus). (**d**) *Piezo1* gene expression. (**e**) Immunologic response to PEtx muscle injury, significant genes with known protein products. Log-transformed counts. (**f**) *Acta2*, *Col1a1*, and *Col4a1* counts at 21 dpi. (**g**) Picrosirius red (PSR) staining at 42 dpi. (**h**) Quantification of thresholded PSR positive area per 200x field, n = 9 fields over 3 mice in each background. Data are means ± SEM.

When evaluating collagen deposition and fibrosis in response to PE particle implantation, we saw that canonical genes associated with fibrosis, such as *Acta2* (encoding aSMA) and *Col1a1* (encoding a collagen I subunit) did not significantly change between B6 and MRL mice (**Figure 6f**). *Col4a1*, a gene encoding non-fibrotic collagen, was higher in all MRL mice than B6 mice but was not significantly due to variability in MRL mice (P = 0.051). Through staining for collagen I with picrosirius red (PSR), we found that both B6 and MRL mice exhibited dense collagen deposition between PE particles 42 days post-injury (**Figure 6g**). When quantified, there was no significant difference in fibrotic collagen deposition in B6 versus MRL mice, agreeing with the RNA sequencing data showing no difference in *Col1a1* gene expression (**Figure 6h**). We also saw no significant differences between B6 and MRL in their normalized grip strength within the trimeframe tested (Supplementary Figure 12).

## DISCUSSION

Wound healing and tissue regeneration in higher mammals have been restricted to specific tissues – such as the liver –. In contrast, the capacity for complex tissue regeneration is retained in amphibians who can regenerate full limbs. The Murphy’s Roth’s Large (MRL) mouse model was developed for its size (from the LG strain), but a spontaneous mutation arose that resulted in lymphoproliferative disorder and autoimmune phenotypes (MRL/lpr). This was traced to a mutation in Fas. However, when bred out, the MRL/+ (“MRL/MpJ”) strain still exhibited autoimmunity. It exhibited a strong ability to regenerate ear punches used in mouse identification, as seen in both parent MRL/lpr and LG strains. This variability of mouse strains in regeneration capacity can be leveraged for tissue engineering studies to identify therapeutic targets [33]. There have been reports of variability in the regenerative capacity of the MRL mouse, with differences in skin wound healing depending upon where the skin is injured and conflicting reports on the recovery from myocardial infarction [34-36]. Large-scale volumetric injuries are less studied in this model, and we found differences in the responses to traumatic skeletal muscle injury in MRL/MpJ and C57BL/6 mice. Surprisingly, as CD206+ macrophages have previously been associated with a pro-regenerative phenotype, while we hypothesized a more robust M2-like macrophage polarization, we found a decrease in M2-related genes as detected by RNA sequencing and confirmed via flow cytometry. This may relate to the persistence of CD206+ macrophages associated with pathologic outcomes in tissue regeneration and wound healing [37, 38]. As type-1 immune signals are needed for myoblast proliferation, this type-2 switch, if occurring too early, may stunt regeneration by signaling the transition to myotube fusion before sufficient precursor cell development [39, 40]. When evaluating outcomes in response to materials that elicit a more canonical foreign body response, we found a decrease in the number of foreign body giant cells (FBGCs) in MRL mice with no changes in fibrosis as determined by picrosirius red staining and gene expression patterns. FBGCs, multinucleate fused macrophages, can be stimulated to fuse through type-2 immune signals but are associated with more type-1 inducing materials like polyethylene used in our studies [41, 42].

There is a reciprocal relationship between tissue microenvironment and immune responses. Firstly, the immune microenvironment of different tissues can alter the downline response to a challenge such as an infection, injury, or device implantation. Adipose tissue is a known reservoir of immune cells; adipocytes are both energy-storage and endocrine regulators affecting other cells within and neighboring a fat depot. Adipose tissue communicates the metabolic stage to the immune system via hormones called adipokines (adiponectin, leptin, omentin, etc.), which control immune cell activity [43]. The two most studied adipokines, adiponectin, and leptin, have been shown as key regulators of cardiac and skeletal muscle function and growth [44]. Adiponectin has been revealed in studies of muscle development, regeneration, and regulation of inflammation [45]. In the presented study, *Adipoq* expression was trending lower in pre-injury but higher in post-injury MRL mice (in comparison to their corresponding B6 group). In addition, white fat bears many resident immune cells, including M2-like macrophages, conventional dendritic cells (cDCs), and regulatory T cells [12]. Previous work has shown increased fibrosis in materials implanted subcutaneously, especially near adipose tissue [46]. Cellularly, there is a strong MHCII+ B cell recruitment to materials implanted intraperitoneally versus those in the subcutaneous space where there are differences in fat composition between visceral and subcutaneous fat depots [29]. Just as the tissue microenvironment influences the response to a challenge, the immune response to injury can subsequently affect tissue development. As mentioned previously, this has been shown in skeletal muscle where signals like IFNγ promote the proliferation of myoblasts to provide a suitable precursor population for new tissue development, followed by an IL-10 mediated switch to type-2 immune activation which, through IL-4, has specifically been shown to promote fusion into myotubes [47]. IL-33-mediated accumulation of Tregs has been associated with positive outcomes in muscle regeneration, and IL-33 is a prominent cytokine enriched in white fat [12, 48]. After a VML, we find that the inguinal fat pad migrates to the injury space, possibly as a depot of immune signaling. Furthermore, after injury and biomaterial implantation, we see an increase in adipogenesis in the MRL mice compared to the B6 mice, confirmed via RNAseq and histologic analyses. Due to the integrated nature of muscle fibers and white fat, we believe this is due to ectopic adipogenesis within tissue defects as opposed to fat pad migration. As white fat can quickly swell, and skeletal muscle is prone to collapse after trauma, causing further damage, the fat may play an additional space-filling role to account for the physical void created by the volumetric injury.

Previous reports through crossing with MRL with B6 mice have shown that the autoimmune phenotype does not significantly correlate with the wound healing phenotype as determined by ear punch closure rate and lymph node cell counts [49]. Antigen specificity of wound healing has been implied, especially in the context of canonical CD4+FoxP3+ Tregs that have repeatable, clonally expanded CDR3s across multiple mice tested [50]. We see a significantly different level of T cell activation with a decreased proportion of Naïve T cells and an increase in Treg formation in B6 mice in the draining lymph nodes compared to their MRL counterparts. Furthermore, CCR9 expression on dendritic cells has previously been associated with self-tolerance and regulating autoimmunity [51]. We see lower levels of the *Ccr9* transcript in MRL mice treated with PE compared to B6 mice, suggesting some evidence of more autoimmune-like profiles in MRL response to injury, albeit the outcome is not yet clear. Though a direct link between autoreactivity and post-trauma tissue development has yet to be established, there are multiple correlations between fat formation and immunologic response to injury. This includes lipid-associated macrophages in MRL mice and upregulation of the transcript encoding CD1d, which assists in NK-cell mediated presentation of lipid antigens to adaptive immune cells.

When evaluating the differences between MRL and B6 mice that lie beyond immunologic responses, we also found differences in the mechanics of the muscle tissue, with MRL mice having rheologically “softer” muscles with a lower storage modulus than B6 mice. Previous reports have shown that substrate stiffness can greatly alter the outcome of stem cell polarization; specifically, mesenchymal stem cell (MSC) differentiation on stiffer substrates promoted a more osteogenic lineage commitment, whereas softer substrates promoted more adipogenesis [52]. Not only the mechanics of the tissue but mechanically sensitive genes, such as *Piezo1,* a stiffness-sensitive ion channel, were different in B6 and MRL mice with response to injury, with MRL mice expressing higher levels of *Piezo1* mRNA.

## CONCLUSIONS

Previous reports have shown that the MRL/MpJ genetic background predisposes mice to autoimmunity, accelerated cutaneous wound healing [53], and reduced scarring [54]. However, in wound healing and tissue regeneration, immune response to traumatic muscle injury and biomaterial implantation in this background, specifically in contrast to the well-studied C57BL/6 mice, had yet to be delineated. Here, we show that in contrast to C57BL/6 mice, MRL/MpJ mice display ectopic adipogenesis within the muscle injury site after VML and fibrotic material implantation. First, through histology and bulk RNA sequencing, we observed that white fat and an adipogenic gene signature were enriched in MRL/MpJ mice muscles. This further correlated with an M1 macrophage gene set, specifically potentiated in mice that received PEtx. Using flow cytometry, we confirmed the enrichment of CD68+ F4/80- M1 macrophages in all MRL/MpJ mice, most notably in PEtx compared to C57BL/6 mice, which received identical treatments. Further, we showed that MRL/MpJ mice displayed attenuated local B and T cell responses in response to traumatic muscle injury and biomaterial implantation. The unexpected finding of ectopic adipogenesis in MRL/MpJ mice and the underlying differences in immune response in contrast to C57BL/6 mice may yield insight into the dynamics between stem cells and the tissue environment during post-trauma tissue development. The cell types identified in this study, along with the atypical phenotypic presentation and correlations with gene expression, could serve as a template for understanding tissue development, mechanics, immune response to novel biomaterials, and identifying potential targets for therapeutic development.

## ACKNOWLEDGMENTS

The authors would like to acknowledge Dr. Lisa Portnoy and Monica Bur for their guidance on animal studies. We would also like to acknowledge Vanathi Sundaresan for laboratory organization and support and Dr. Nicole Morgan and the Biophysics, Engineering, and Physical Science Shared Resource of NIBIB for assistance with rheology and helpful discussions. This work was funded by the intramural research program of the National Institutes of Health, National Institute of Biomedical Imaging and Bioengineering, National Institutes of Health (NIH). <u>Disclaimer</u>: The contents of this publication are the authors’ sole responsibility and do not necessarily reflect the views, opinions, or policies of the NIH and the Department of Health and Human Services (HHS). Mentioning trade names, commercial products, or organizations does not imply endorsement by the U.S. Government.

## DATA AND CODE AVAILABILITY

All data associated with this study, including flow cytometry experiments and histology images, are present within this manuscript and associated supplementary information. Raw sequencing data is available at The Sequence Read Archive (SRA; project number: [*to be deposited upon manuscript acceptance]]*). The code required to analyze the sequencing data and the associated study design and R markdown files are available at Code-Ocean as a reproducible “capsule” and as a PDF in the supplementary information.

## CONFLICT OF INTEREST

Authors have no conflict of interest to declare.

## SUPPLEMENTARY MATERIALS

Ngo & Josyula et al.

**Supplementary Figure 1.**
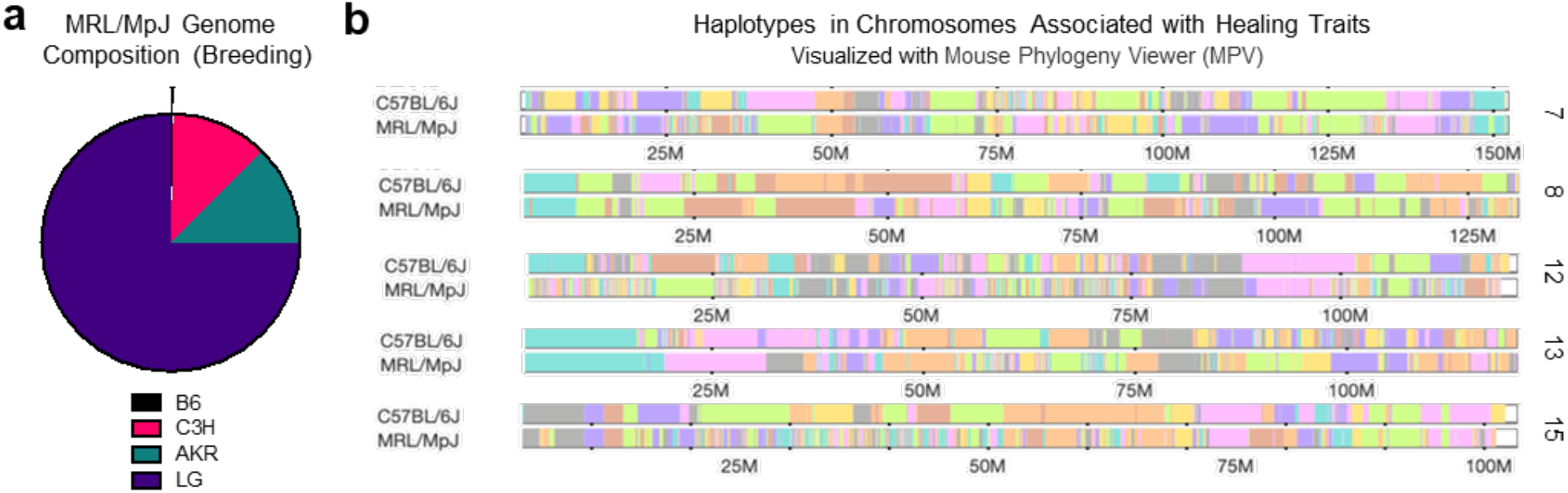
(a) Genomic composition of MRL/MpJ mouse from parent strains (b) Haplotype display on chromosomes 7, 8, 12, 13, and 15 from Mouse Phylogeny Viewer JR Wang, F Pardo-Manuel de Villena, and L McMillan. *Comparative analysis and visualization of multiple collinear genomes*. BMC Bioinformatics, 2012.

**Supplementary Figure 2.**
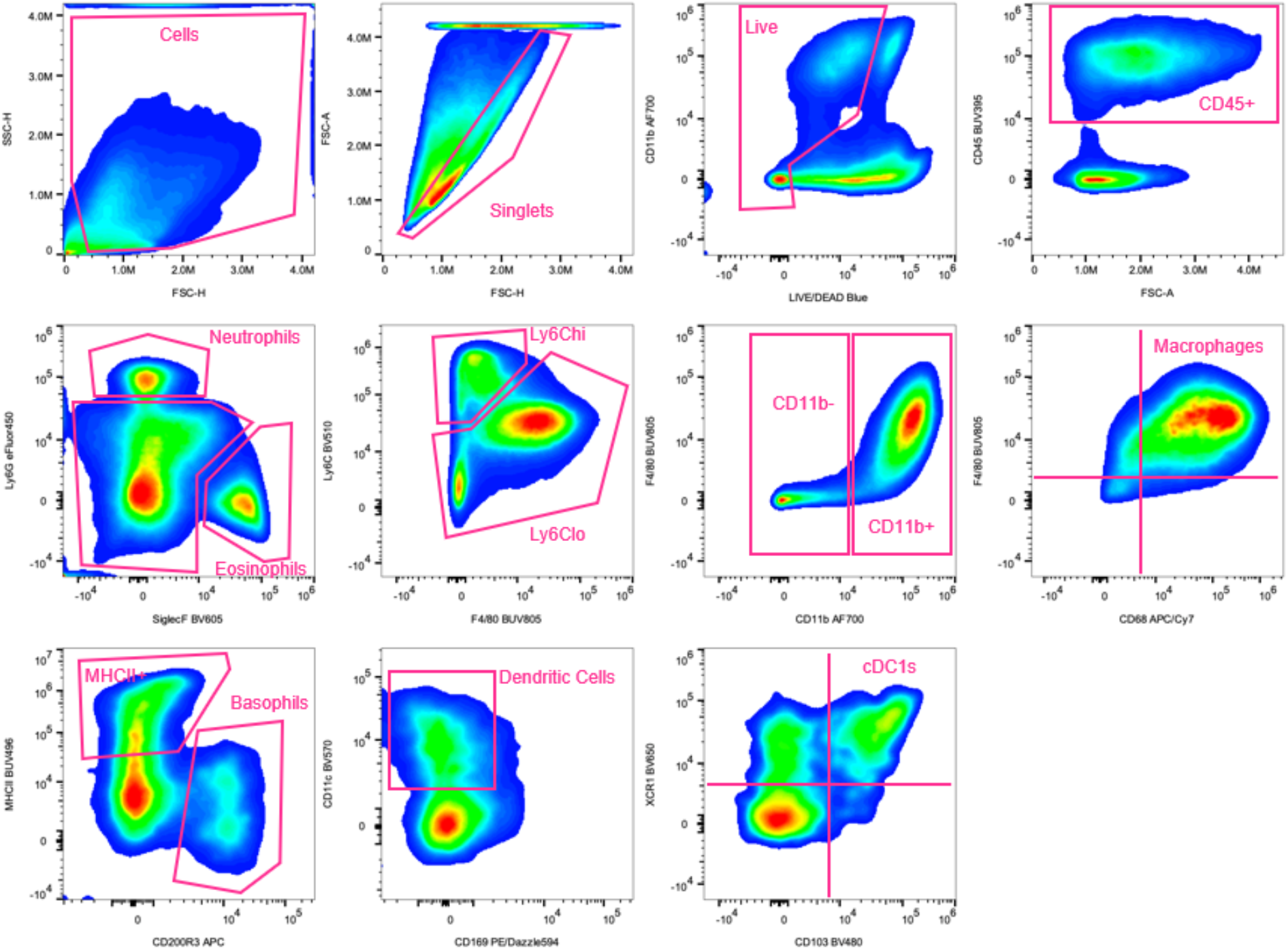
Gating Strategy for myeloid flow cytometry panel. Top row = control injury at 21 days post-op. The bottom two rows = Control, ECMtx, and PEtx concatenated (21 days post-injury). C57BL/6 mouse.

**Supplementary Figure 3.**
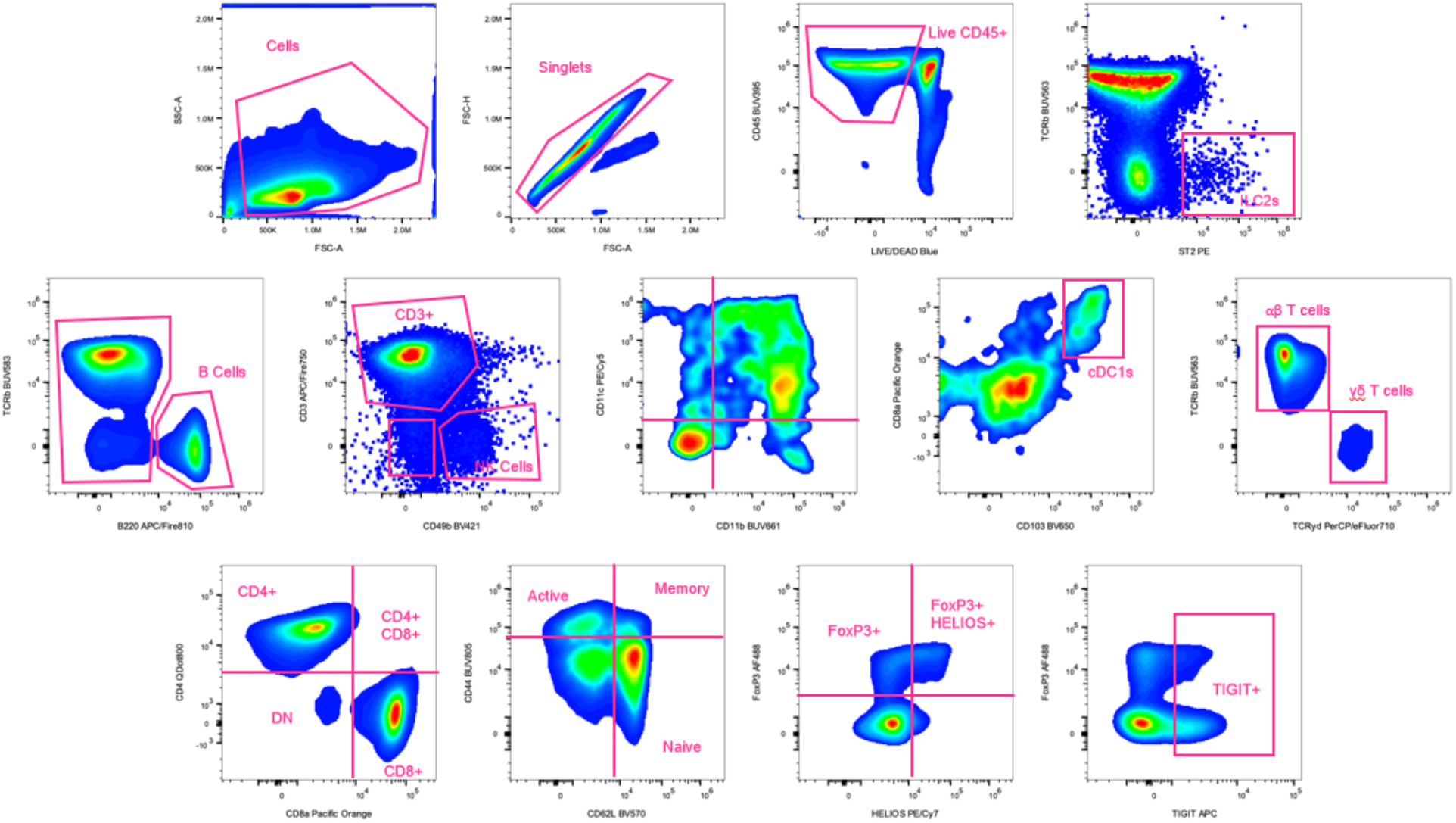
Gating Strategy for lymphoid flow cytometry panel. Control injury at 21 days post-injury. Inguinal lymph node from C57BL/6 mouse.

**Supplementary Figure 4.**
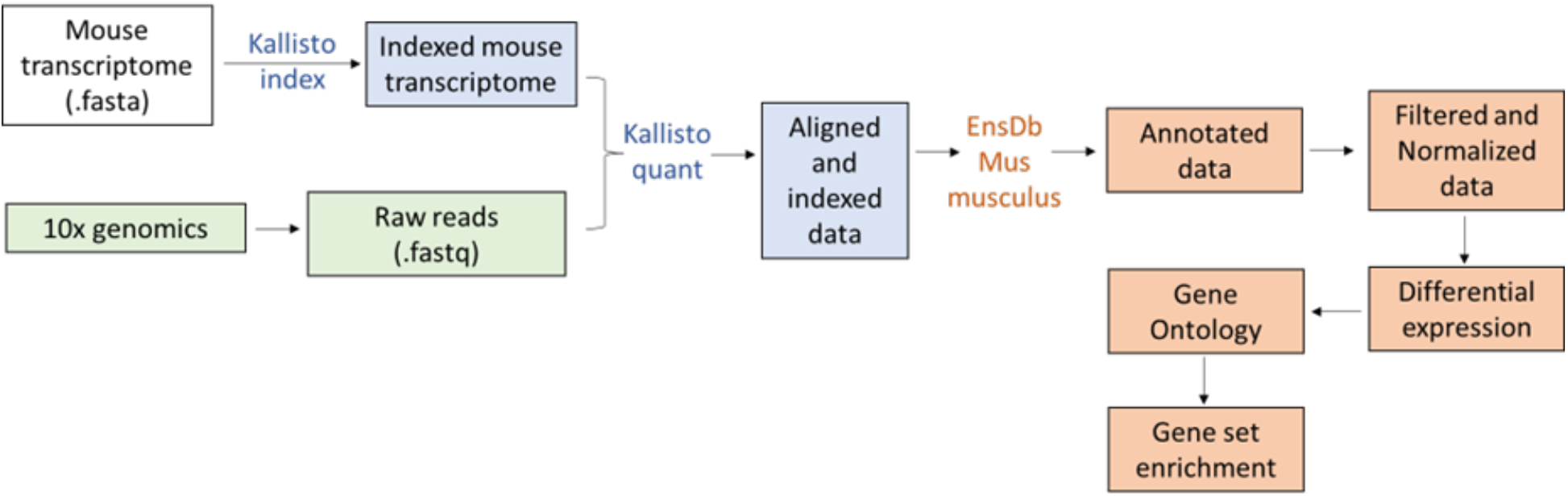
Schematic of bulk RNA sequencing data analysis workflow. Panels in green represent data obtained directly from Azenta Genewiz. Panels in blue represent data analyzed using the command line program Kallisto. Panels in red represent data analyzed using R (version 4.2.1 ‘Funny-looking kid’).

**Supplementary Figure 5.**
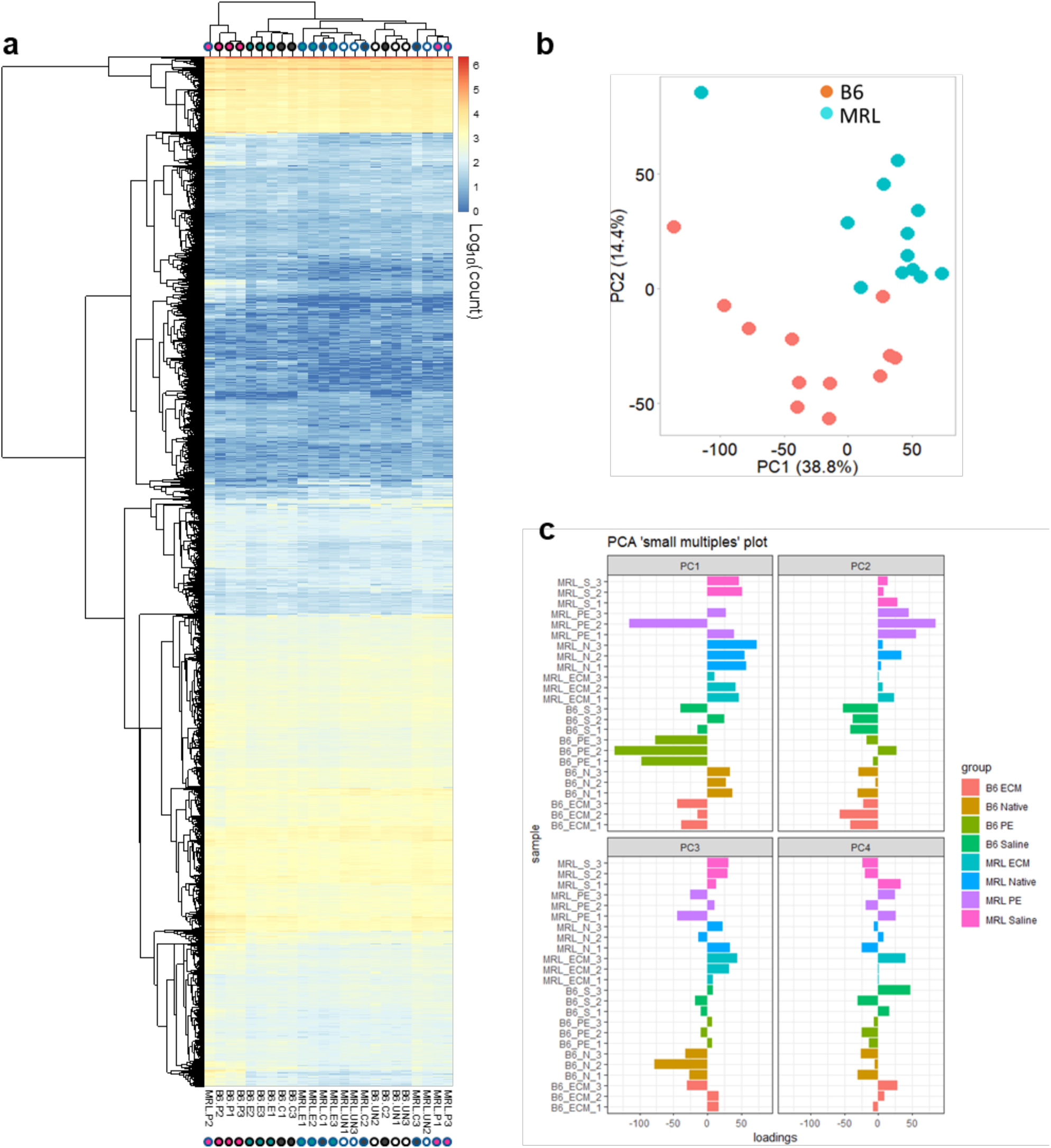
(a) Heatmap of differentially expressed genes across all samples with dendrograms representing clustering performed by row (left) and column (top). (b) Principal component analysis of all samples with dot plot showing variation represented by PC1 and PC2 (top) and (c) small multiples plot of principal components 1-4. The color gradient scale bar in (a) represents Log10(CPM).

**Supplementary Figure 6.**
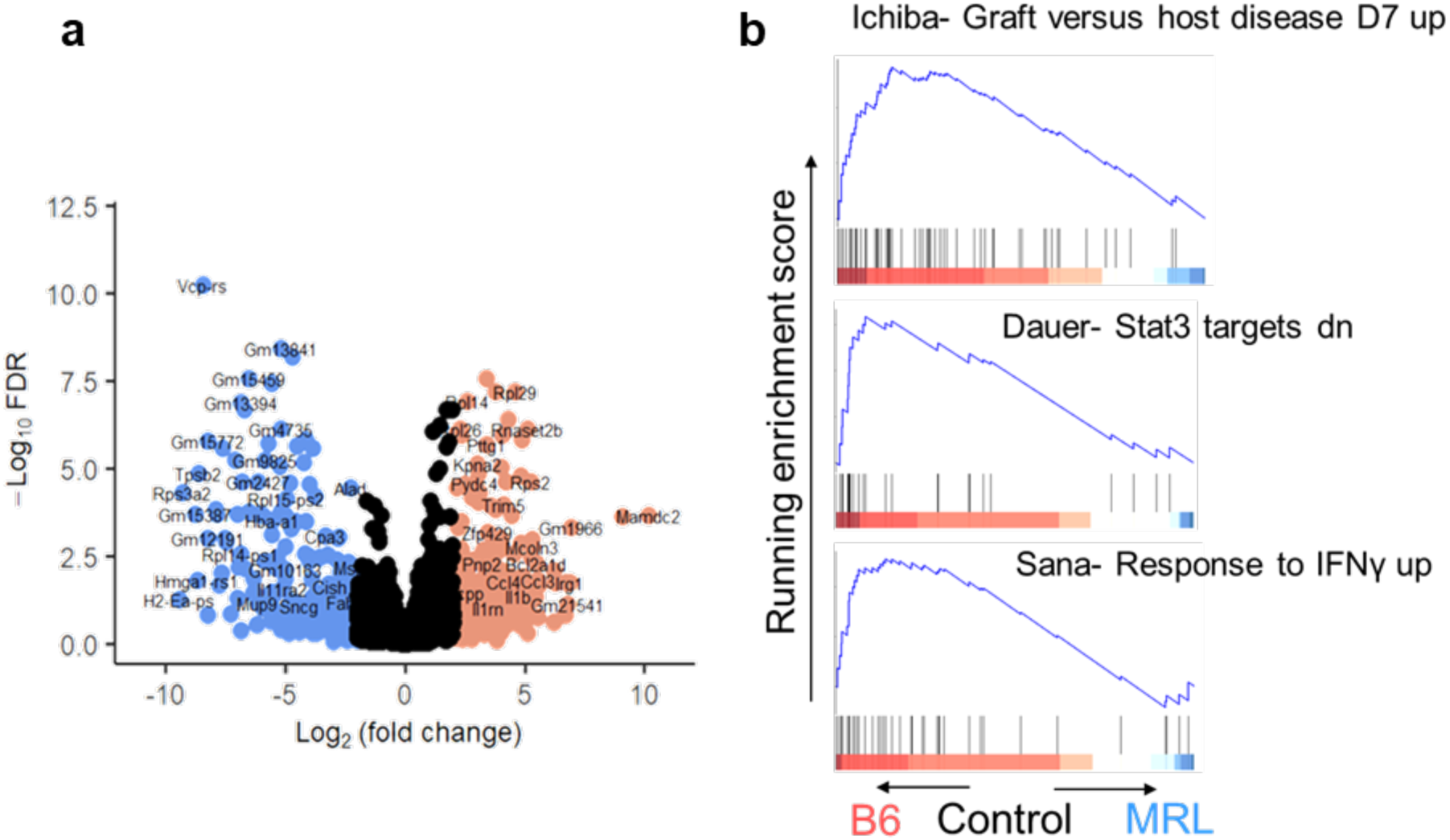
(a) Volcano plot of differentially expressed genes in the control injured group in B6 versus MRL mice. (b) GSEA of top 250 differentially expressed genes from (a) with relevant gene sets displayed.

**Supplementary Figure 7.**
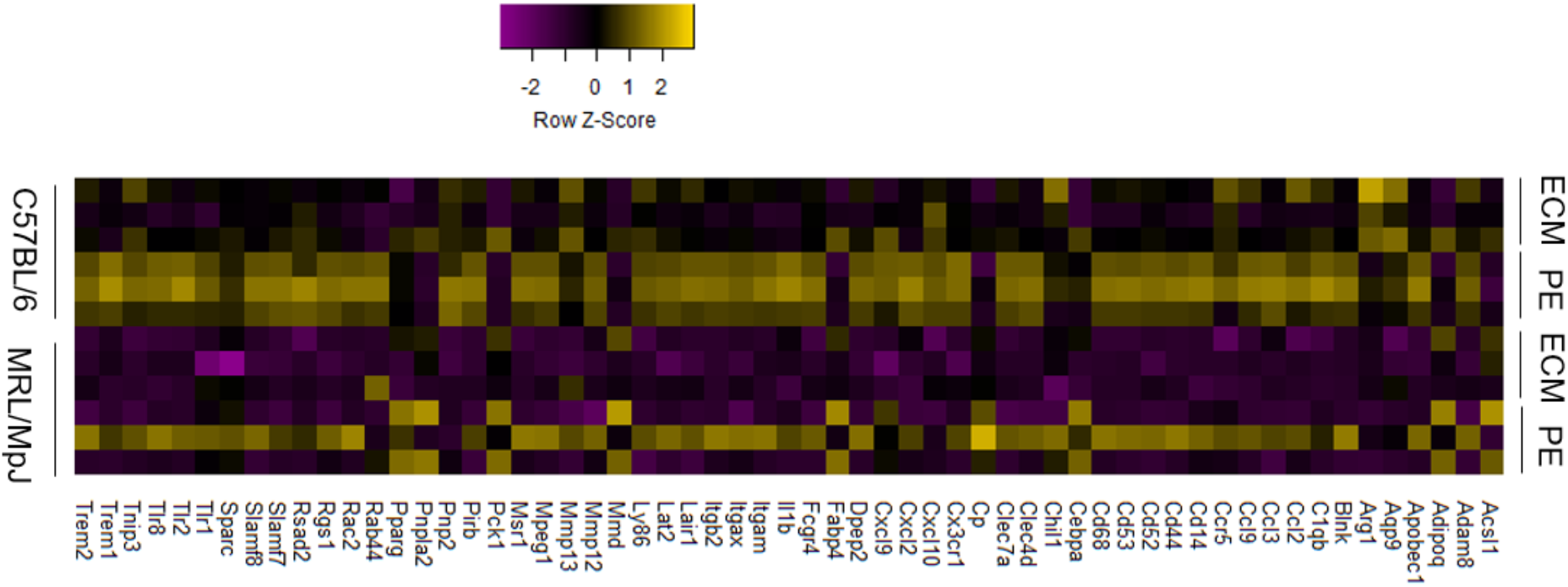
Heatmap of genes potentiated by PEtx compared to ECMtx in C57BL/6 and MRL/MpJ mouse strain.

**Supplementary Figure 8.**
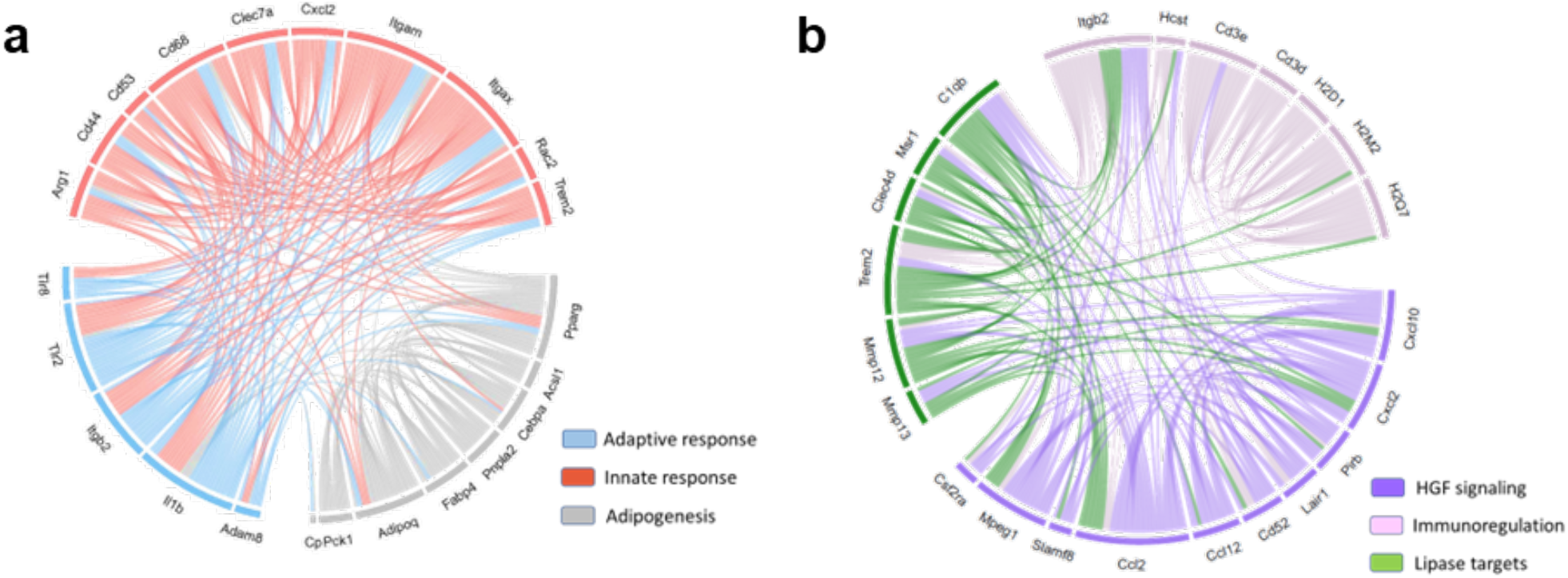
Chord diagrams of protein-protein interactions sourced from the string protein database. Interactions of leading-edge markers from gene sets enriched in (a) PEtx and (b) ECMtx obtained from top 250 differentially expressed genes across B6 and MRL mice.

**Supplementary Figure 9.**
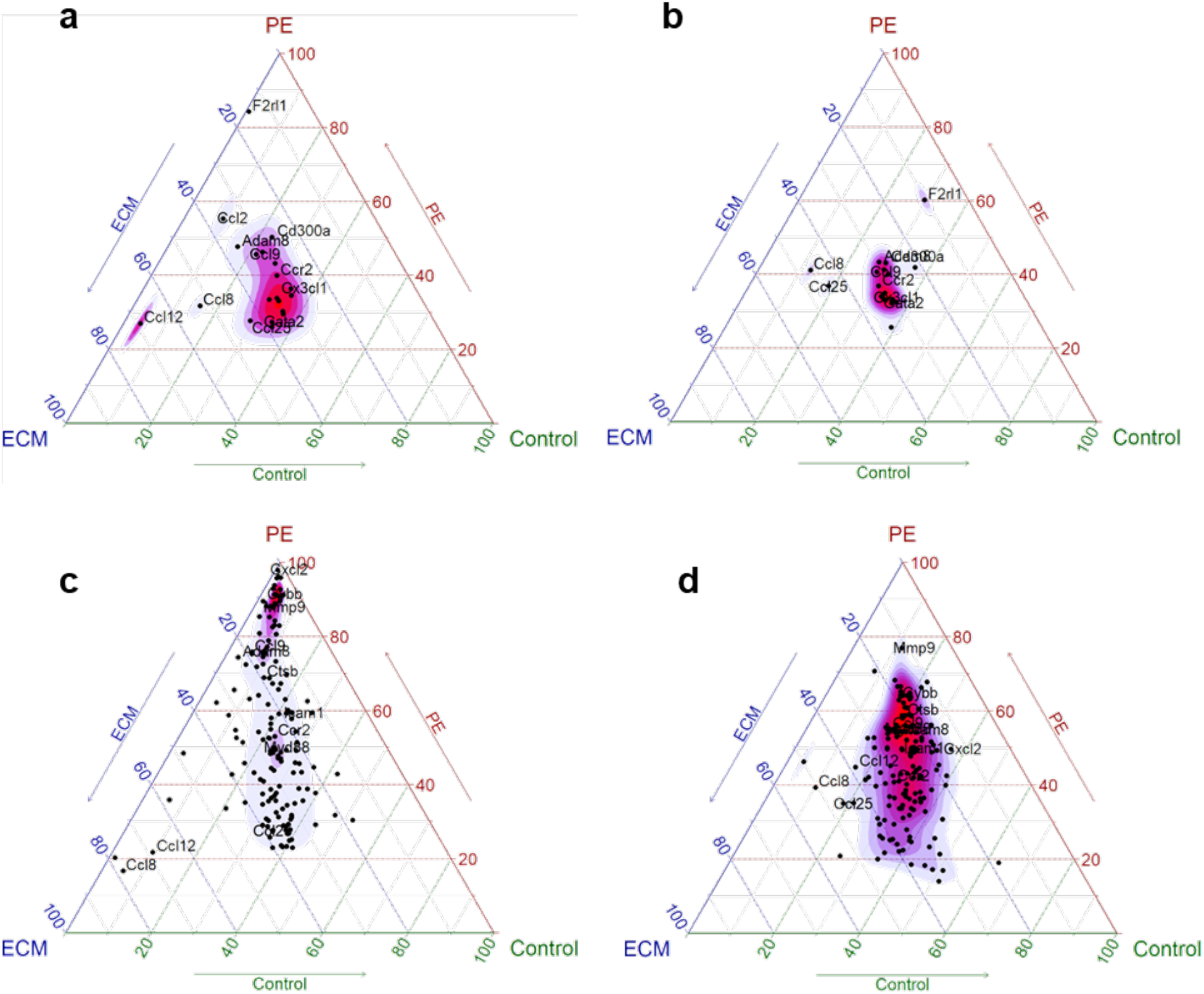
Ternary plots showing rescaled normalized log2(CPM) gene expression of eosinophil markers in (a) C57BL/6 and (b) MRL/MpJ mice. Similarly, neutrophil markers in (c) C57BL/6 and (d) MRL/MpJ mice. All data points represent individual genes’ average log2(CPM) values at 21 days post VML and material implantation. Eosinophil markers were subset from the gene set “GOBP: Eosinophil mediated immunity” and neutrophil markers from “Biocarta: Neutrophil degranulation.”

**Supplementary Figure 10.**
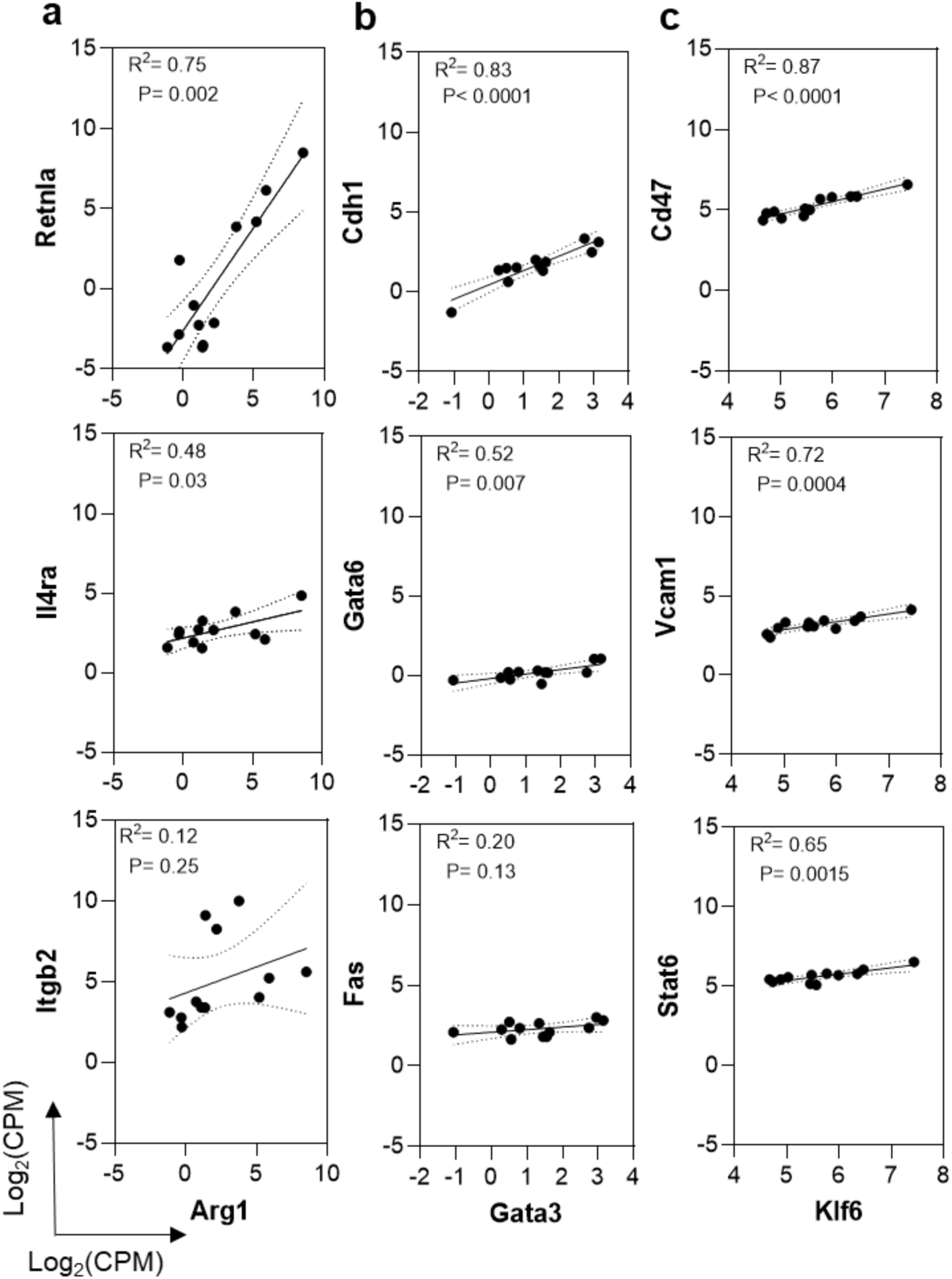
The M2 macrophage markers (a) *Arg1,* (b) *Gata3,* and (c) *Klf6* were strongly co-expressed (R^2^ ≥ 0.75) with *Retnla, Cdh1,* and *Cd47,* respectively, in all B6 mice across all treatments.

**Supplementary Figure 11.**
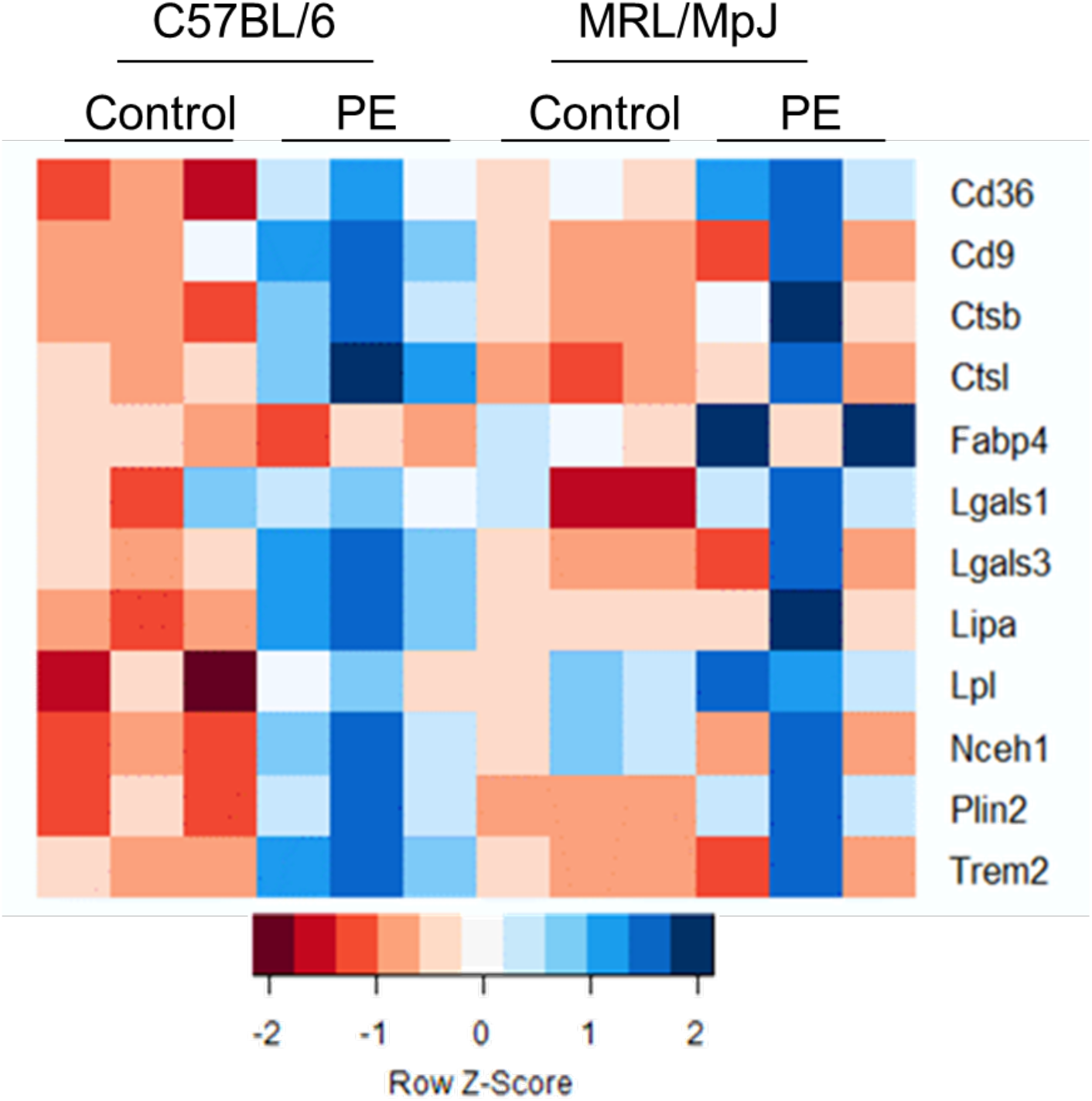
Heatmap of genes comprising lipid-associated macrophage signature enriched in mice that received PEtx compared to saline controls.

**Supplementary Figure 12.**
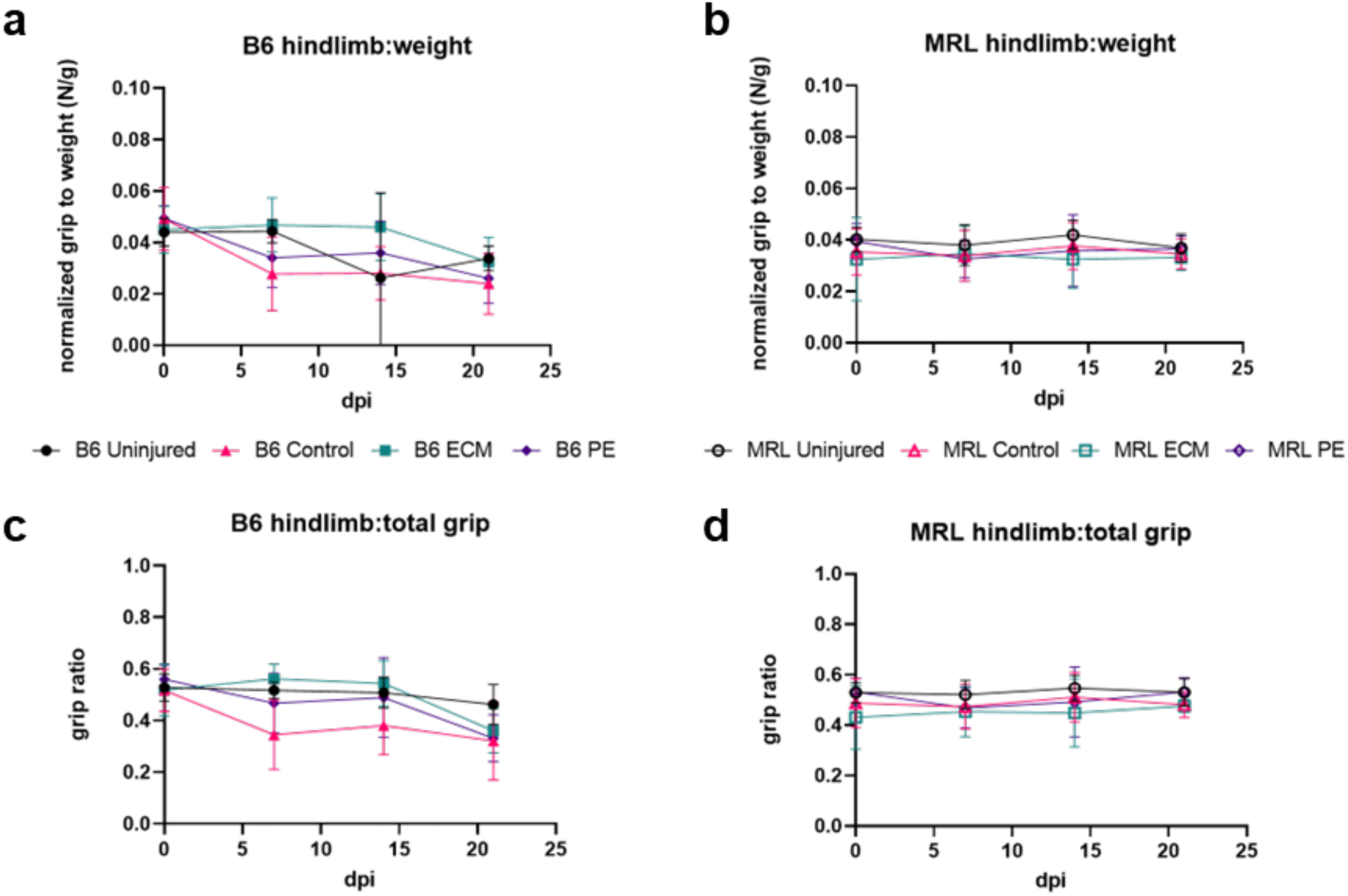
| Normalized grip strength of B6 versus MRL mice pre- and post-injury with different material implantations. **(a)** and **(b)** Hindlimb grip strength measurements normalized to body weight at pre-injury and 7-, 14-, and 21-days post-injury of B6 versus MRL mice, respectively. **(c)** and **(d)** Hindlimb grip strength measurements normalized to total 4-limbs strength at pre-injury and 7-, 14-, and 21-days post-injury of B6 versus MRL mice, respectively. dpi = days post-injury. Black = uninjured, pink = control injury, teal = ECMtx, purple = PEtx. Data are means ± SEM.

**Supplementary Table 1.** Bulk RNA sequencing quality control metrics for all 24 samples in this study, including mean quality score and total number of reads per sample.

| Sample ID | Barcode Sequence | # Reads | Yield (Mbases) | Mean Quality Score | % Bases >= 30 |
| --- | --- | --- | --- | --- | --- |
| B6-PE-2 | GTAGAGGA+C<br>TTCGCCT | 23341152 | 7002 | 35.55 | 91.8 |
| B6-Uninjured-3 | GCTCATGA+T<br>CAGAGCC | 30067196 | 9020 | 35.58 | 91.93 |
| B6-PE-1 | GTAGAGGA+T<br>CAGAGCC | 32820587 | 9846 | 35.52 | 91.65 |
| B6-Uninjured-1 | GCTCATGA+G<br>CCTCTAT | 30468441 | 9141 | 35.55 | 91.77 |
| B6-Uninjured-2 | GCTCATGA+A<br>GGATAGG | 29612058 | 8884 | 35.59 | 91.99 |
| B6-PE-3 | GTAGAGGA+T<br>AAGATTA | 33383202 | 10015 | 35.6 | 92.06 |
| MRL-ECM-1 | GCTCATGA+C<br>TTCGCCT | 26235642 | 7871 | 35.62 | 92.11 |
| MRL-ECM-2 | GCTCATGA+T<br>AAGATTA | 31055947 | 9317 | 35.67 | 92.4 |
| MRL-PE-1 | GCTCATGA+G<br>TCAGTAC | 30772510 | 9232 | 35.58 | 91.96 |
| MRL-Uninjured-3 | ATCTCAGG+G<br>TCAGTAC | 39984070 | 11995 | 35.53 | 91.68 |
| B6-Saline-2 | GTAGAGGA+G<br>TCAGTAC | 34568555 | 10371 | 35.5 | 91.56 |
| MRL-PE-2 | ATCTCAGG+A<br>GGCTATA | 35226447 | 10568 | 35.47 | 91.42 |
| MRL-Uninjured-2 | ATCTCAGG+A<br>CGTCCTG | 41136587 | 12341 | 35.65 | 92.29 |
| MRL-Uninjured-1 | ATCTCAGG+T<br>AAGATTA | 36836367 | 11051 | 35.62 | 92.11 |
| B6-ECM-2 | GTAGAGGA+G<br>CCTCTAT | 27006422 | 8102 | 35.5 | 91.58 |
| MRL-ECM-3 | GCTCATGA+A<br>CGTCCTG | 25466670 | 7640 | 35.62 | 92.21 |
| B6-Saline-1 | GTAGAGGA+A<br>CGTCCTG | 29484030 | 8845 | 35.61 | 92.11 |
| B6-ECM-3 | GTAGAGGA+A<br>GGATAGG | 31221810 | 9367 | 35.56 | 91.88 |
| MRL-Saline-2 | ATCTCAGG+T<br>CAGAGCC | 34998055 | 10499 | 35.54 | 91.71 |
| B6-Saline-3 | GCTCATGA+A<br>GGCTATA | 25044049 | 7513 | 35.53 | 91.73 |
| MRL-PE-3 | ATCTCAGG+G<br>CCTCTAT | 30143707 | 9043 | 35.52 | 91.61 |
| B6-ECM-1 | GTAGAGGA+A<br>GGCTATA | 28875619 | 8663 | 35.51 | 91.63 |
| MRL-Saline-1 | ATCTCAGG+A<br>GGATAGG | 36356638 | 10907 | 35.51 | 91.59 |
| MRL-Saline-3 | ATCTCAGG+C<br>TTCGCCT | 24880999 | 7464 | 35.6 | 92 |

**Supplementary Table 2.** Full Gene set names displayed in GSEA plots. Gene set names in figures were abridged for brevity.

| Name used in figure | Full gene set name |
| --- | --- |
| Adipogenesis | Burton Adipogenesis |
| Adaptive Immune Response | Reactome-Adaptive Immune System |
| Innate Immune Response | Reactome- Innate Immune System |
| HGF Signaling | Rutella-Response to HGF Up |
| Lipase targets | Lian Lipa Targets 3M |
| IL4 singlaing | GOBP: Interleukin 4 mediated signaling pathway |
| Neutrophils | Reactome- Neutrophil Degranulation |
| TNF-p38 | Phong- TNF response via p38 complete |
| Lipase targets | Lian Lipa Targets 3M |
| Ribosome | KEGG- Ribosome |
| Fibrosis | WP-Overview of proinflammatory and profibrotic mediators |
| Steroid | Reactome- Metabolism of Steroids |
| Lipids | Reactome- Metabolism of Lipids |
| Energy | Reactome- Integration of energy metabolism |
| Mitochondria | Mootha Mitochondria |
| Pyruvate | Reactome- Pyruvate Metabolism |
| Cytokine receptor | KEGG- Cytokine-Cytokine Receptor Interaction |
| M2 Macrophage | Coates- Macrophage M1 vs M2 |

**Supplementary Methods** | PDF document with code used to analyze bulk RNA sequencing data. See subsequent pages of this document.

produced on 2023-09-20

## 1 Background

Murphy roths large (MRL/MpJ) mice are known to regenerate skin and skeletal muscle wounds without scar formation but are also susceptible to metabolic dysregulation and obesity. How biomaterials influence immune cell recruitment and immune-adipocyte interaction after muscle injury in MRL/MpJ mice is yet to be delineated. To investigate this, we performed volumetric muscle loss surgeries (VML) in C57BL/6 (B6) and MRL/MpJ mice and evaluated immune responses to injury, regenerative (decellularized extracellular matrix, ECM) and fibrotic (polyethylene, PE) material implants using multiparametric flow cytometry, RNA sequencing, and histopathology. In MRL/MpJ but not B6 mice, there was white fat deposition at the site of VML. This was correlated with enrichment of the adipogenesis gene sets including Adipoq, Cebpa, Pparg and Fabp4. In addition, we observed that F4/80- CD68+ macrophages were significantly more abundant in MRL/MpJ mice at 3 weeks post-injury in PE and control groups. Furthermore, in MRL/MpJ mice, the lipid associated macrophage gene signature comprising Trem2, Ly6c2, Cd9, Cd63, Lyz2 were highly potentiated by muscle injury as well as PE implantation. In contrast, ECM implantation produced a largely SiglecF+ eosinophilic response (∼60% of CD45+ cells) and partially suppressed CD68+ macrophage abundance. Furthermore, ECM potentiated expression of the pro-regenerative Arg1, Chil3, Retnla and Gata3 in both B6 and MRL/MpJ. In PE groups, Ly6G+ neutrophils were the second most abundant cell type (∼ 25% of CD45+ cells) which was corroborated by increased abundance of Vav1, Ccl2, Adam8 and Rac2. Taken together, our results suggest a potential mechanistic role for CD68+ macrophages in promoting adipogenesis after VML.

## 2 Read mapping from raw fastq files using kallisto

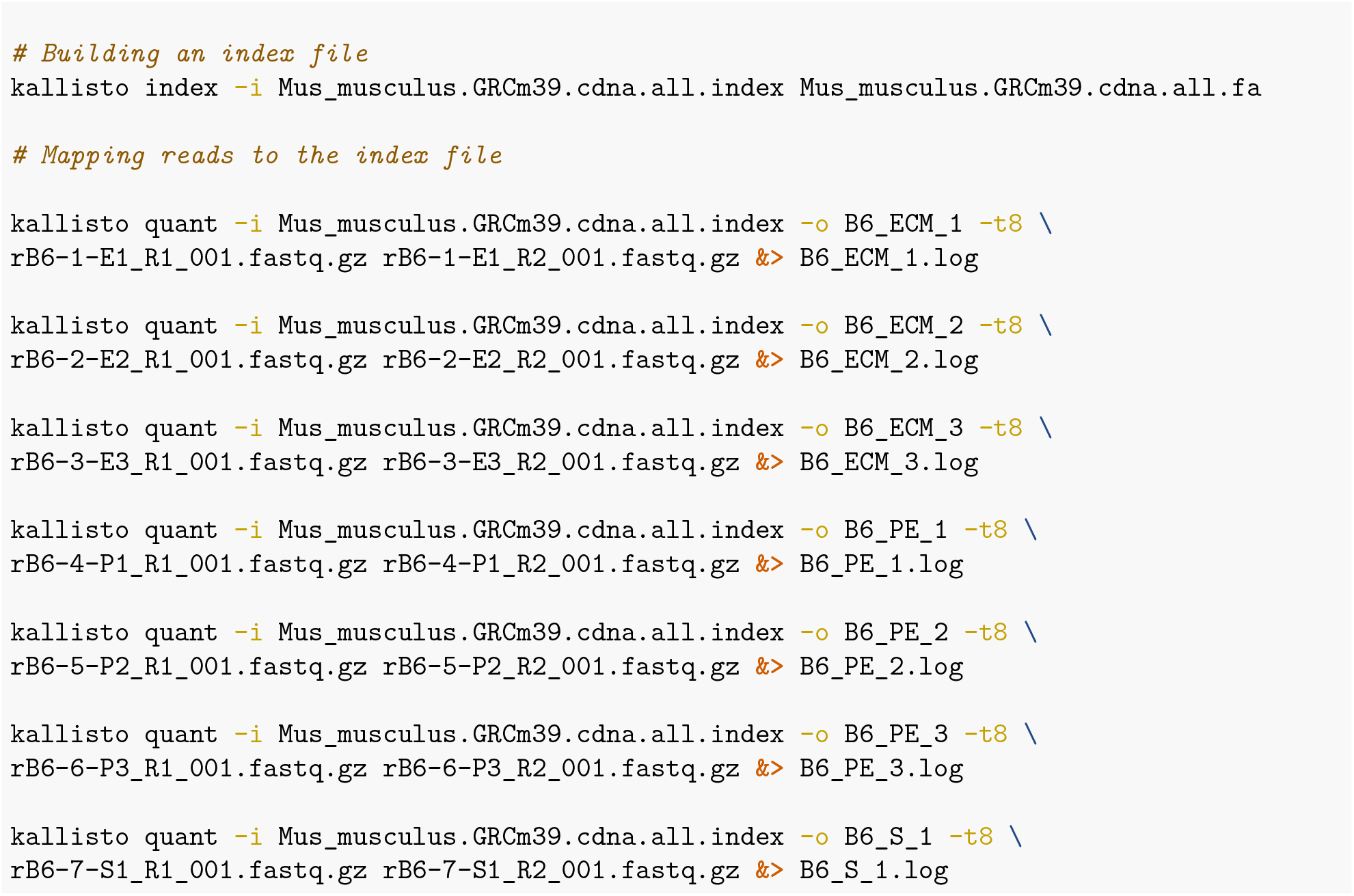

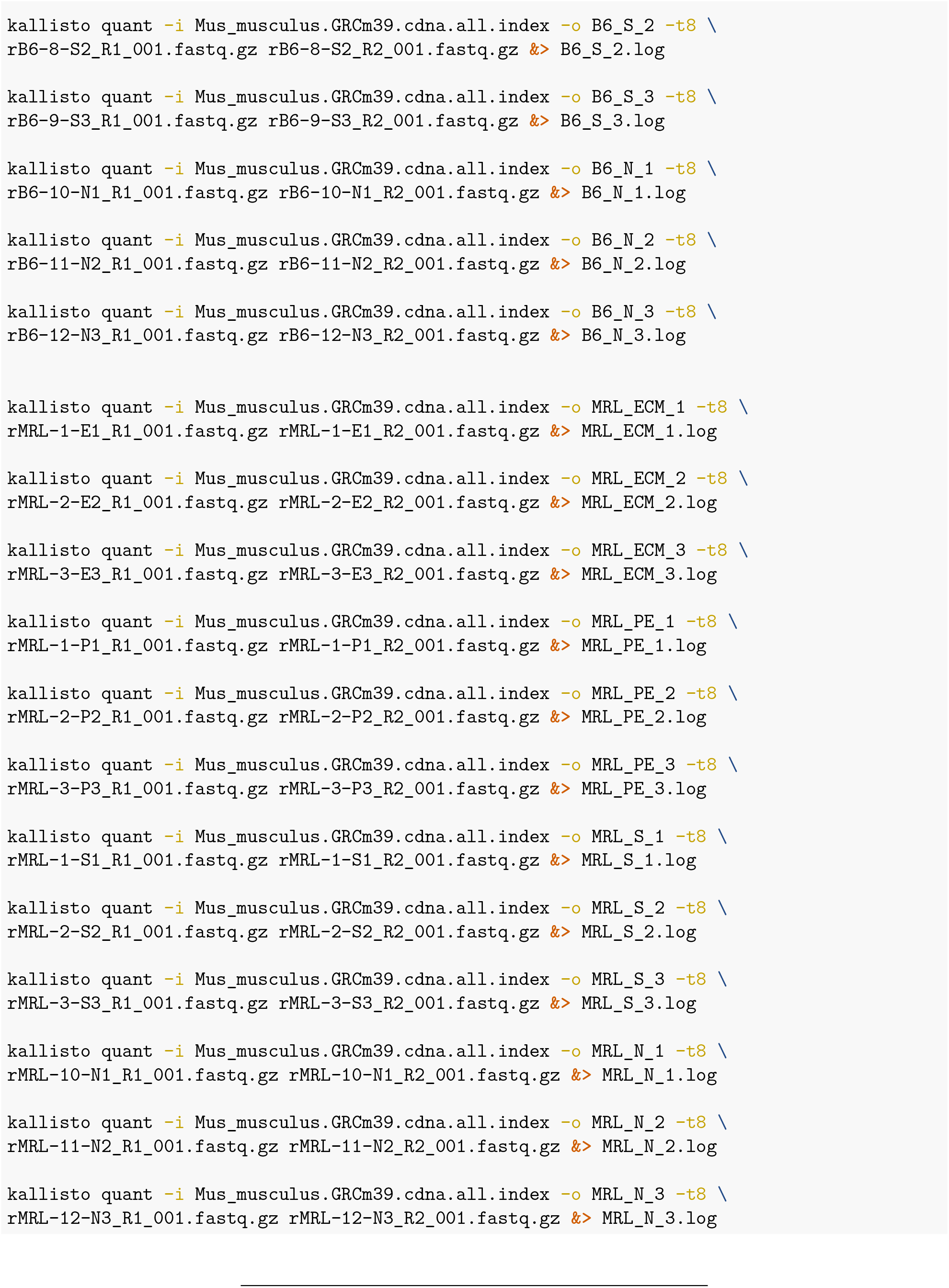

## 3 R packages used for analysis

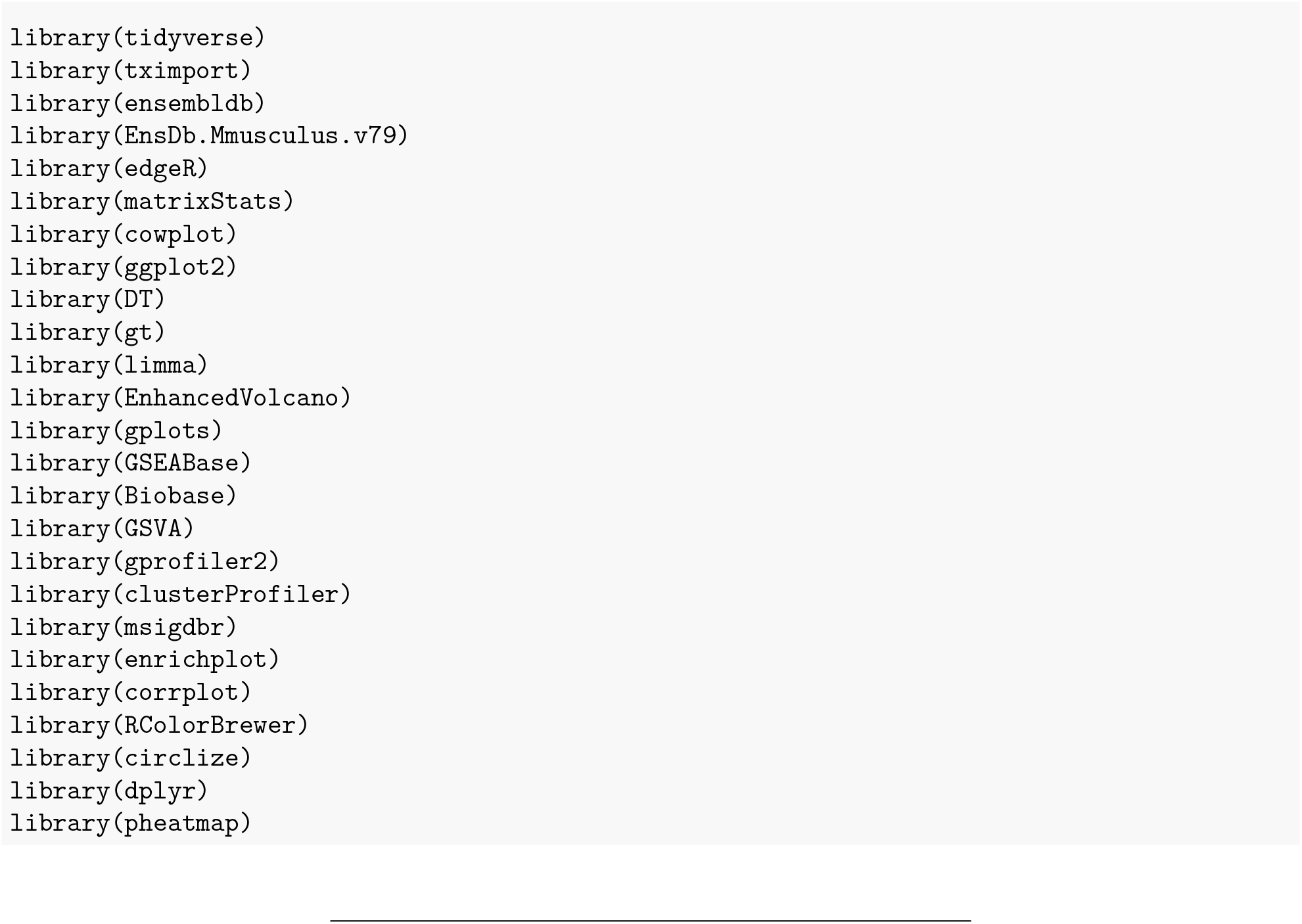

## 4 Gene Annotation

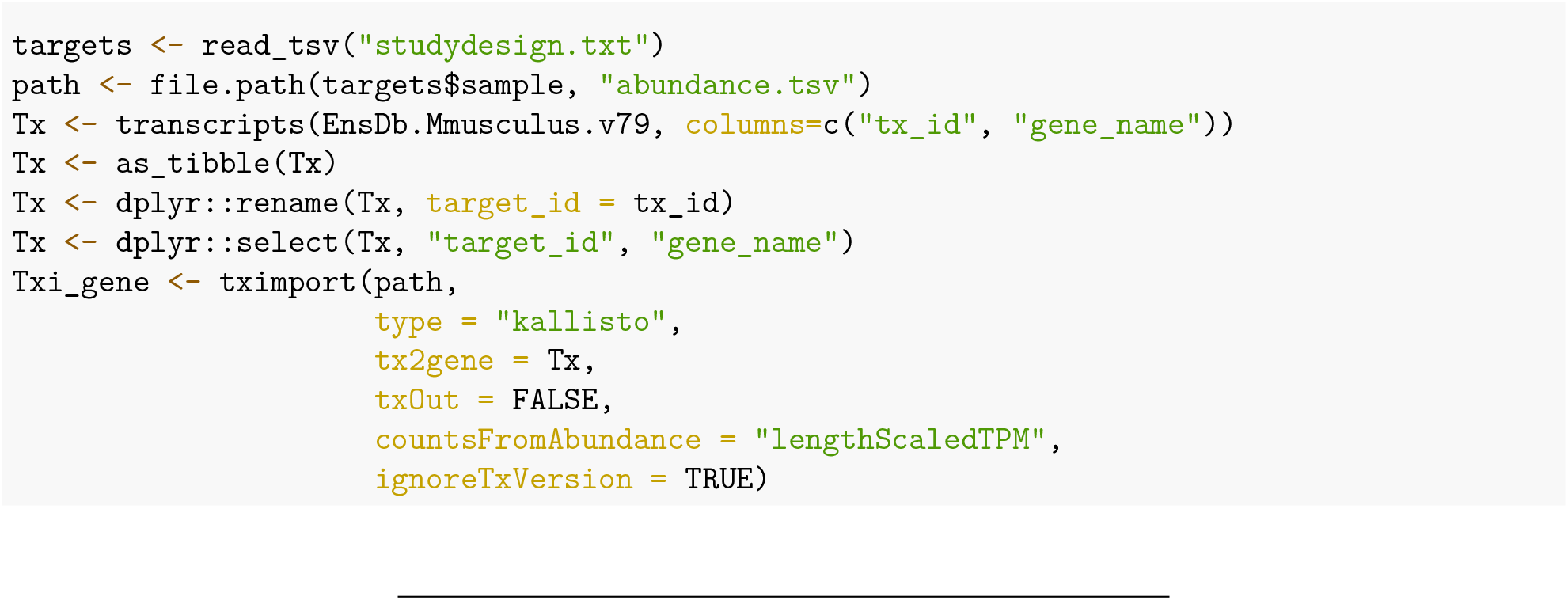

## 5 Data filtering and normalization

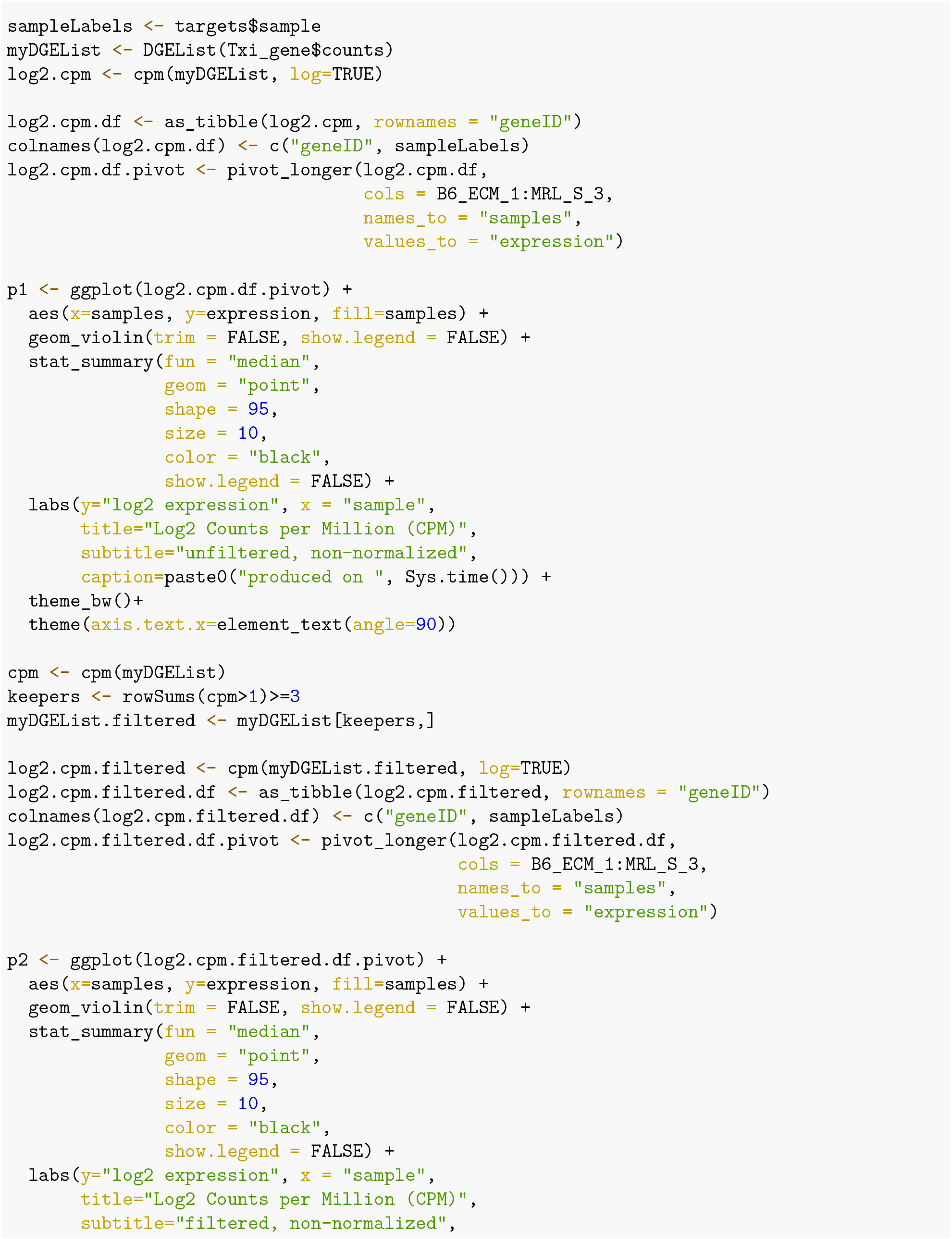

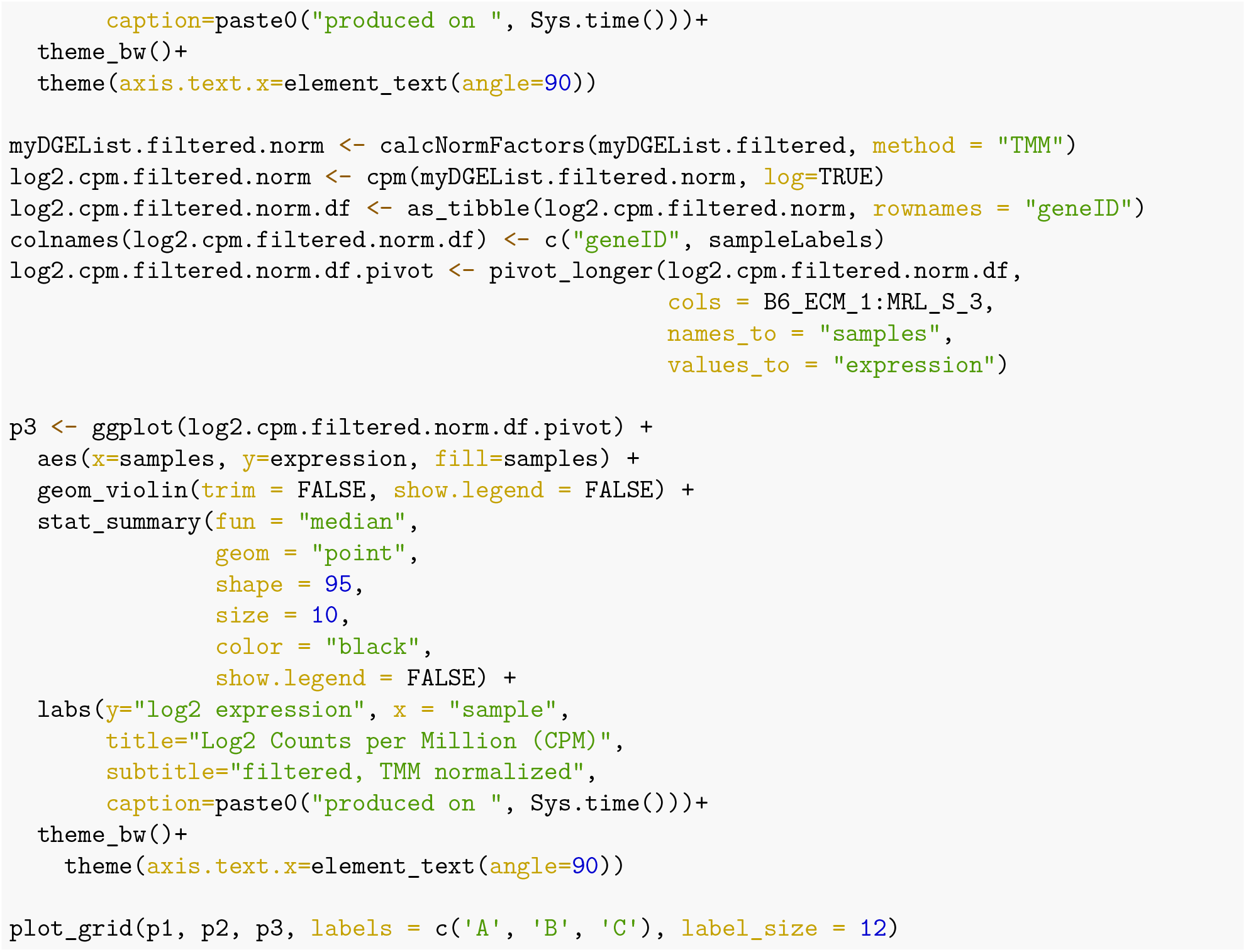

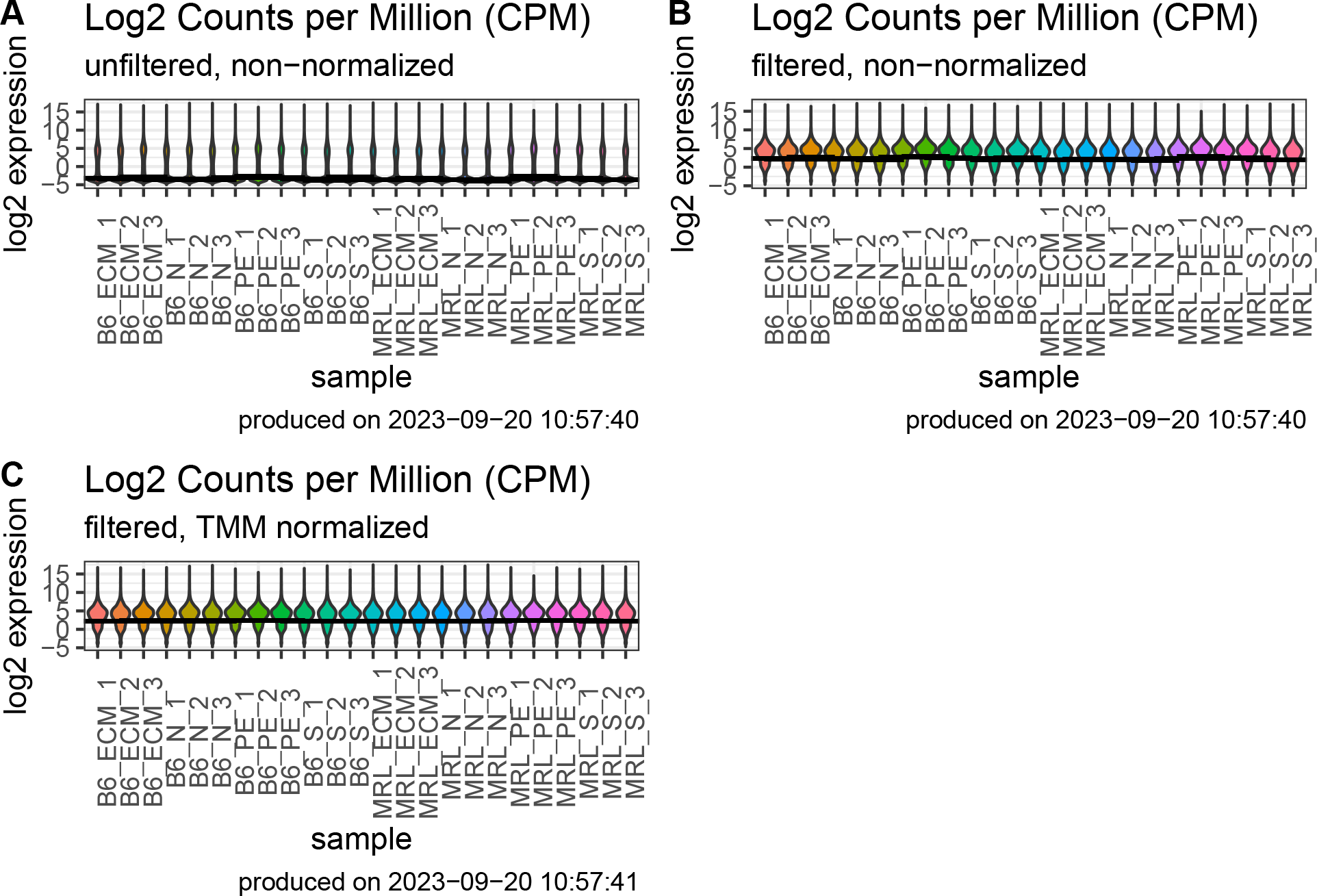

## 6 Data Analysis

### 6.1 PCA plot

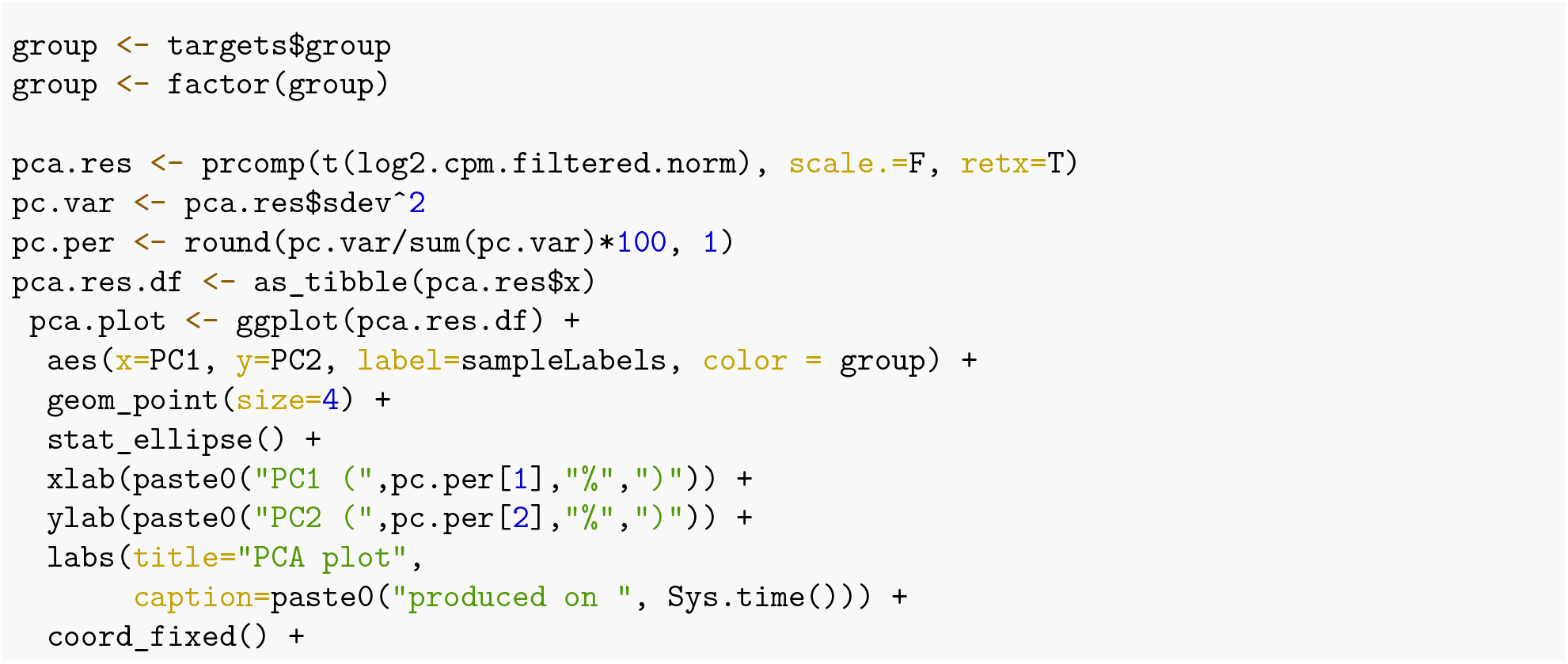

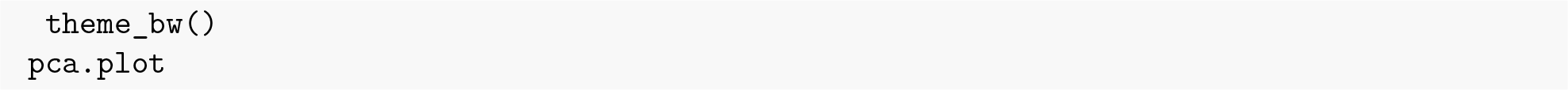

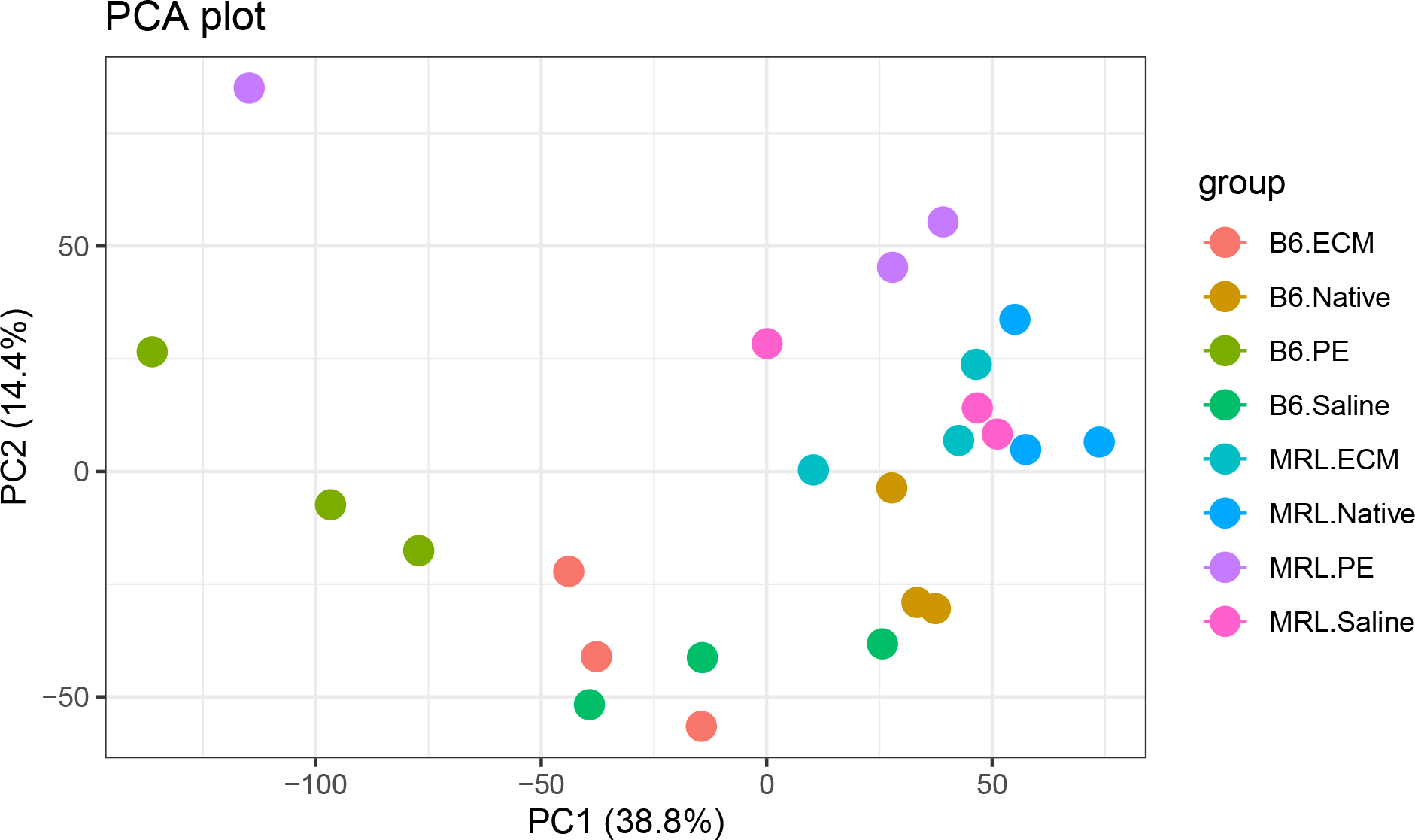

### 6.2 Volcano plot-Differential expression analysis: B6 vs MRL Saline

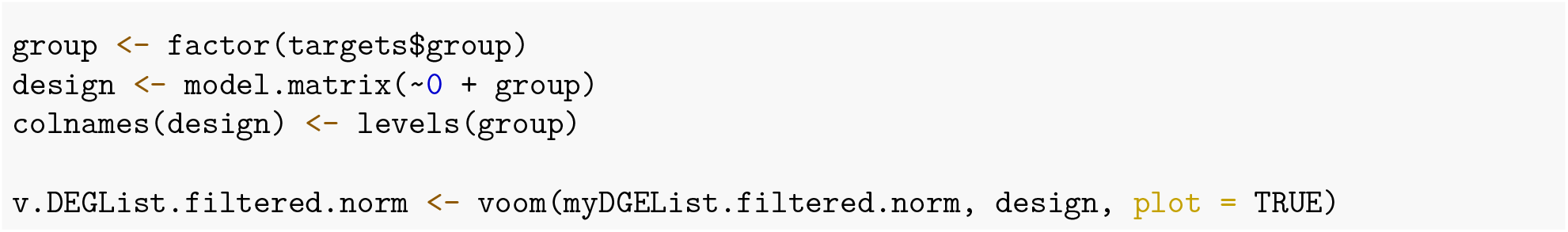

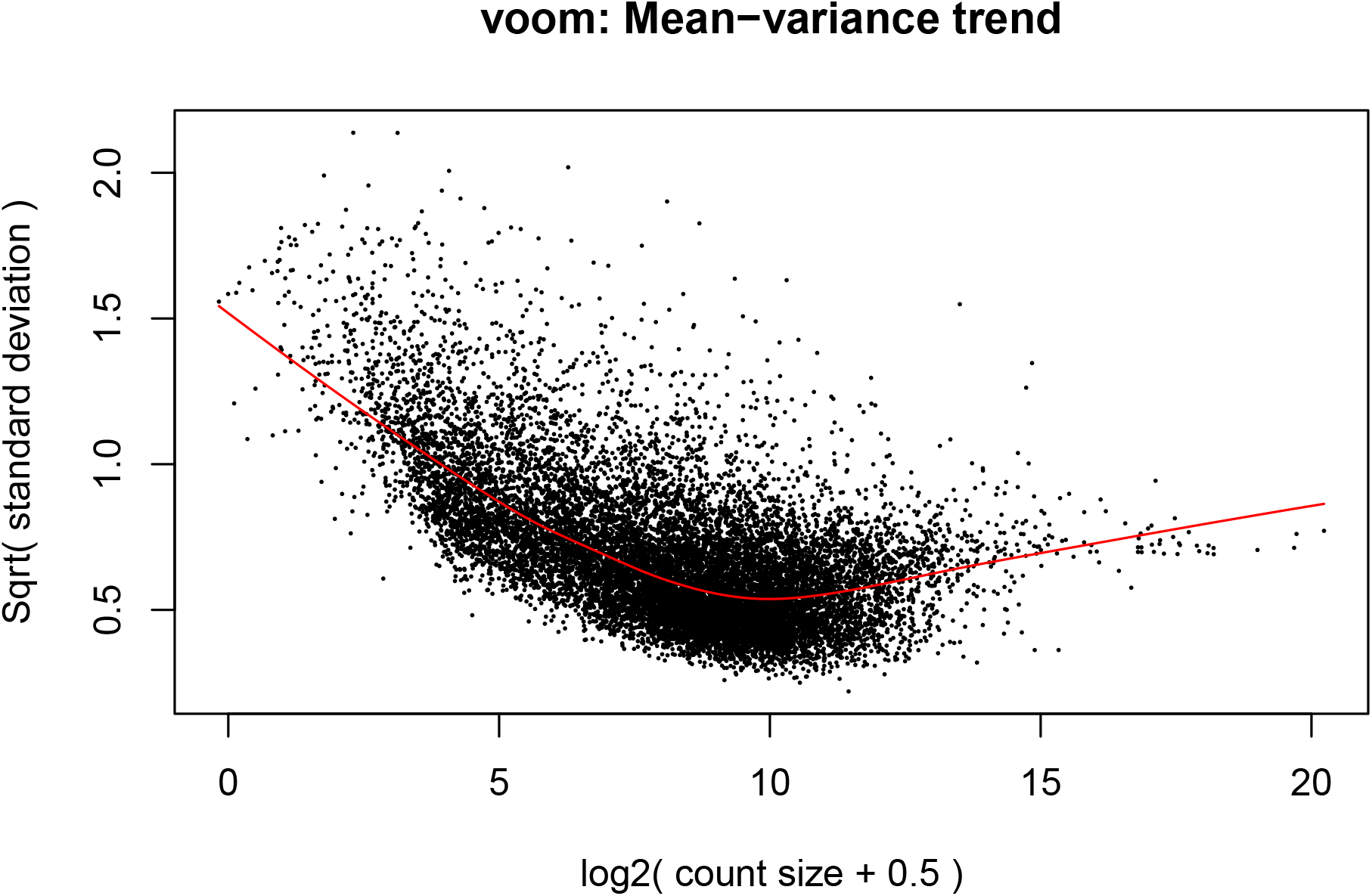

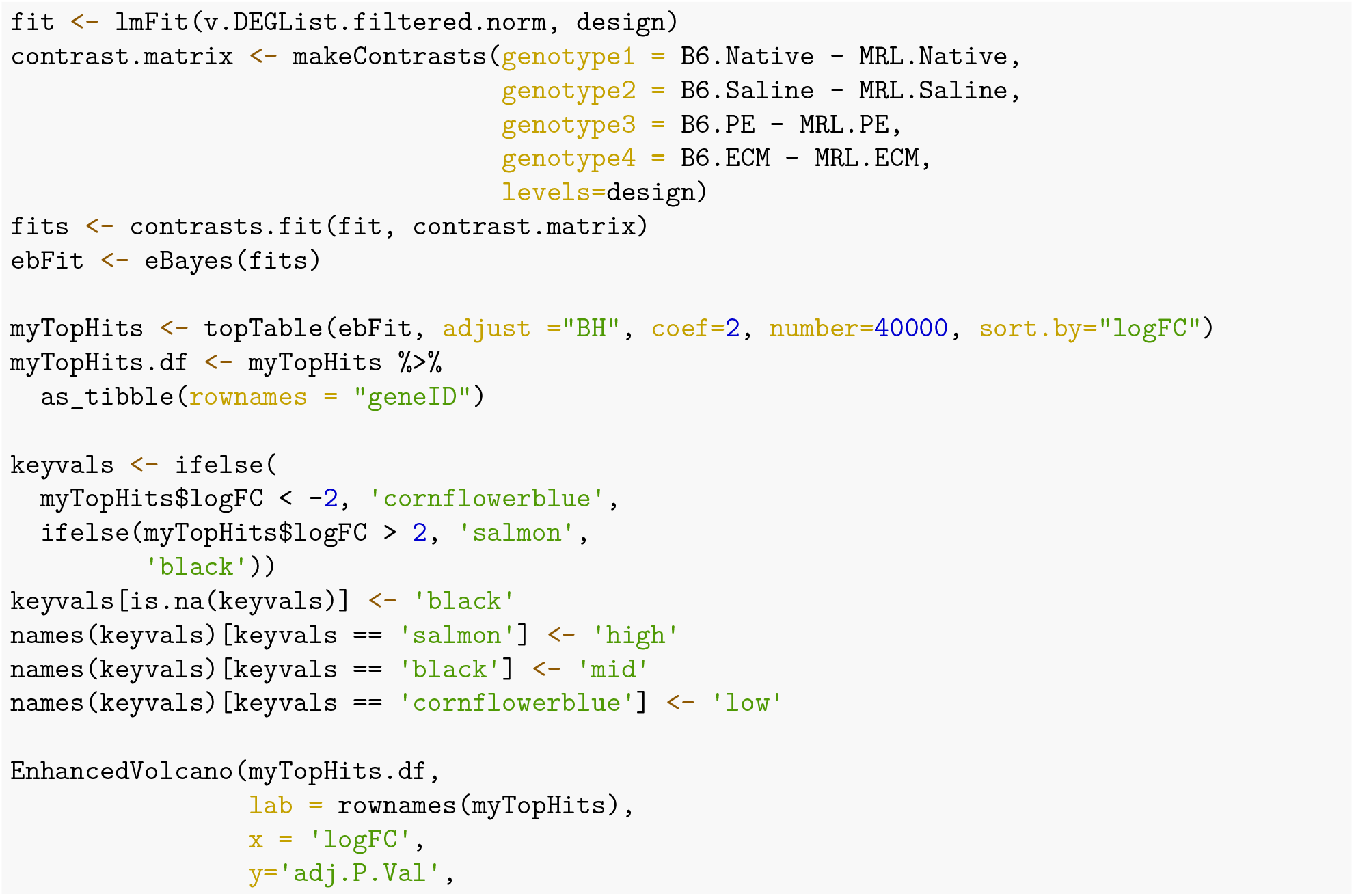

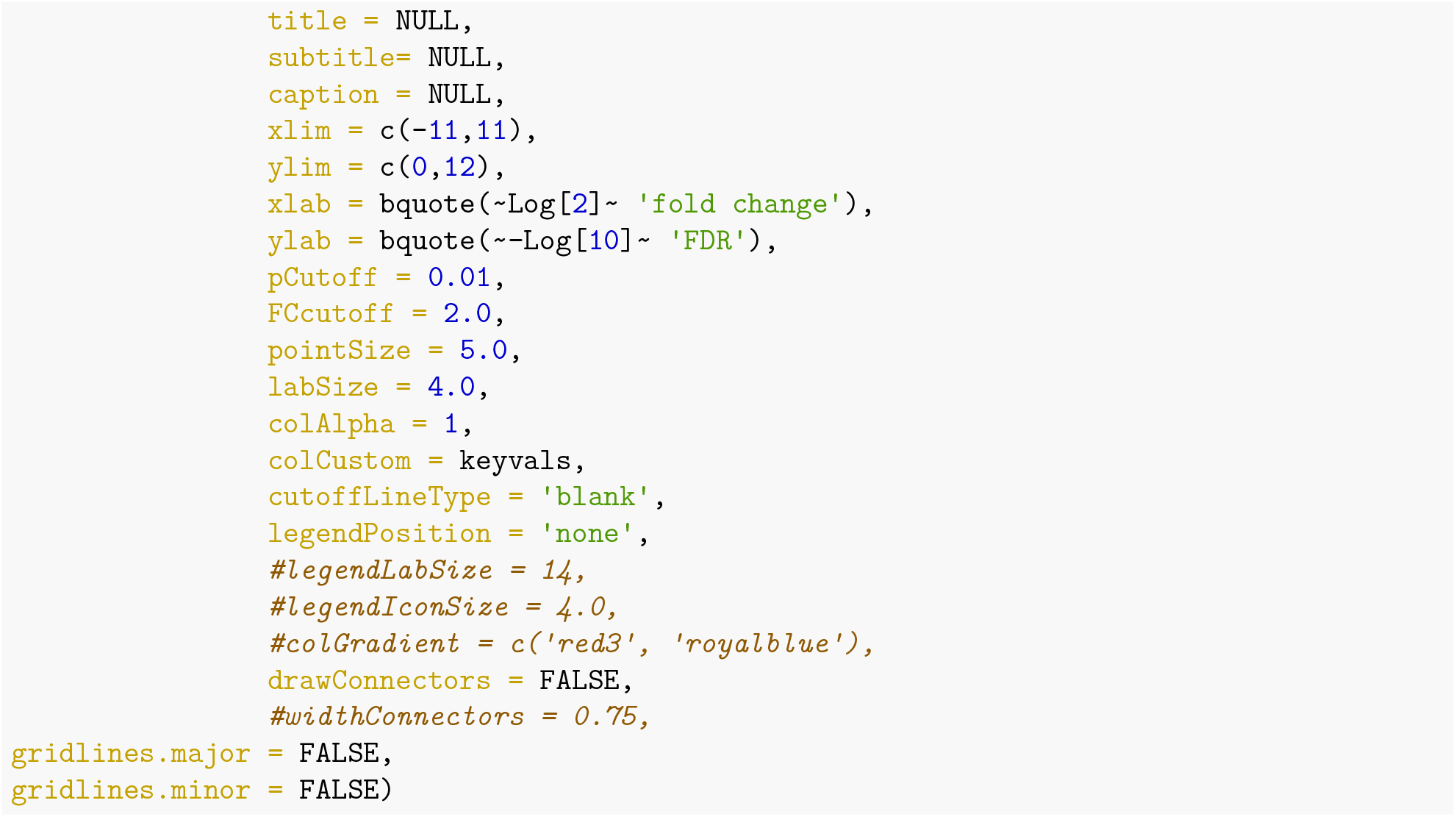

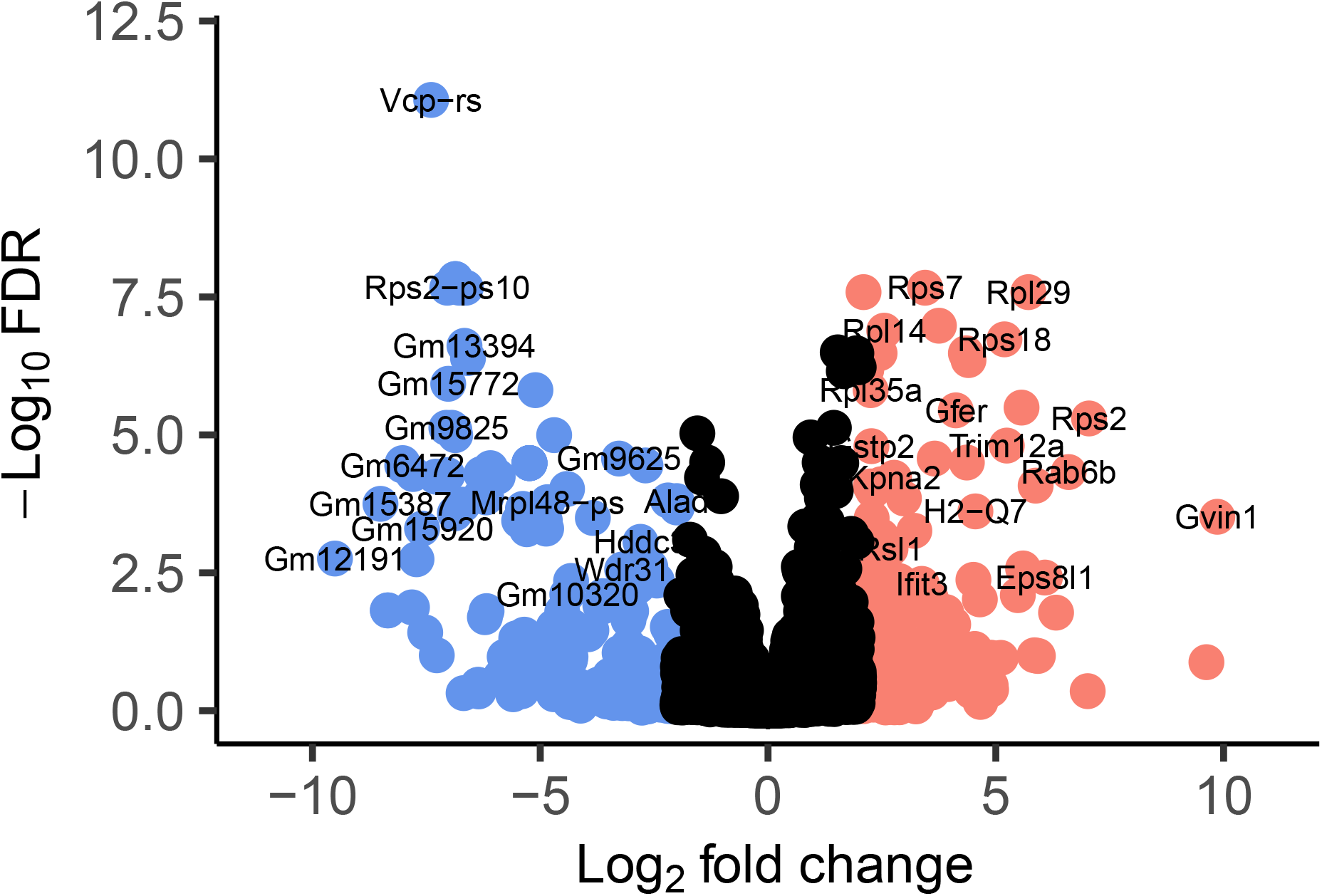

### 6.3 Heatmap: Top 50 differentially expressed genes-B6 vs MRL Uninjured

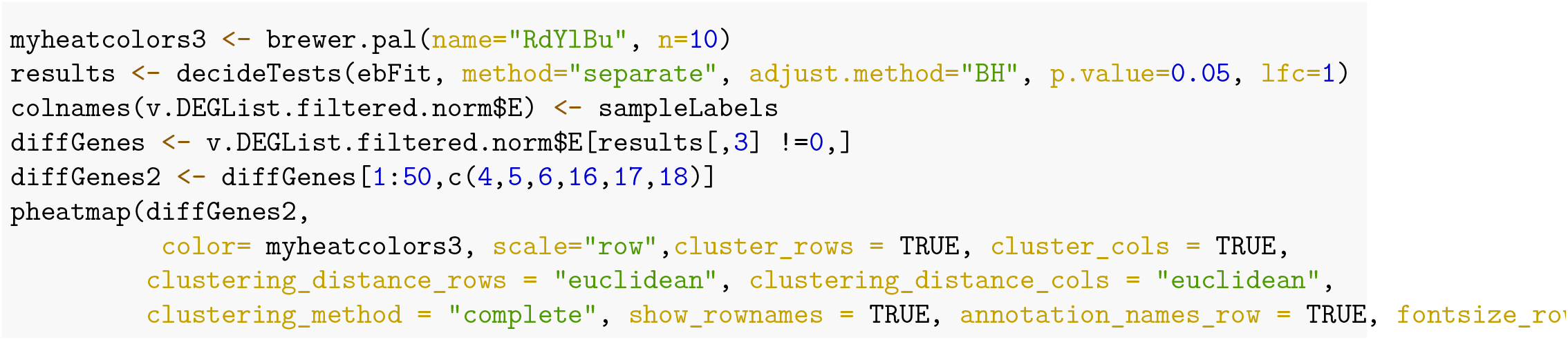

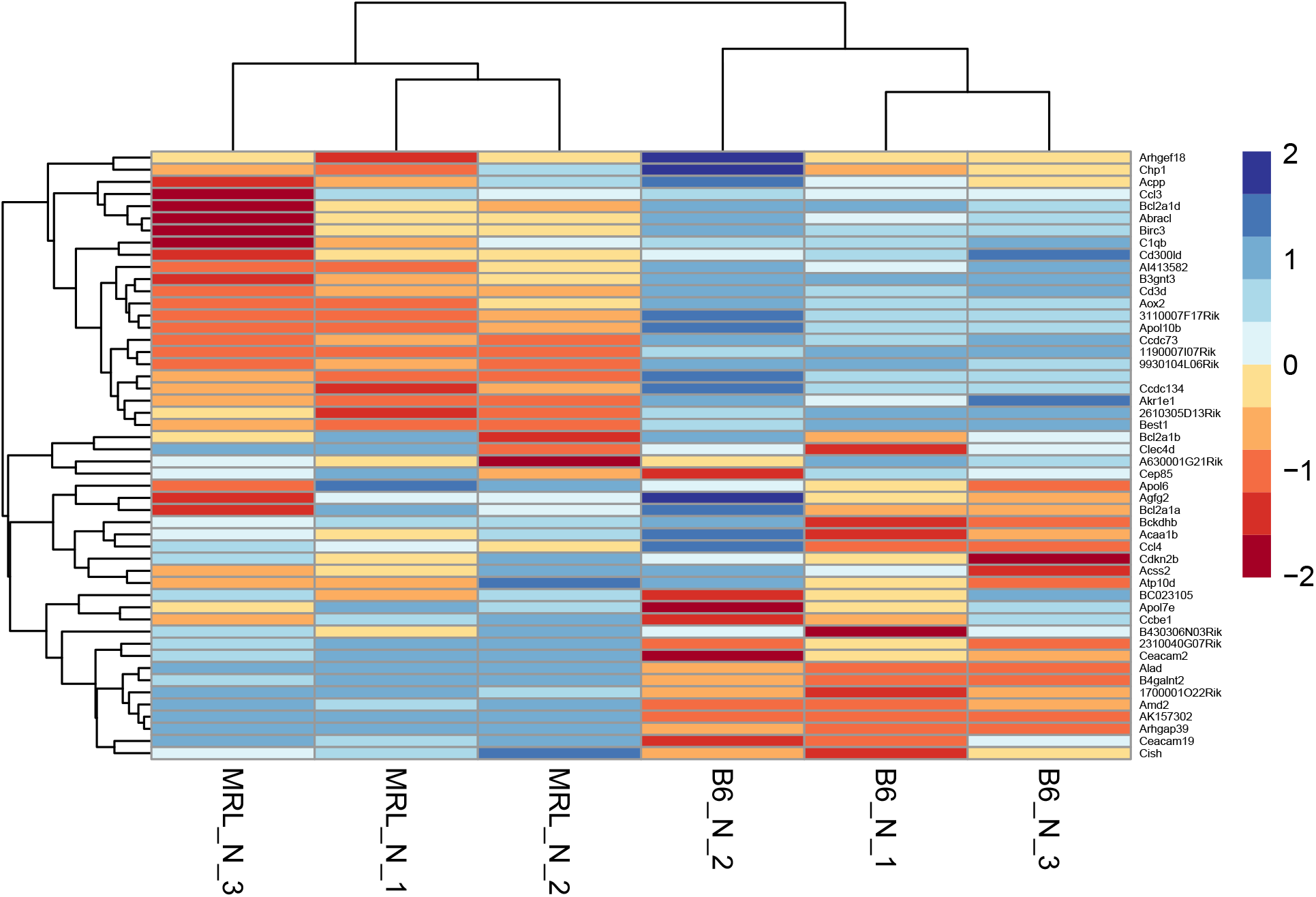

### 6.4 Gene Ontology and Pathway Assignment: Saline

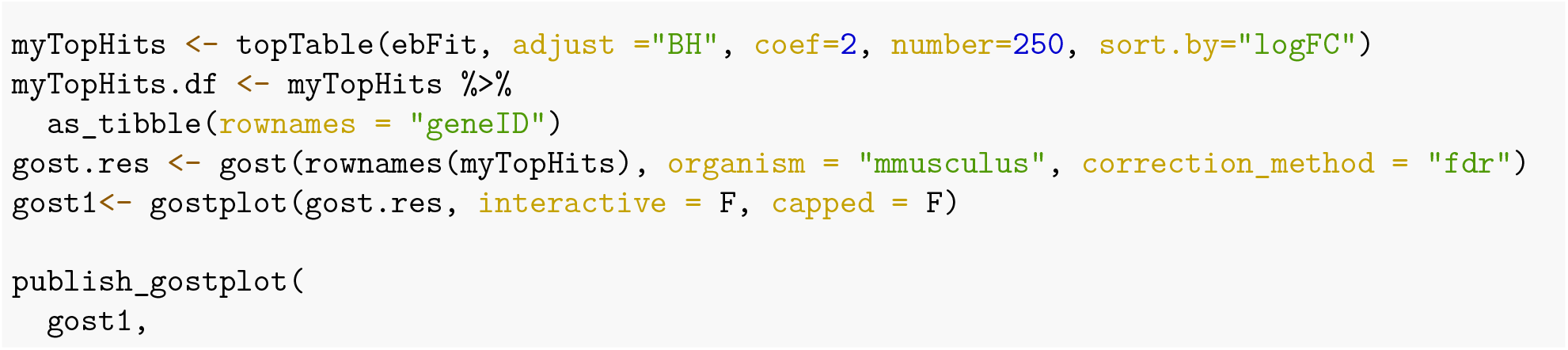

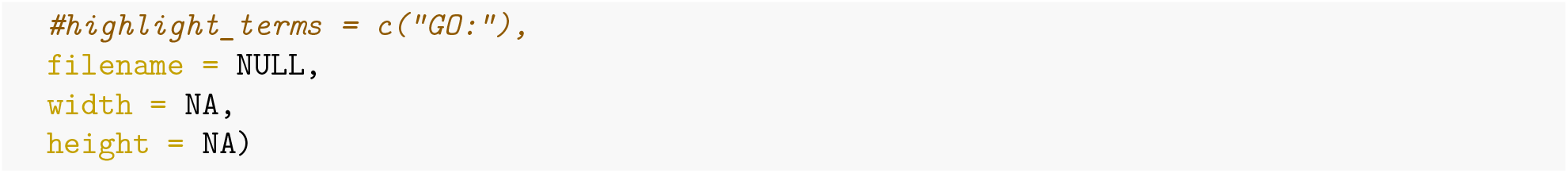

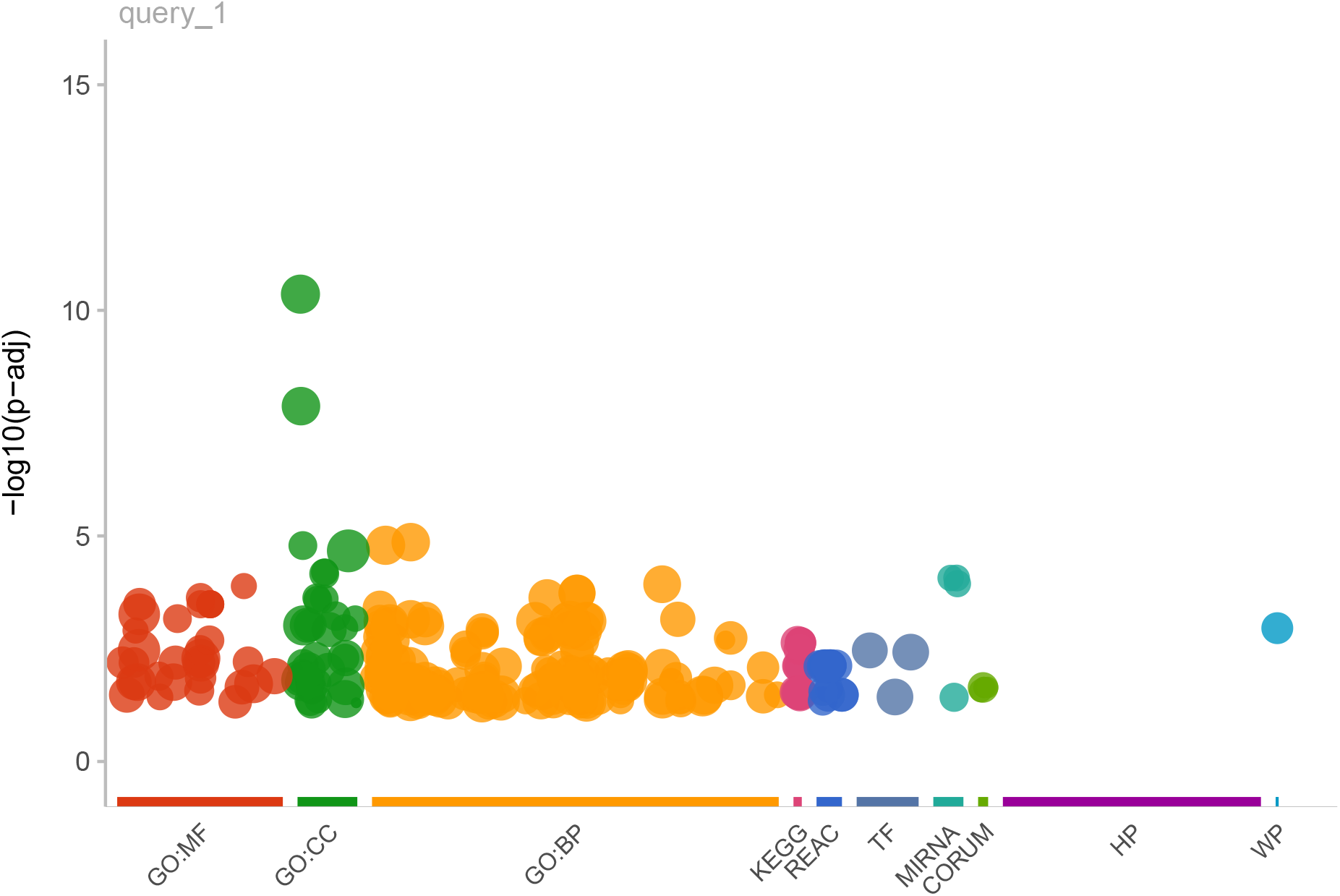

### 6.5 Gene Set Enrichment Analysis-B6 vs MRL: Saline

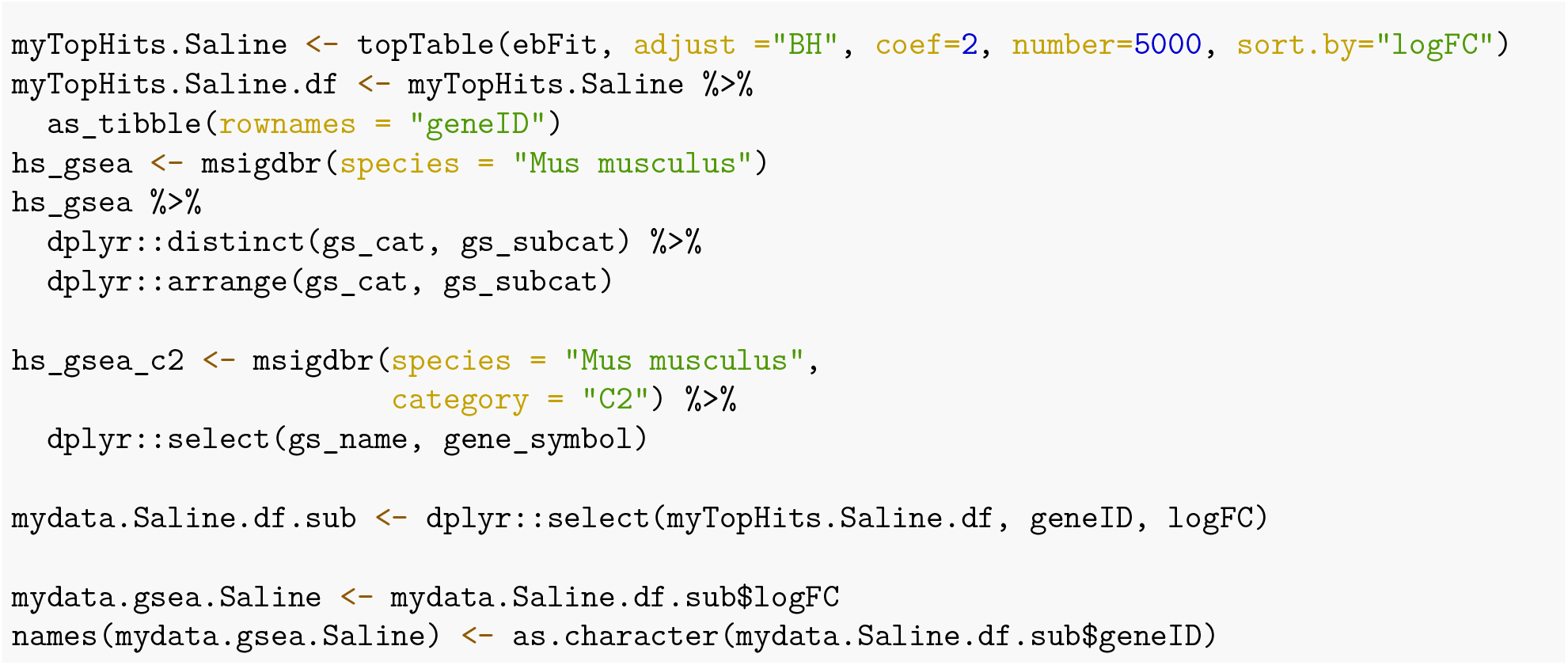

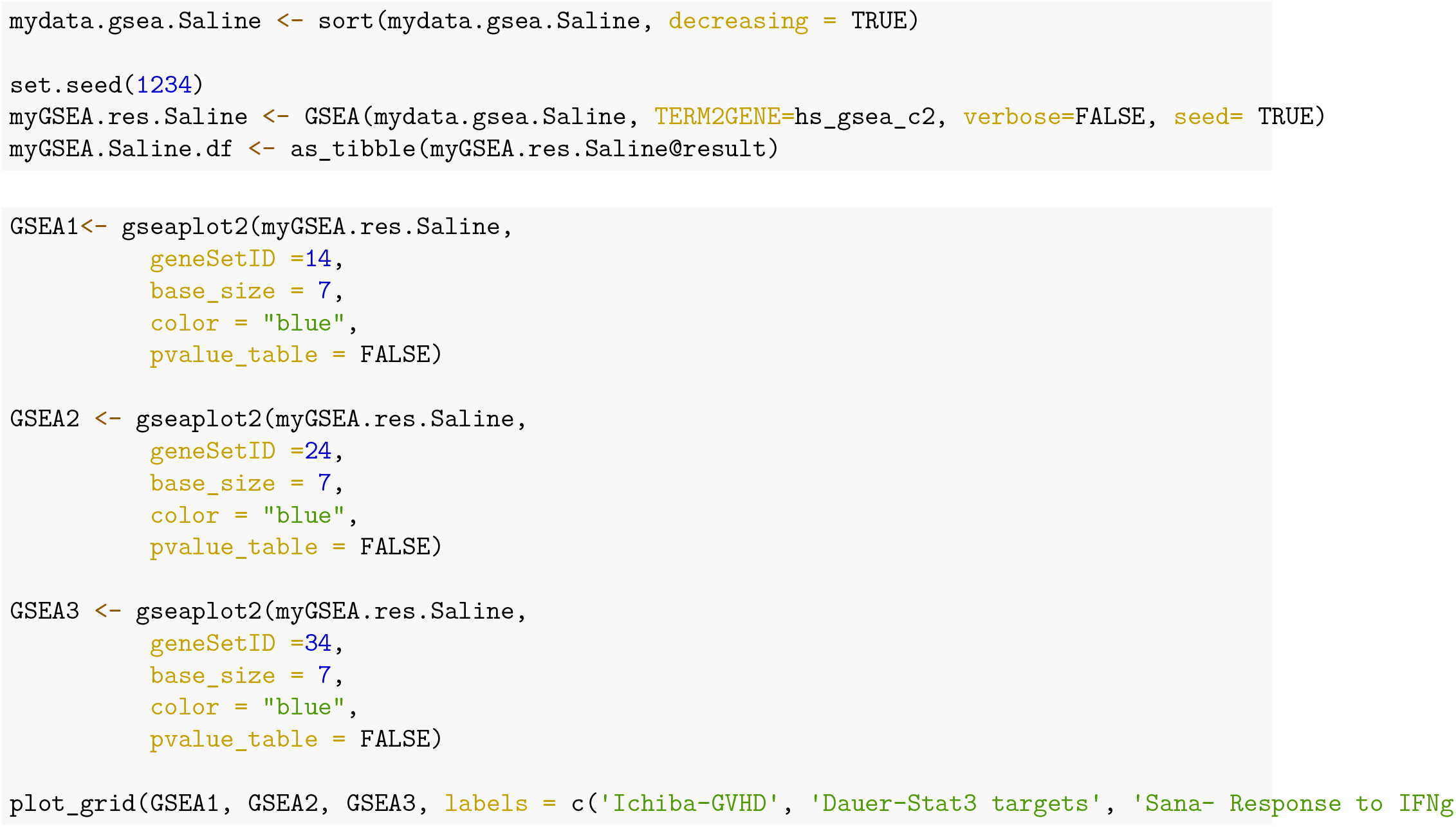

### 6.6 Gene Set Enrichment Analysis-B6 vs MRL: PE

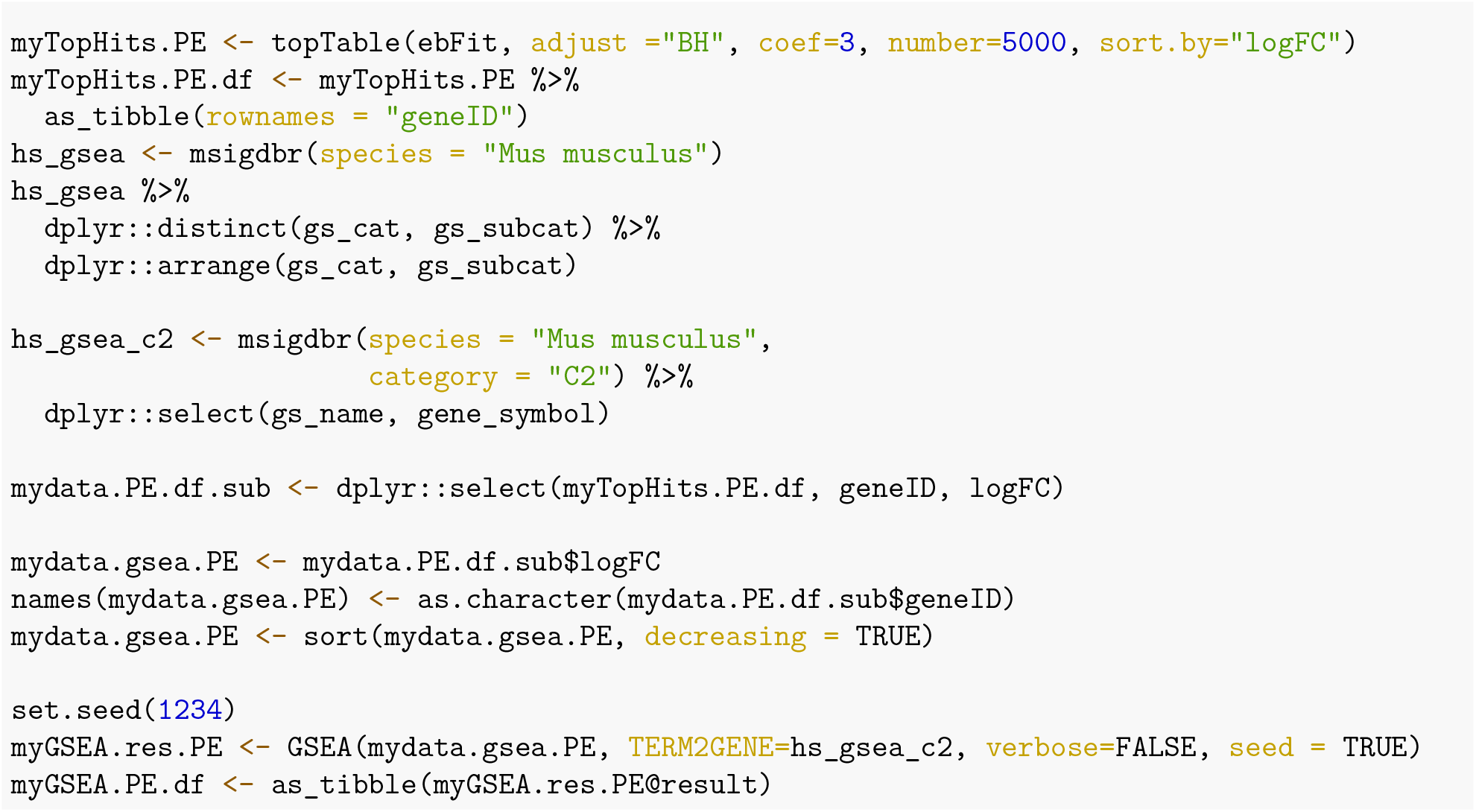

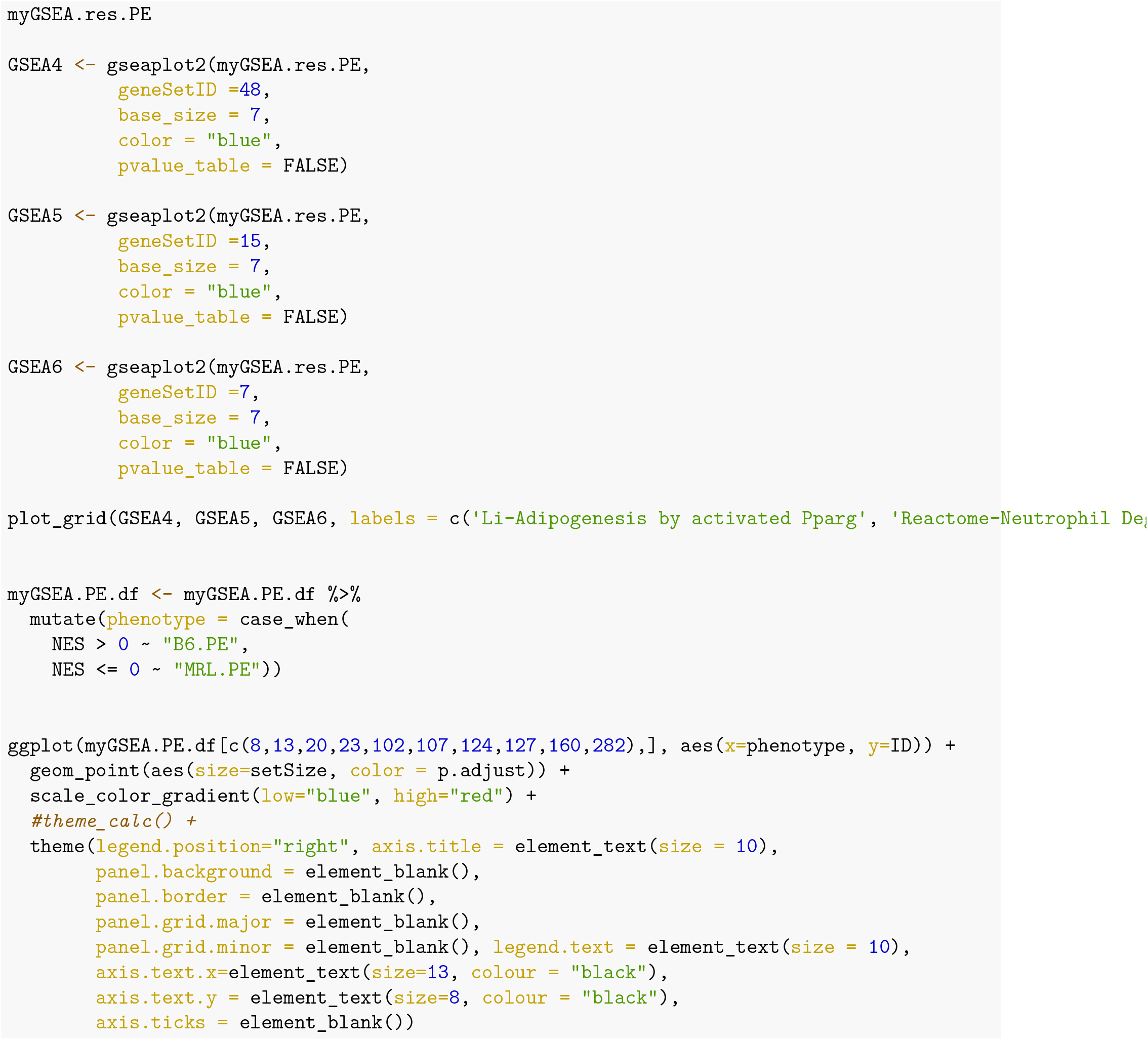

### 6.7 ImmQuant scores and gene signatures correlation plot: PE

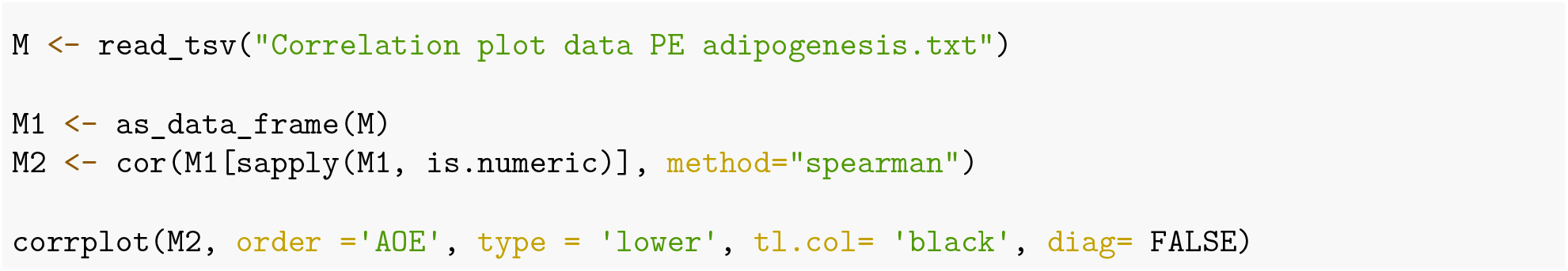

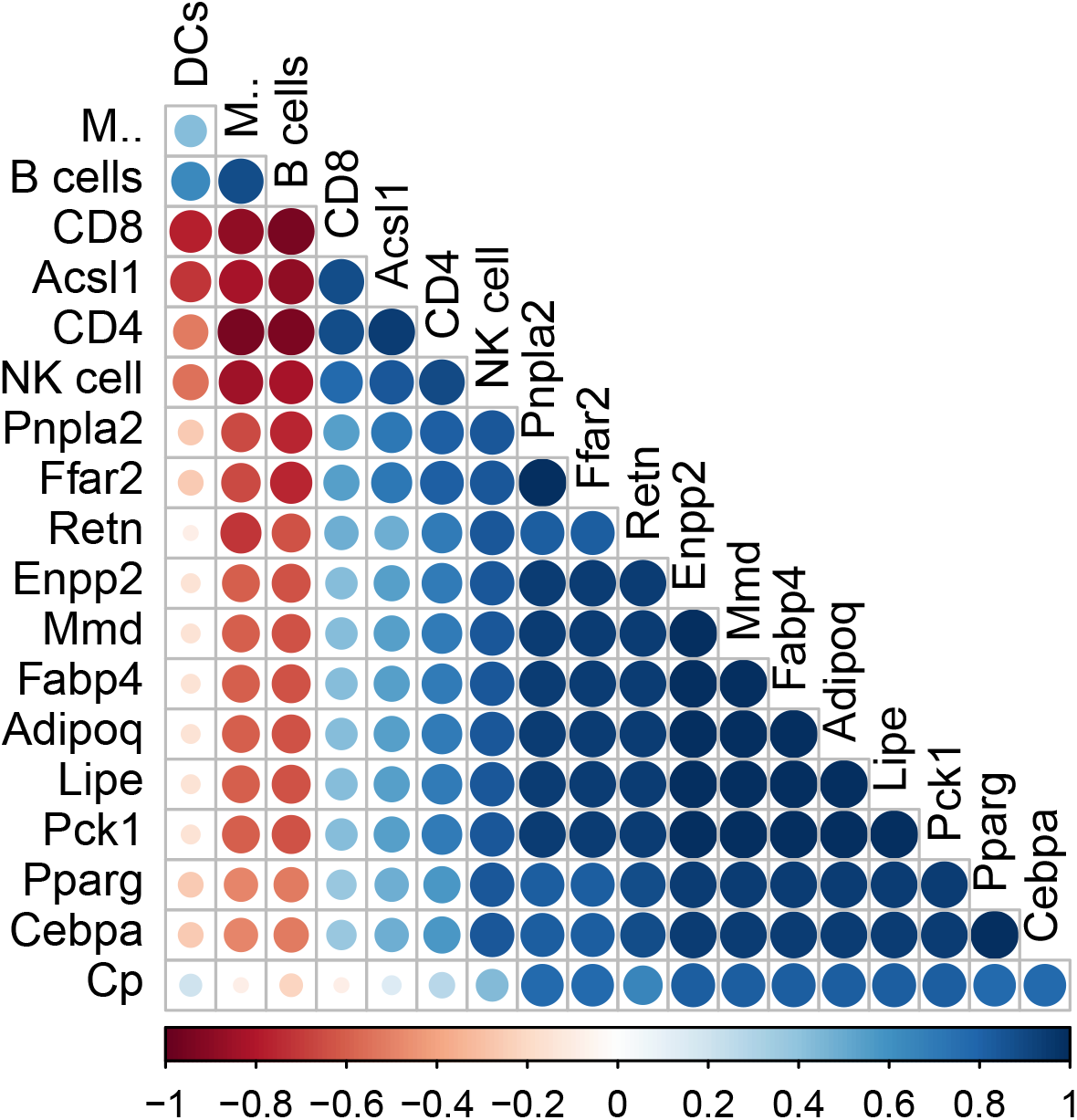

### 6.8 Heatmap of differentially expressed genes: ECM vs PE

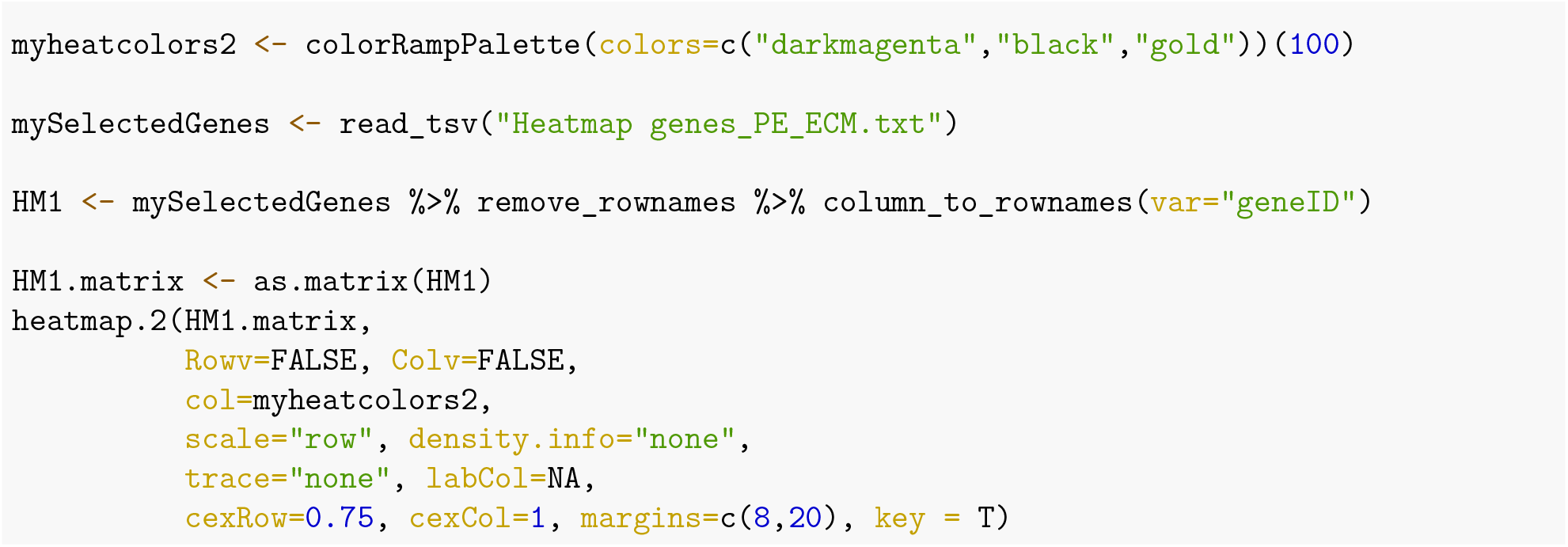

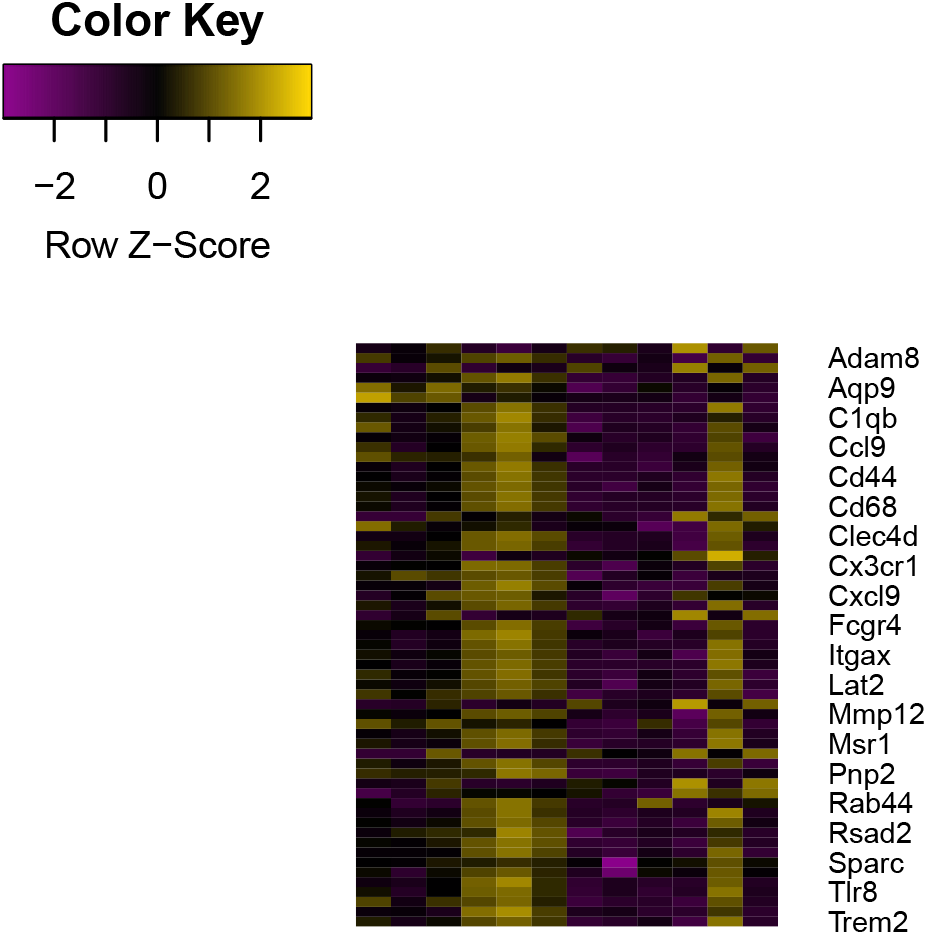

### 6.9 Gene Set Enrichment Analysis-B6 vs MRL: ECM

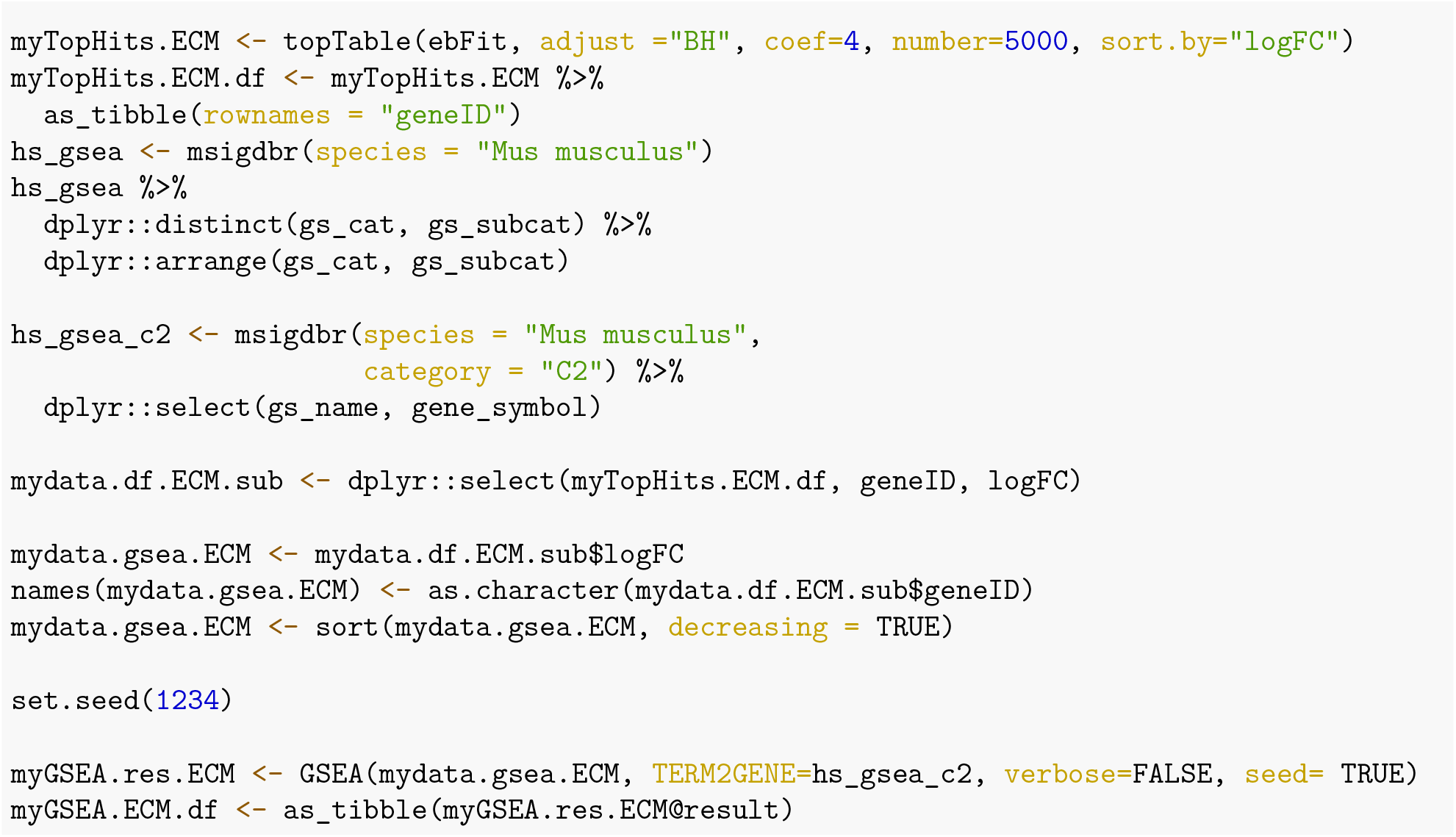

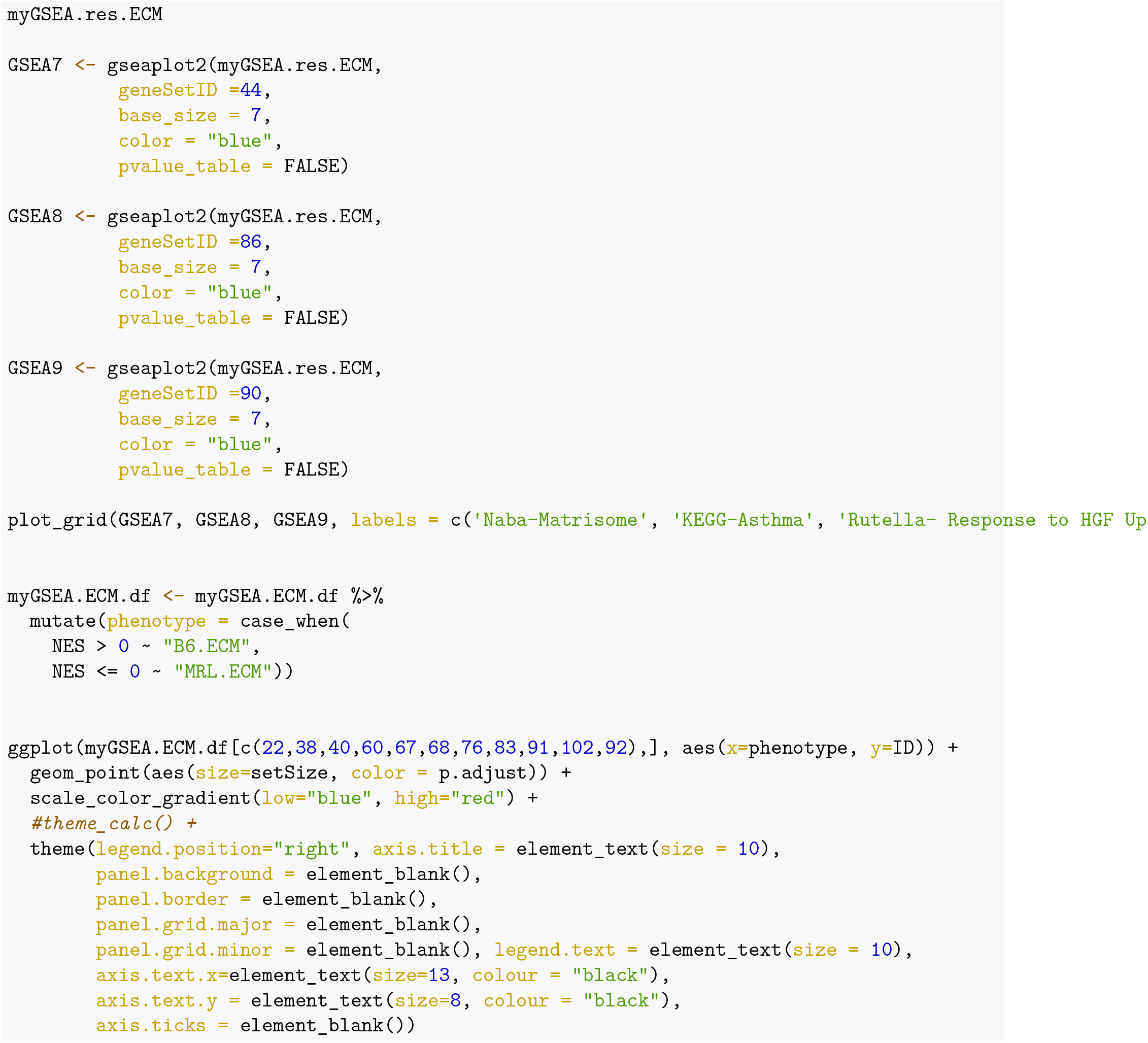

### 6.10 ImmQuant scores and gene signatures correlation plot: ECM

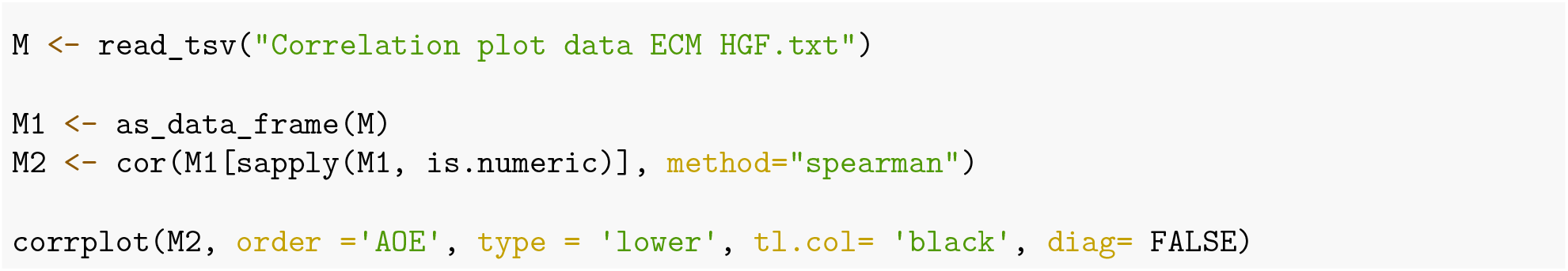

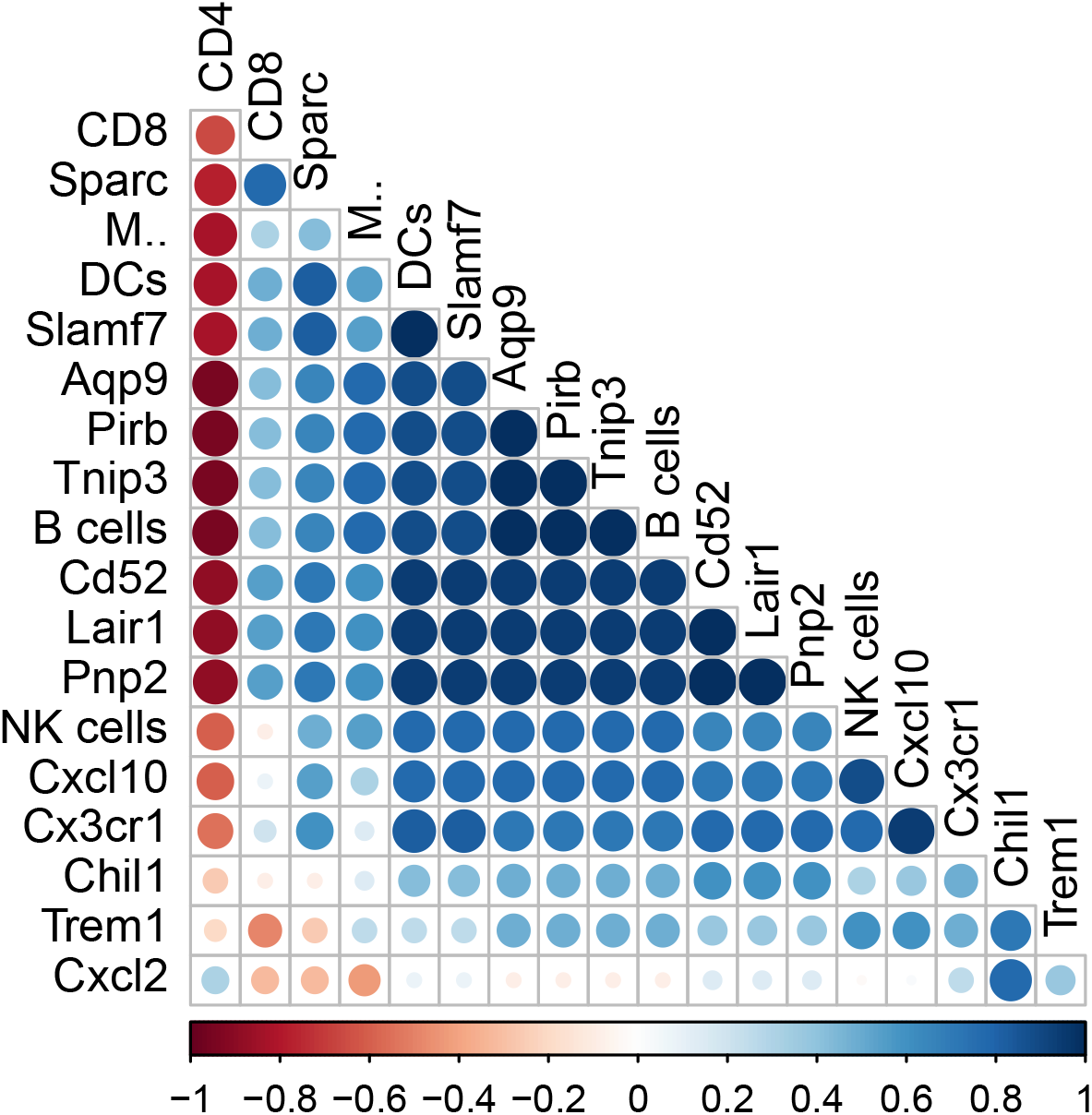

### 6.11 Interaction plot-PE: Immune Cell Targets and Adipogenesis

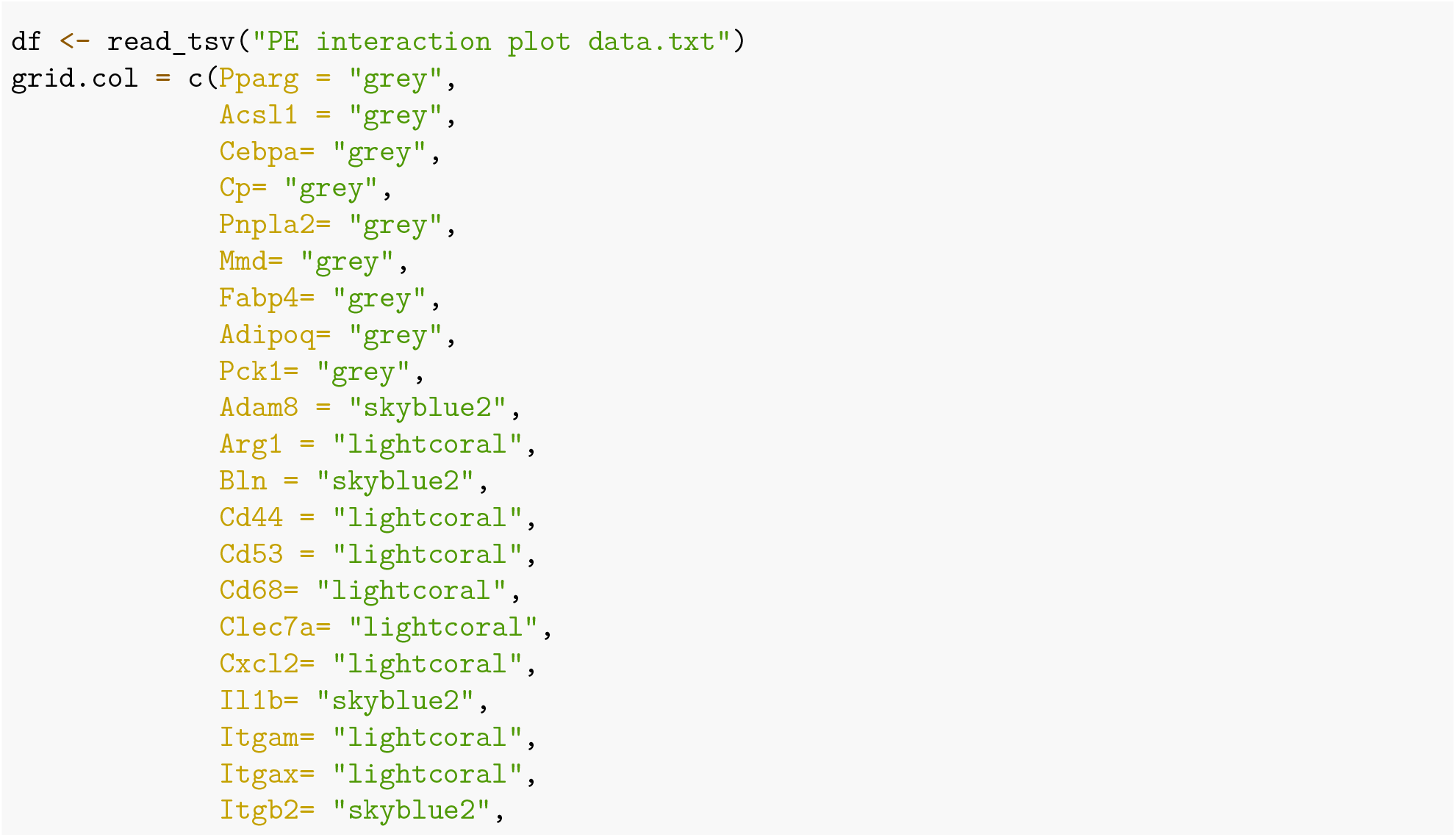

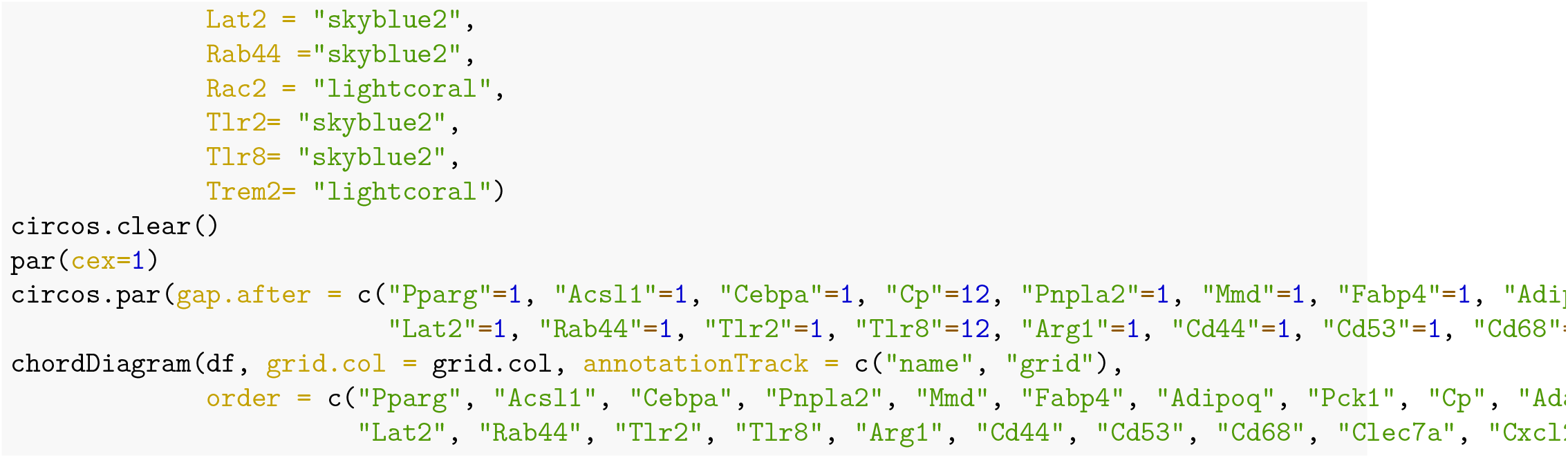

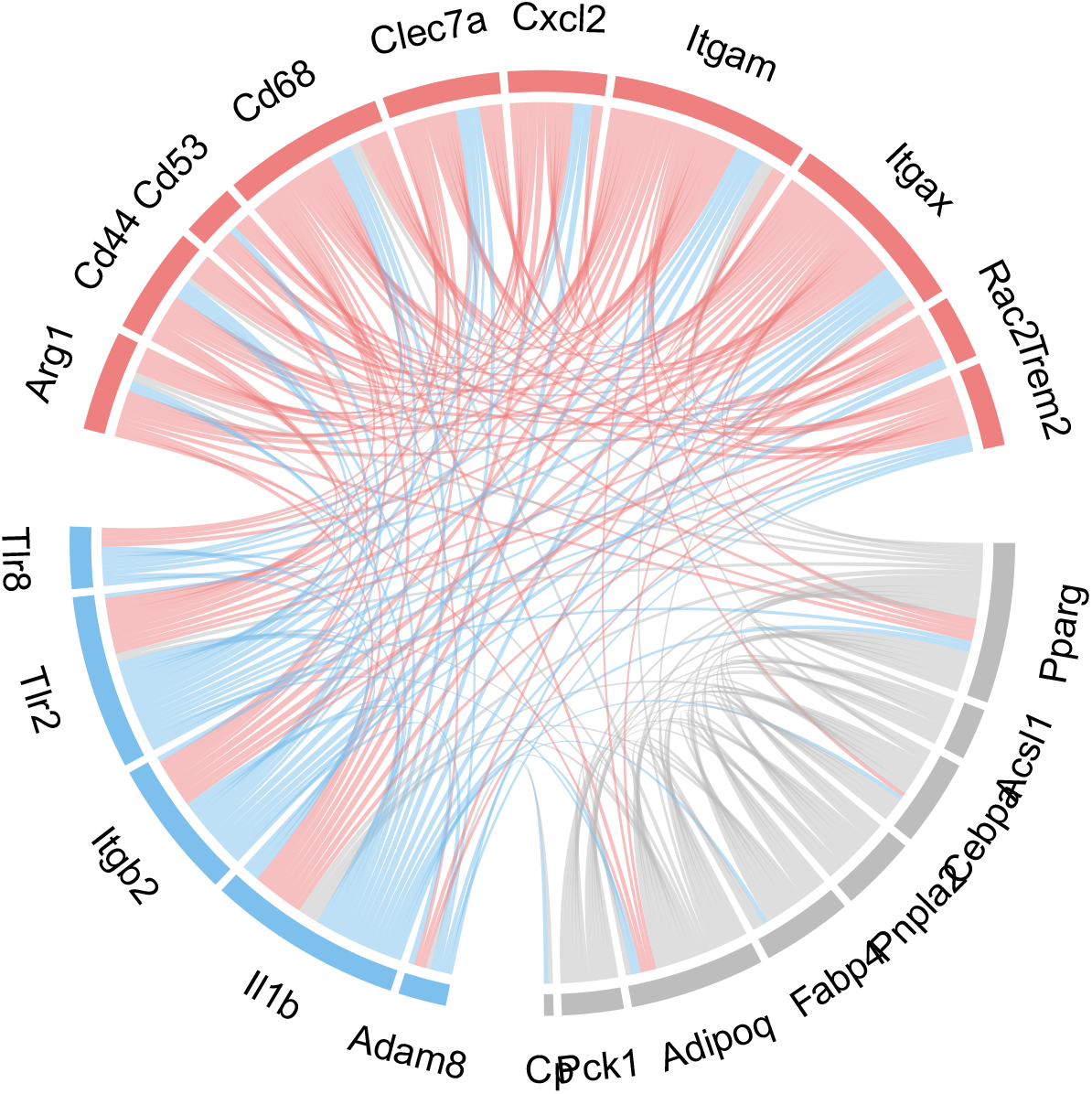

### 6.12 Interaction plot-ECM:HGF response and Eosinophil gene set

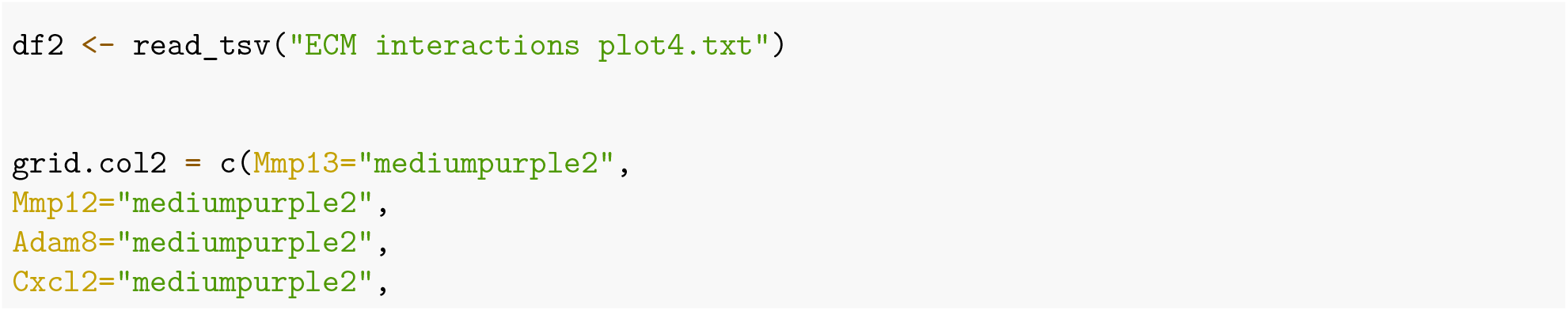

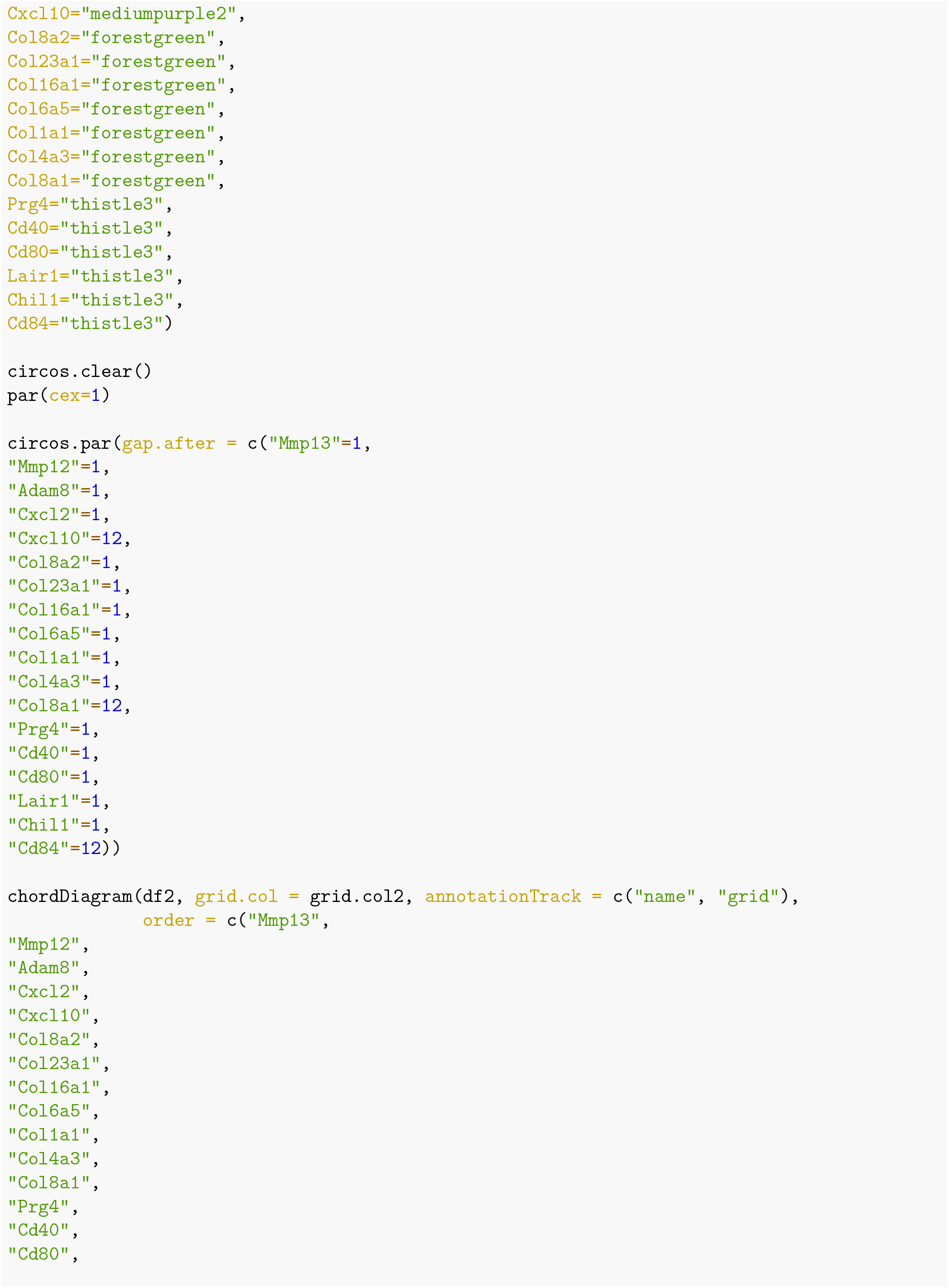

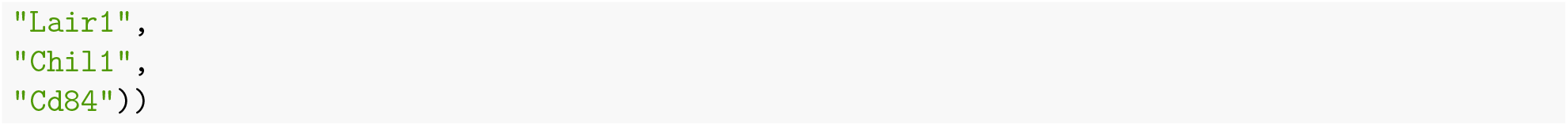

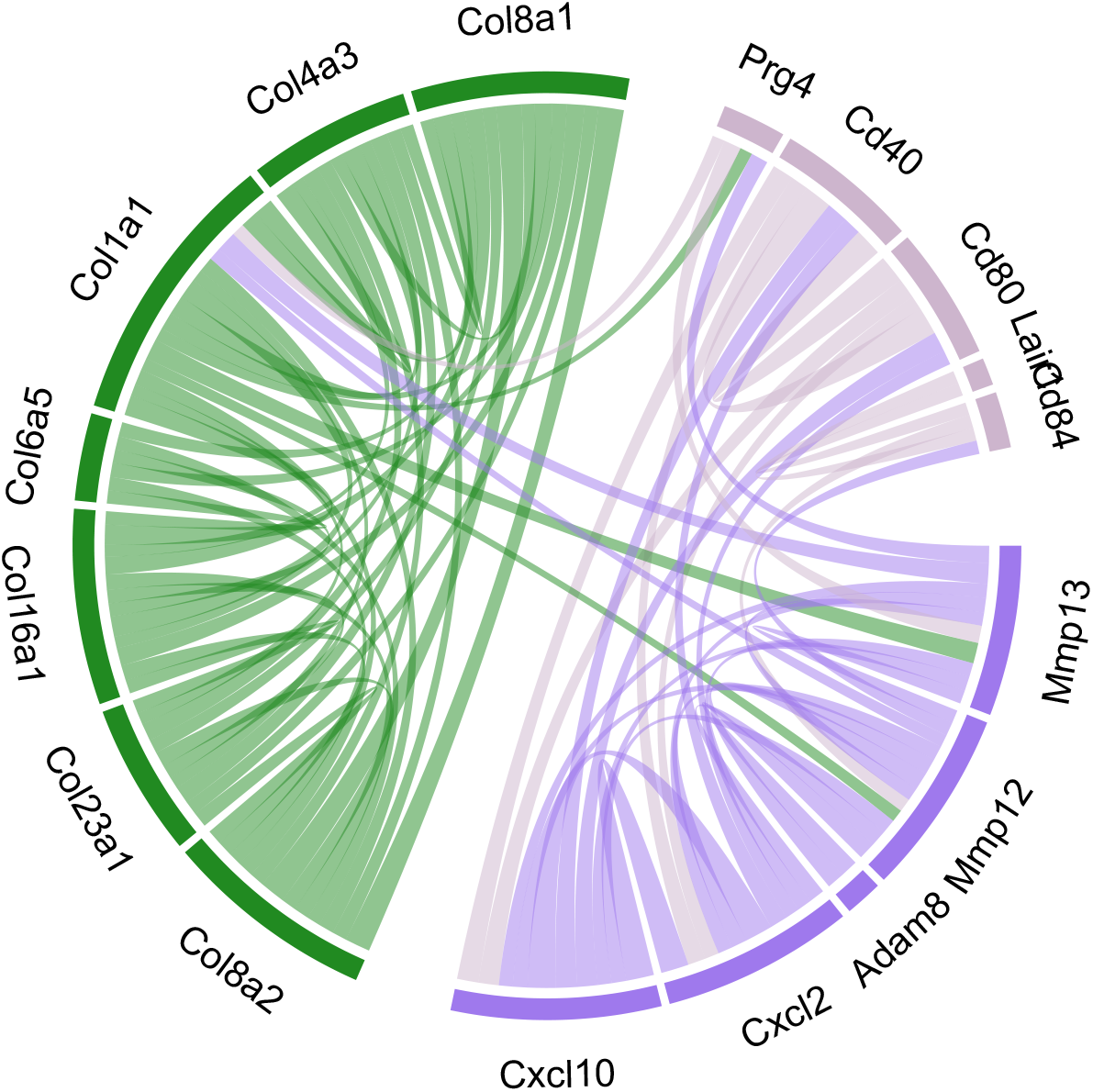

### 6.13 Ternary plot: M1 vs M2: Coates macrophage gene set

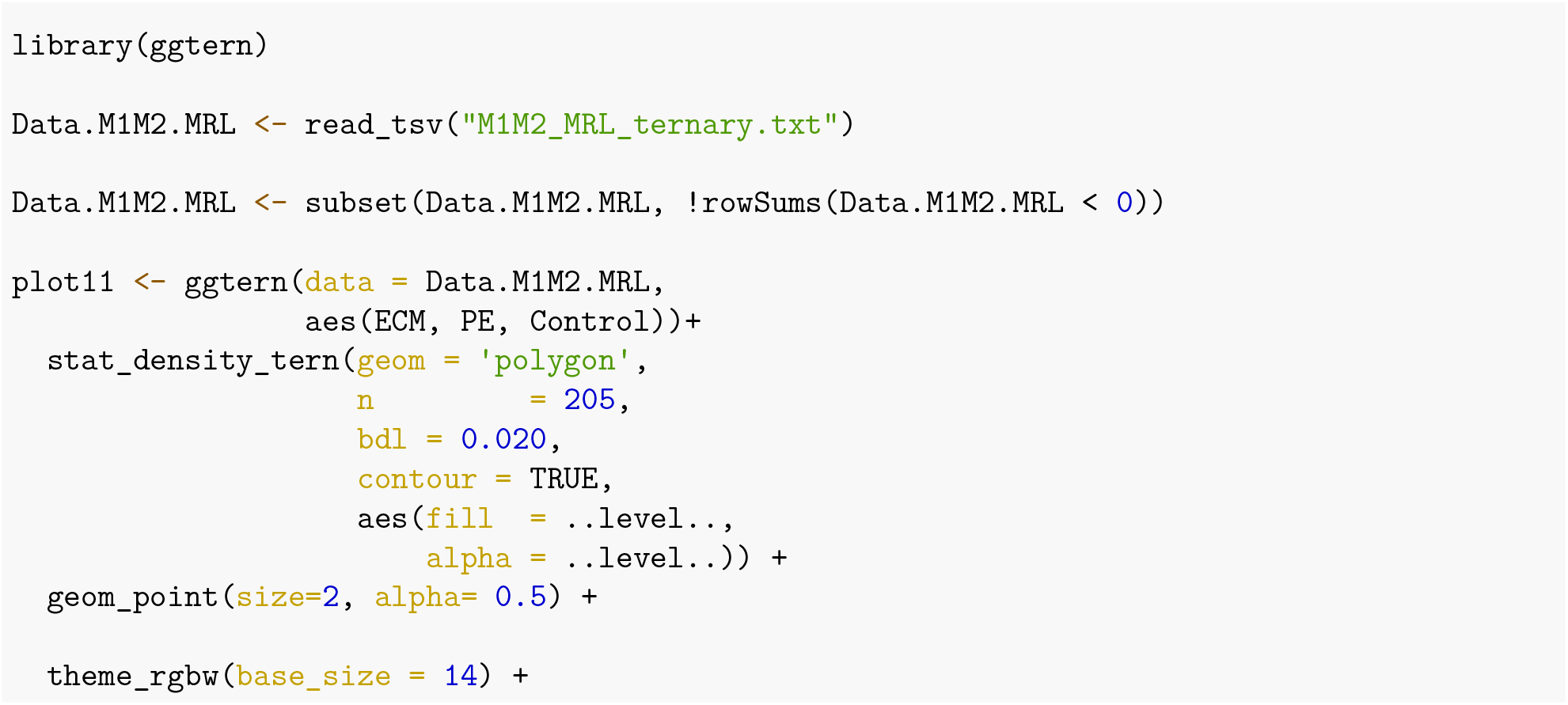

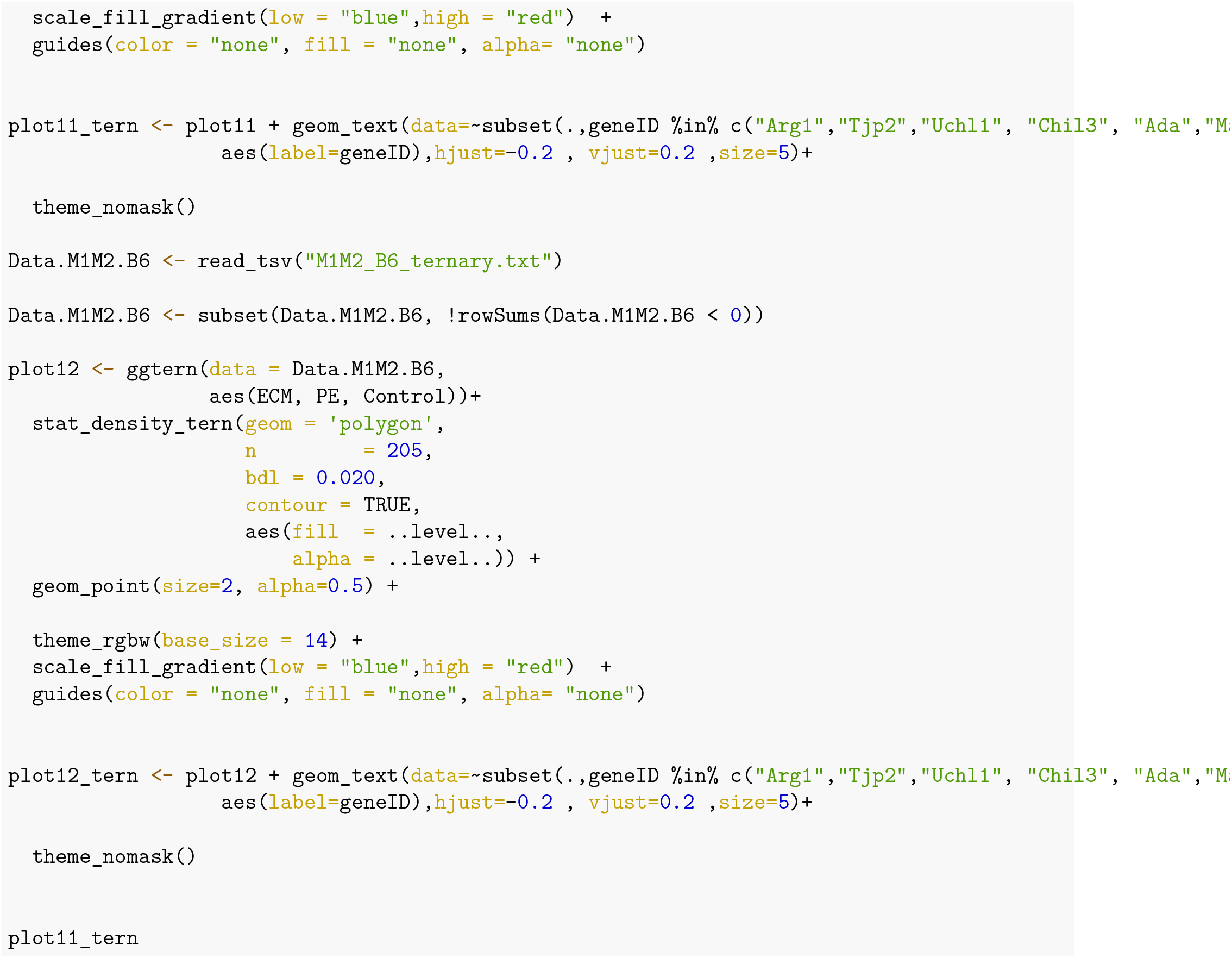

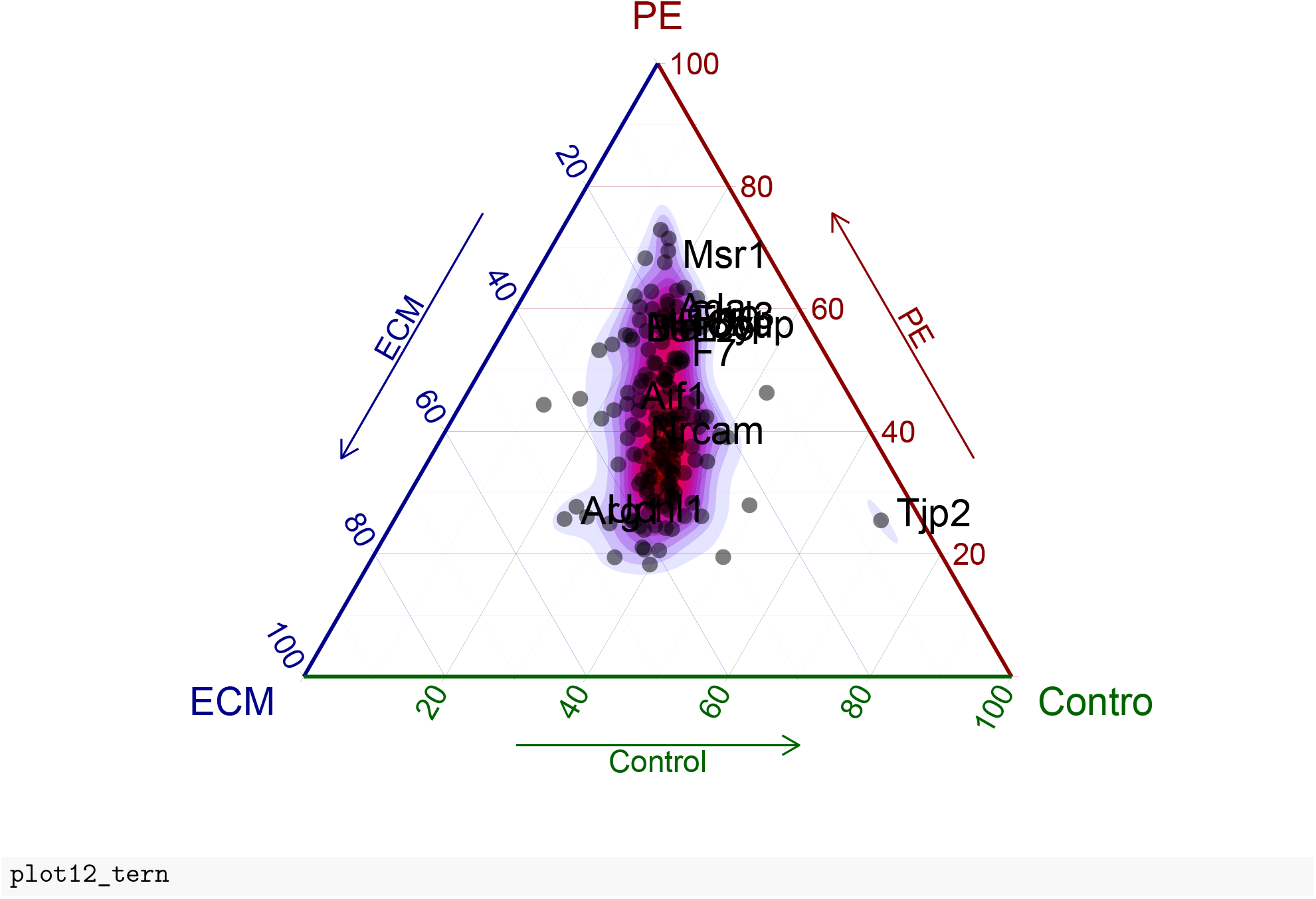

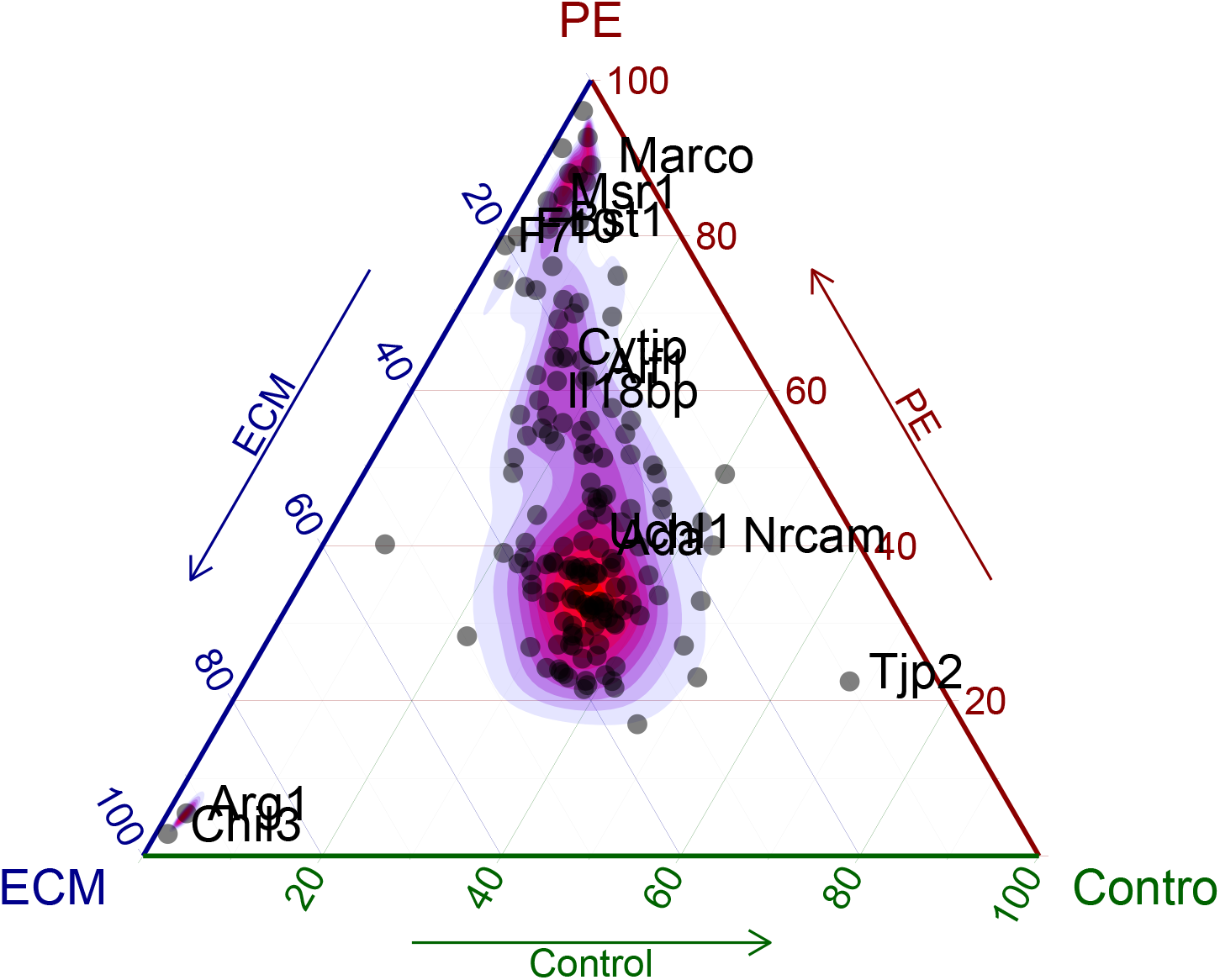

### 6.14 Ternary plot: Neutrophil gene set

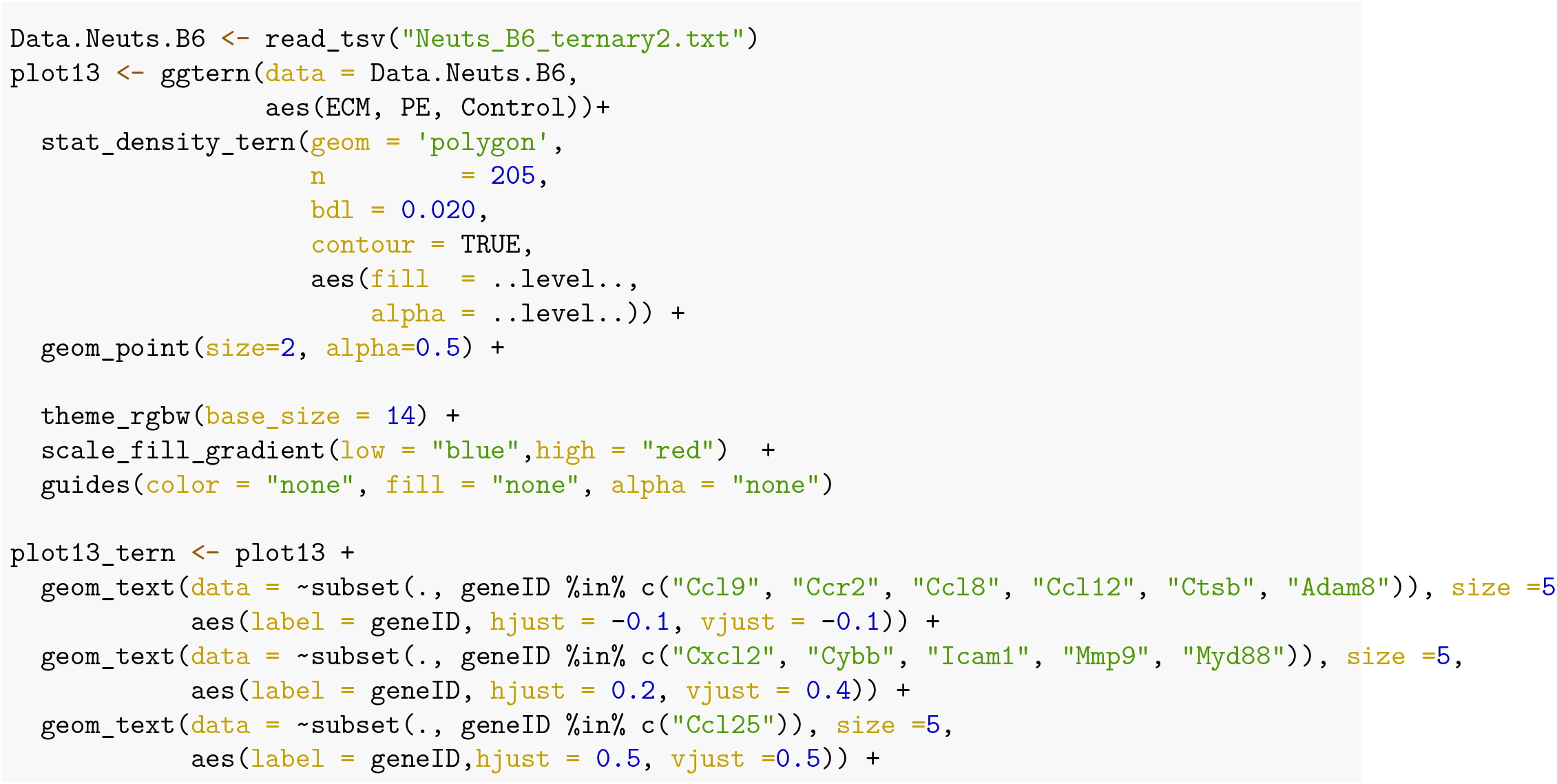

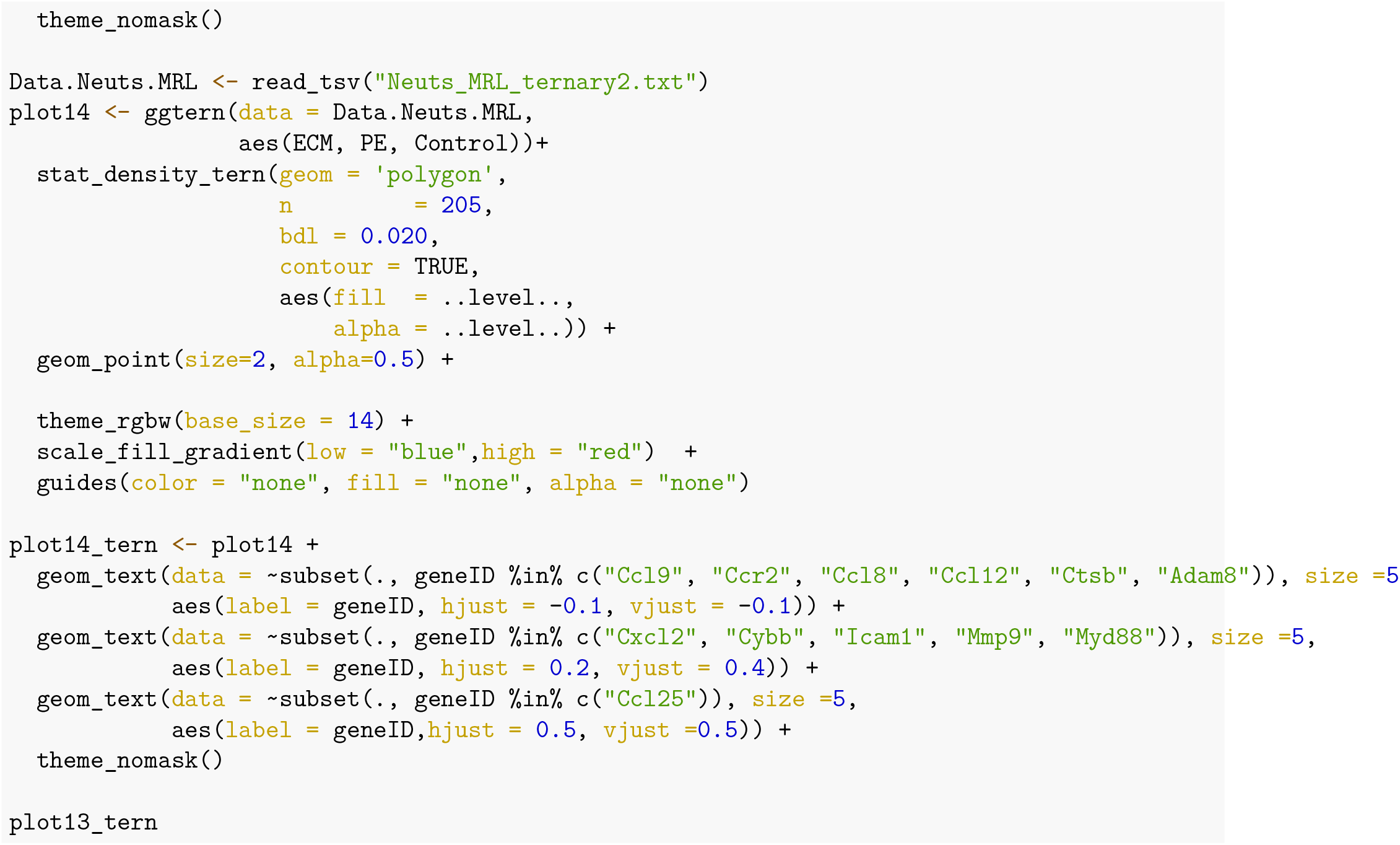

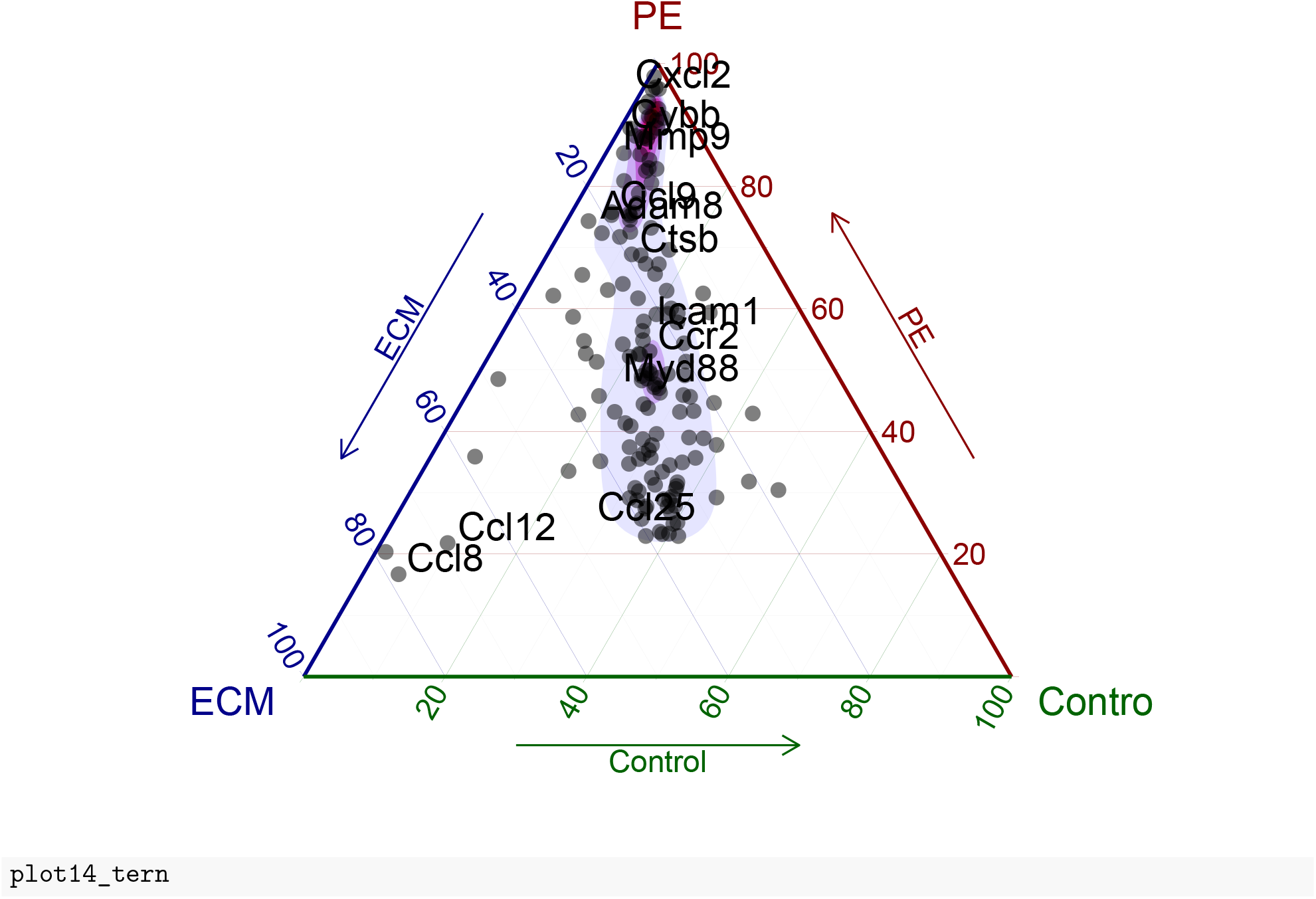

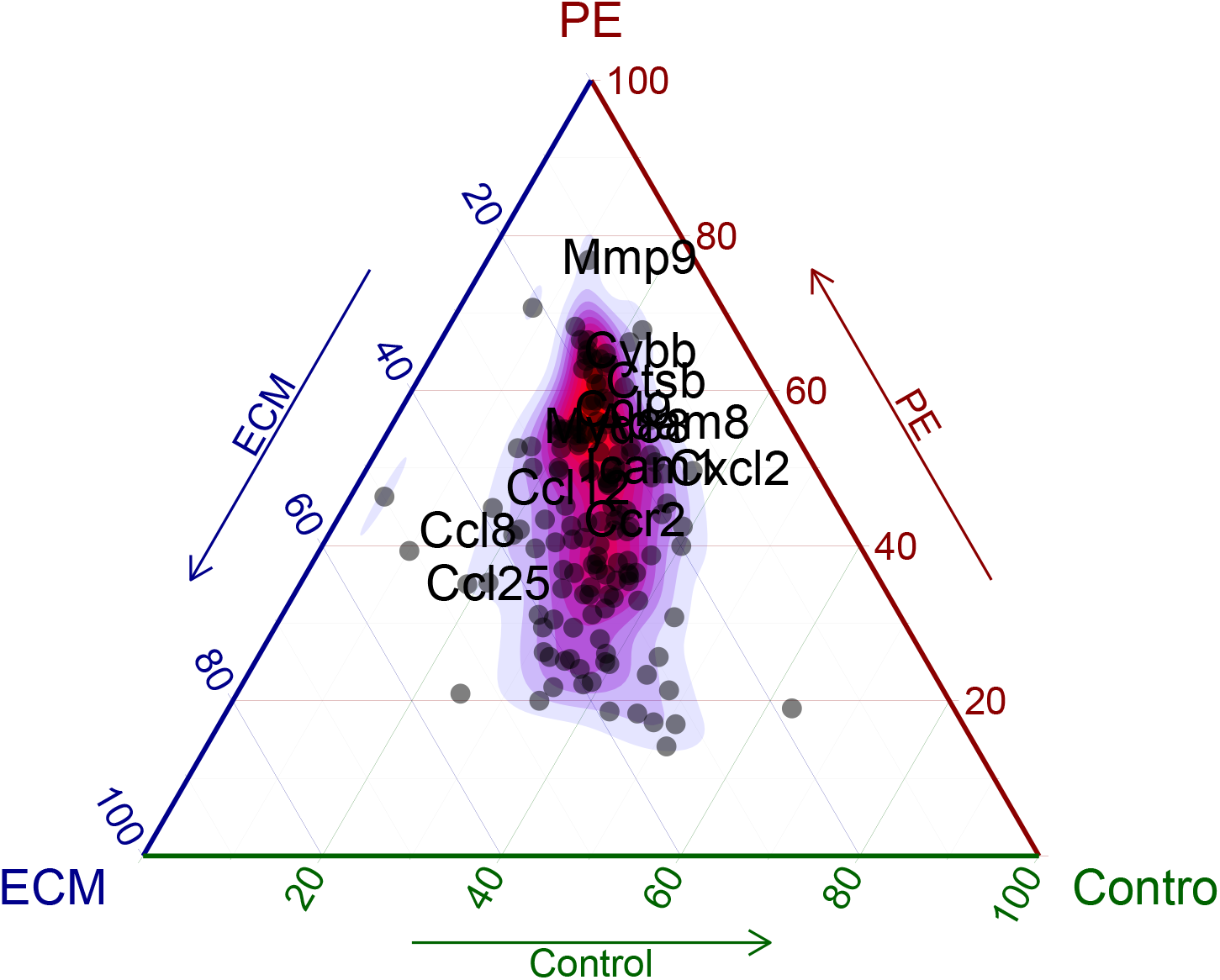

### 6.15 Ternary plot: Eosinophils gene set

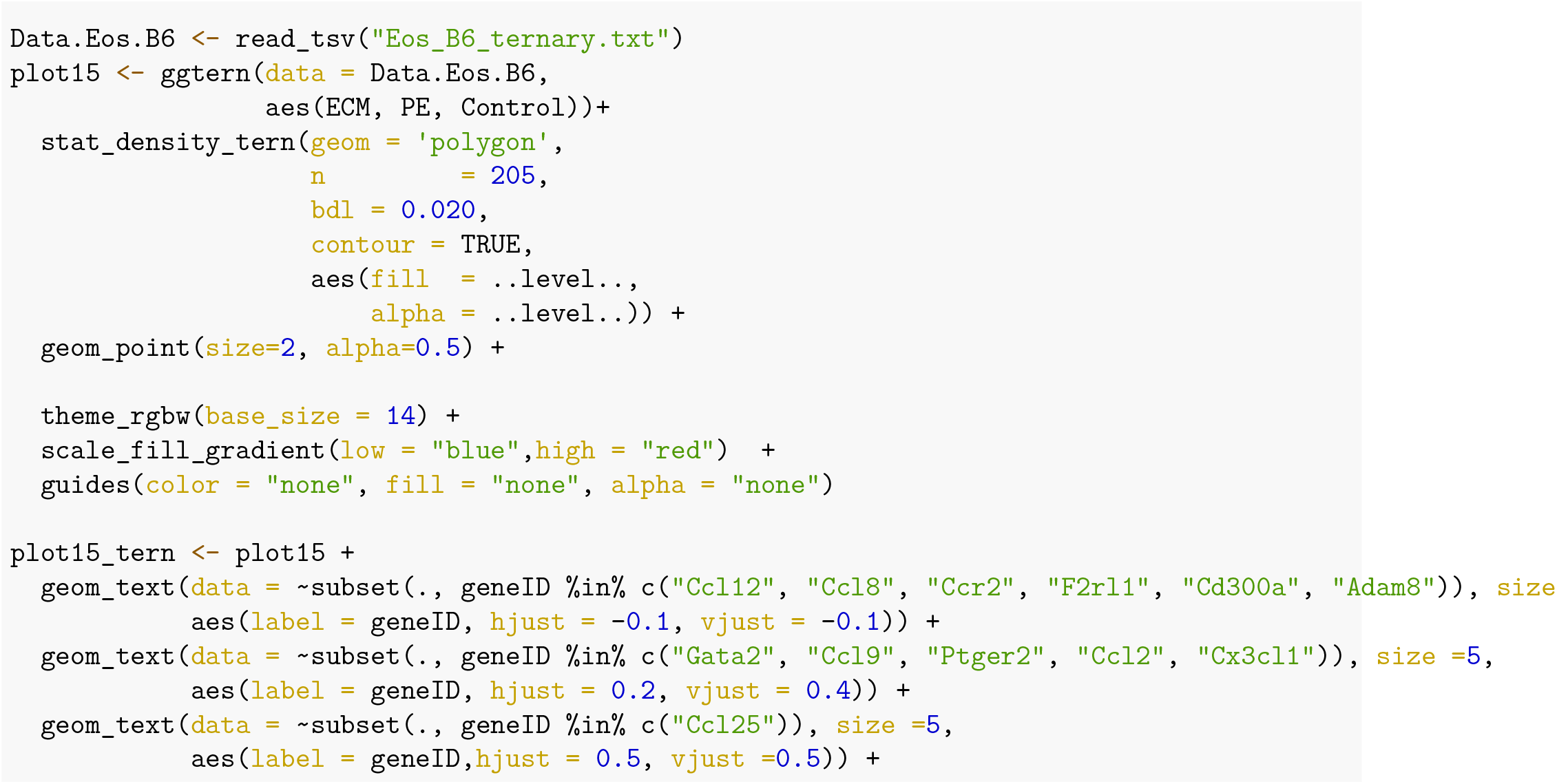

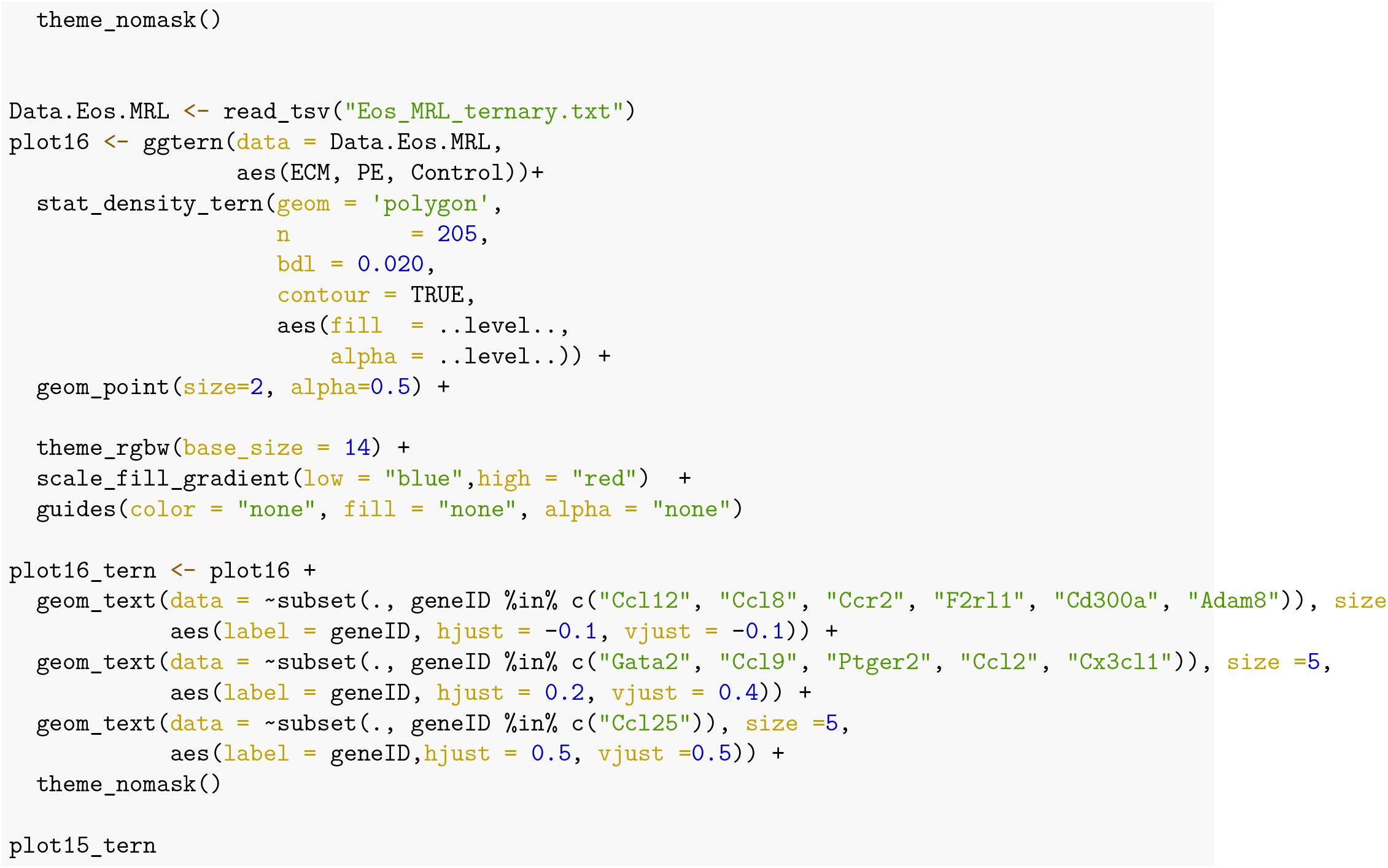

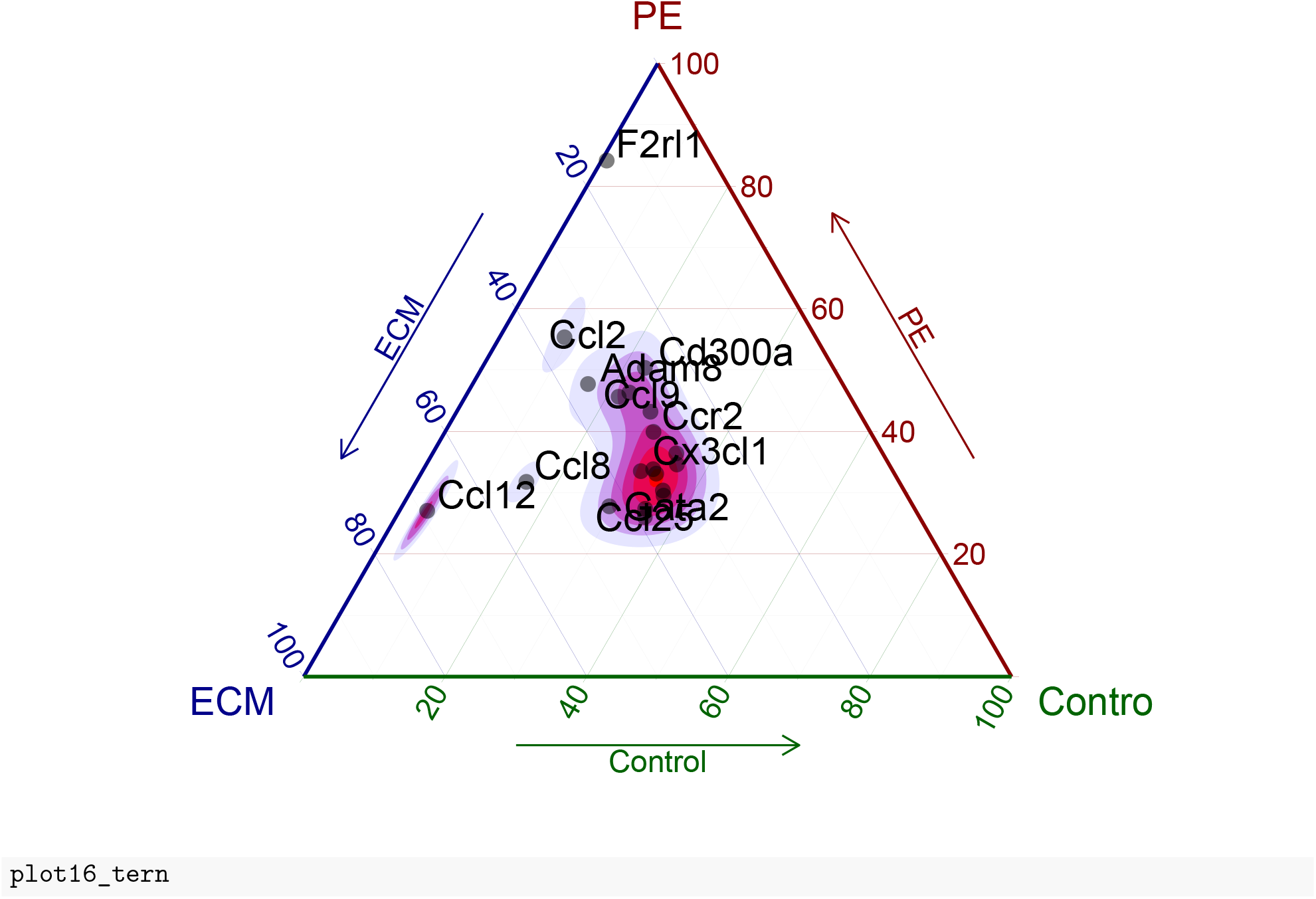

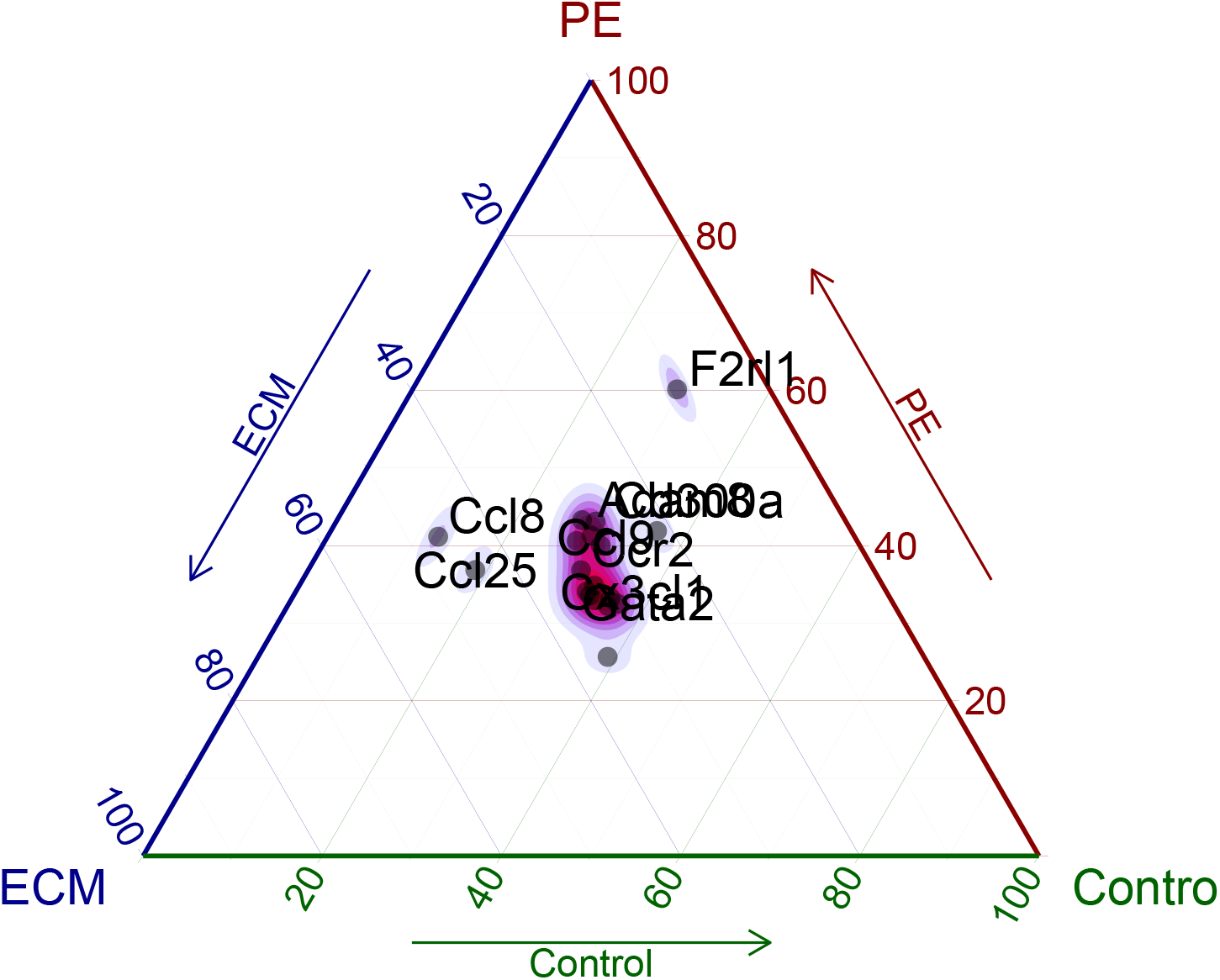

## Notes

### Competing Interest Statement

The authors have declared no competing interest.

